# Mixtures of stably coexisting species overyield

**DOI:** 10.64898/2026.09.17.752039

**Authors:** Mario Desallais, Michel Loreau, Jean-François Arnoldi, Rudolf P. Rohr

**Affiliations:** Department of Biology—Ecology and Evolution, University of Fribourg; Ch. du Musée 10, CH-1700 Fribourg, Switzerland; Institute of Ecology, College of Urban and Environmental Sciences, Peking University; Beijing 100871, China; Theoretical and Experimental Ecology Station, CNRS; 2 route du CNRS, 09200 Moulis, France

## Abstract

Diverse plant communities usually produce more biomass than monocultures^1–3^, an advantage that strengthens over time^1,4^ and is attributed mainly to the complementarity effect^5^—the extra biomass produced, relative to monocultures, when species exploit different resources, share fewer enemies or facilitate one another. Yet whether this advantage is a mere tendency of diverse communities, or something that every persistent community must have, has remained unknown. Here we show that stable coexistence requires a positive complementarity effect, with vanishingly rare exceptions, and that underyielding communities are significantly more likely to lose a species within two years or sampling steps, across four long-term biodiversity experiments, with odds of losing a species nearly three times higher than in other communities. In the twenty-five years since the complementarity effect was defined^5^, the link between biodiversity effects and species coexistence had only been established theoretically for two species in competition^6^ or observed as a correlation^7^; our theoretical result makes it a general property of multispecies communities, in which the complementarity effect scales with two established measures of how securely species coexist—local and structural stability^8–10^. Our analyses are, to our knowledge, the first to test a coexistence prediction—impending species loss—prospectively in biodiversity experiments, from a functioning measure alone, turning a measure of what diversity does into a diagnostic of whether diversity can persist. Stable coexistence and higher productivity are thus intrinsically linked: biodiversity does not merely benefit ecosystem production on average; biodiversity that endures cannot but benefit it through complementarity. We anticipate that this diagnostic will turn existing biodiversity experiments, sown grasslands and intercrops into early-warning systems for species loss, since it requires only the biomass of each species in mixture and in monoculture—data these systems already collect, and that agronomists already combine into the land equivalent ratio—and that it will extend to forests, through the growing networks of tree diversity experiments and mixed-species plantations where the same measurements are made.

---

Ecology seeks to understand why so many species coexist and how this diversity governs the functioning of ecosystems. There is broad agreement on the latter issue: more diverse communities usually produce more biomass^1–3^ and this advantage strengthens as communities assemble over time^1,4^. This pattern has been mainly explained by the *complementarity effect*^5^, the extent to which species produce more biomass than expected when grown together, through differences in the resources they exploit or the natural enemies they share, or by facilitating each other. Consequently, these species perform better together than predicted from their performance in monoculture^11,12^. But this consensus leaves a deeper question open: is this advantage of diversity something that merely tends to occur on average, or something that every persistent community must have?

Answering this question requires bringing together two branches of ecology that have long been pursued independently. Biodiversity–ecosystem functioning (BEF) research asks what diversity does—how much biomass a community produces and how stable this production is—while coexistence theory asks whether species can persist together at all^13^. The two have rarely met: whether a strong complementarity effect tells us anything about whether a community can persist has been addressed only in special cases^6,14–19^, such as in communities with two species^6^ or with particular structures^18,19^, and the generality of any such link has been questioned^20,21^. Recent analyses of long-term grassland experiments report that communities better able to coexist tend to show stronger complementarity^7^, but correlations alone cannot establish that the link is necessary.

Here we demonstrate that stable coexistence and positive effects of biodiversity on ecosystem functioning are bound together, and that the advantage of diversity is not a tendency but a requirement. Specifically, a negative complementarity effect arises, with vanishingly rare exceptions, only in transient communities whose equilibria are unfeasible or unstable—a warning of impending extinctions. Persisting in the long run, therefore, requires a positive complementarity effect. Importantly, the larger the complementarity effect is, the greater the community’s potential to coexist.

We establish this mathematically, building the argument from the bottom up. For two species, the complementarity effect can be considered a continuous measure of the potential for feasibility and stability, and thus of stable coexistence, extending a known qualitative result^6^. We then demonstrate that this result generalizes to species-rich communities. This link is established at community equilibrium, proving that any observation of a negative complementarity in an empirical dataset is a signal of a transient community. To predict the fate of such a transient state and confirm that it leads to at least one extinction, we extend our theory to the non-equilibrium regime. We show that a community’s total yield settles far faster than its composition shifts and that a negative complementarity effect can occur while a community is still changing—as a predictor of future extinction. Finally, we test the resulting prediction—that negative complementarity precedes species loss—in four long-term biodiversity experiments, from grasslands to protist microcosms. Our results demonstrate that negative complementarity foreshadows extinction, whereas higher levels of complementarity are associated with lower odds of extinction.

## The complementarity effect measures how securely two species coexist

Overyielding can be captured by a single, tractable quantity, the relative yield total RYT: the sum of the ratios of each species’ biomass in mixture to its biomass in monoculture^11,22,23^, itself a key component of the complementarity effect. Indeed, the complementarity effect of a community of *S* species with an average carrying capacity (i.e., monoculture production) 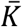 relates to the relative yield total as 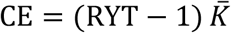 (5, 24) (Methods). That is, CE > 0 is equivalent to RYT > 1, which defines overyielding; conversely, a negative complementarity effect (CE < 0) is equivalent to underyielding (RYT < 1).

In two competing species governed by Lotka–Volterra dynamics—the classical model used in theoretical ecology—stable coexistence implies overyielding (CE > 0, i.e., RYT > 1) (6), and, consequently, a mixture that underyields cannot stably coexist. This classic result fixes only the *sign* of the complementarity effect, not its magnitude. Yet the same two-species geometry (Fig. 1A) contains a quantitative law: the complementarity effect relates continuously to two established measures of how securely the species coexist—one indicating whether the species will recover after a disturbance (*λ*_max_), the other how wide a range of environments lets both species exist (Ω).

**Fig 1.**
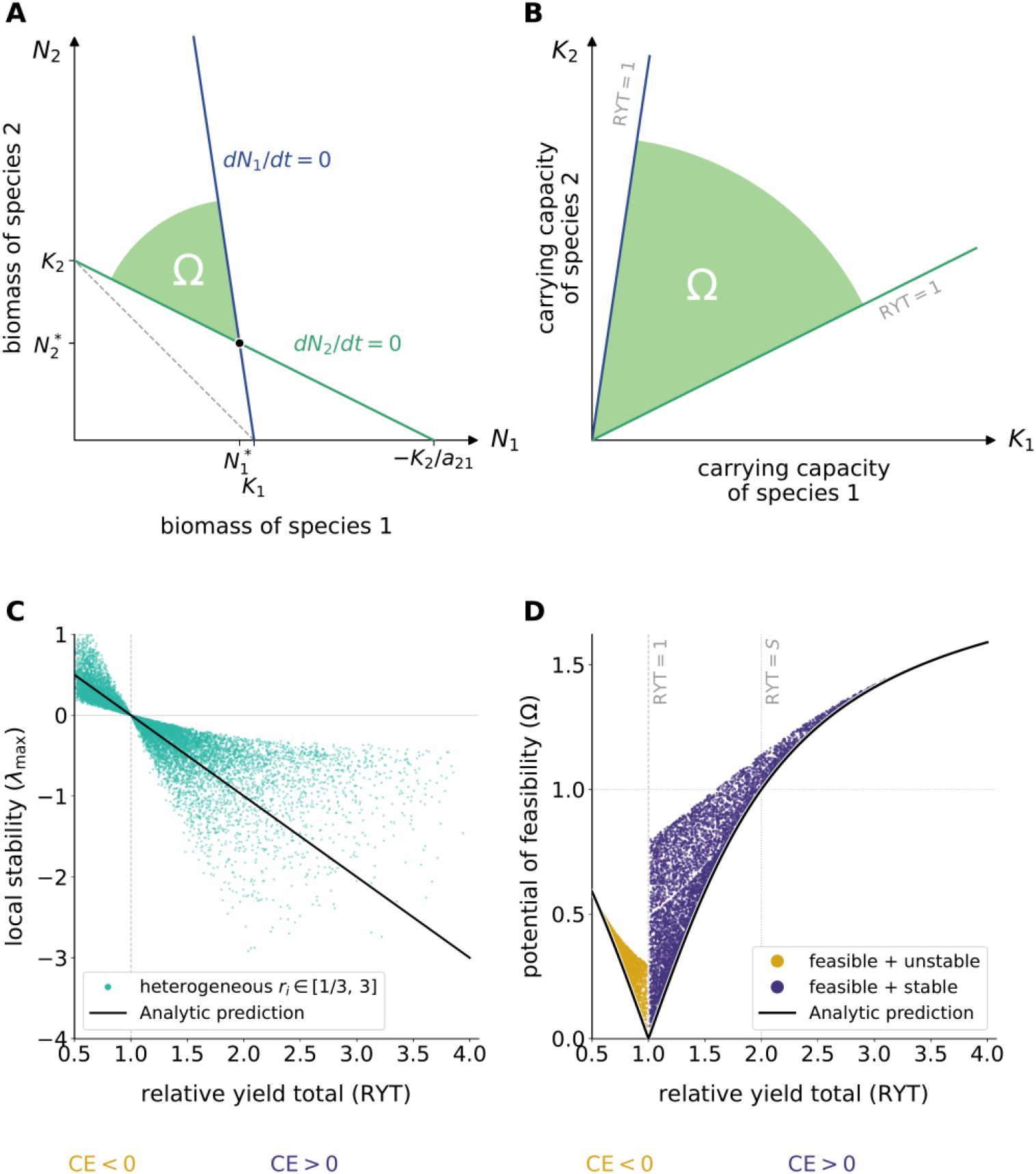
In two interacting species, for competition and facilitation, RYT relates quantitatively to stability and to the range of viable environments. **(A)** Zero-growth lines (isoclines) in the (*N*_1_, *N*_2_) plane cross at the equilibrium (black dot). The shaded green region is the cone of viable conditions bounded by these lines, whose normalized angle defines Ω. The dashed line marks RYT = 1. **(B)** The same cone in the (*K*_1_, *K*_2_) plane of carrying capacities: every vector inside it gives a community in which both species persist. RYT = 1 is the cone’s edge, and RYT is largest deep inside the cone, where *λ*_max_ is most negative. **(C)** *λ*_max_ as a function of RYT across two-species systems with a wide range of growth rates (*r*_*i*_ ∈ [1/3,3]). Points concentrate along the black line (the unit-rate case *r*_*i*_ = 1, SI S2.1), and across these sampled rates the line marks the average trend. **(D)** Ω as a function of RYT; within the feasible–stable region, Ω varies continuously and monotonically with RYT (SI S4.1). The black line is the analytical relation of Eq. 2, obtained in the case with equal interaction and carrying capacities (*a*_12_ = *a*_21_, *K*_1_ = *K*_2_).

Formally, *λ*_max_ determines whether a community is locally stable. If negative, the system will return to equilibrium following a slight disturbance; if not, the perturbation will be amplified. The magnitude of |*λ*_max_| measures the distance to the stability boundary, i.e., how far the community sits from switching from stable to unstable^8^. Among two-species competitive communities with heterogeneous growth rates, *λ*_max_ follows RYT closely along a single line (Fig. 1C):

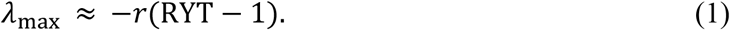

The relation is exact when growth rates are equal (SI S2.1), and the line marks the average trend across the heterogeneous rates of Fig. 1C. The earlier qualitative equivalence^6^ is recovered as the sign of RYT ™ 1. Moreover, the community moves away from local instability as RYT moves away from 1 toward higher values. This adds a quantitative dimension to the relationship between RYT and stability.

The second measure, the feasibility domain size Ω, captures how wide a range of environments a community can tolerate while retaining the existence of a feasible equilibrium, i.e., an equilibrium at which every species is present^9,10^: the larger it is, the more robustly the species coexist. Formally, it is given by the opening of the feasibility domain, which is the cone of carrying capacities for which every species holds a positive equilibrium abundance—that is, the community is feasible. In two species this cone is planar and Ω reduces to its normalized opening angle (Fig. 1B). The normalization is chosen such that if the cone of feasibility equals the (*K*_1_ > 0)–(*K*_2_ > 0) orthant, that is, if every combination of positive carrying capacities leads to feasibility, then Ω = 1, i.e., the full feasibility potential. Importantly, the cone of feasibility can also be read directly in the biomass (*N*_1_, *N*_2_) plane (Fig. 1A; SI S2.2). The link between RYT and feasibility is thus both algebraic and geometric (Fig. 1D). For RYT > 1, the cone widens continuously and monotonically with RYT and as a first-order approximation (equal interspecific interactions; derived in SI S2.3), the cone angle relates to the maximum relative yield total RYT_max_ through

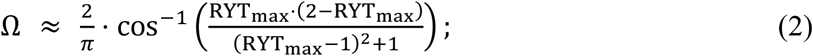

across heterogeneous parameters, this curve bounds the relation from below (Fig. 1D).

In two species, then, the complementarity effect is more than a yes/no signal of coexistence: it is a quantitative proxy for two established conditions of coexistence—local stability and feasibility potential. The larger a positive complementarity effect, the larger the two species’ potential for stable coexistence. We now extend this link to any number of species.

### Stable coexistence leads to overyielding in species-rich communities

The link between overyielding and coexistence in two-species communities rests on a geometry that does not obviously generalize to more diverse communities. With three or more species, isoclines become hyperplanes, the feasibility cone becomes a higher-dimensional object, and pairwise intuition breaks down. Yet one exact statement survives this transition. Assuming equal growth rates (*r*_1_ = *r*_2_ = *⋯* = *r*_*S*_ = *r*), the relative yield total relates to the sum of the eigenvalues as 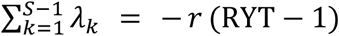. As local stability requires every eigenvalue to have a negative real part, hence their sum (SI S3.1), RYT must be greater than 1, i.e., a stable feasible equilibrium must overyield. If growth rates are heterogeneous, it is still possible to relate stable coexistence and RYT, at the price of a bound on how strongly any pair of species competes:

**Theorem 1**. In a generalized Lotka–Volterra community in which every species can persist on its own (*K*_*i*_ > 0) and no two species compete with each other, in total, more strongly than they each limit themselves (*a*_*ij*_ + *a*_*ji*_ > *a*_*ii*_ + *a*_*jj*_ = −2), any equilibrium at which all species are present— a feasible equilibrium—overyields: RYT > 1, and the complementarity effect is positive.

The model and its sign conventions are specified in Methods^25^. Here *a*_*ii*_ = −1 is the self-regulation term and *a*_*ij*_ the effect of one species on another (negative for competition, positive for facilitation), so the lower bound *a*_*ij*_ + *a*_*ji*_ > −2 caps how strongly any two species can mutually compete. We give the formal statement and full derivation in SI S3.2.

Coexistence requires both feasibility and stability, and the theorem only concerns feasibility. Yet the same bound on mutual competition also governs stability. Communities that break the bound are seldom stable: competition that is too strong within a pair tends to amplify disturbances across the whole community, tipping it into local instability (SI S3.3.1). As a result, violation of the interaction bound makes instability overwhelming (Fig. 2).

**Fig 2.**
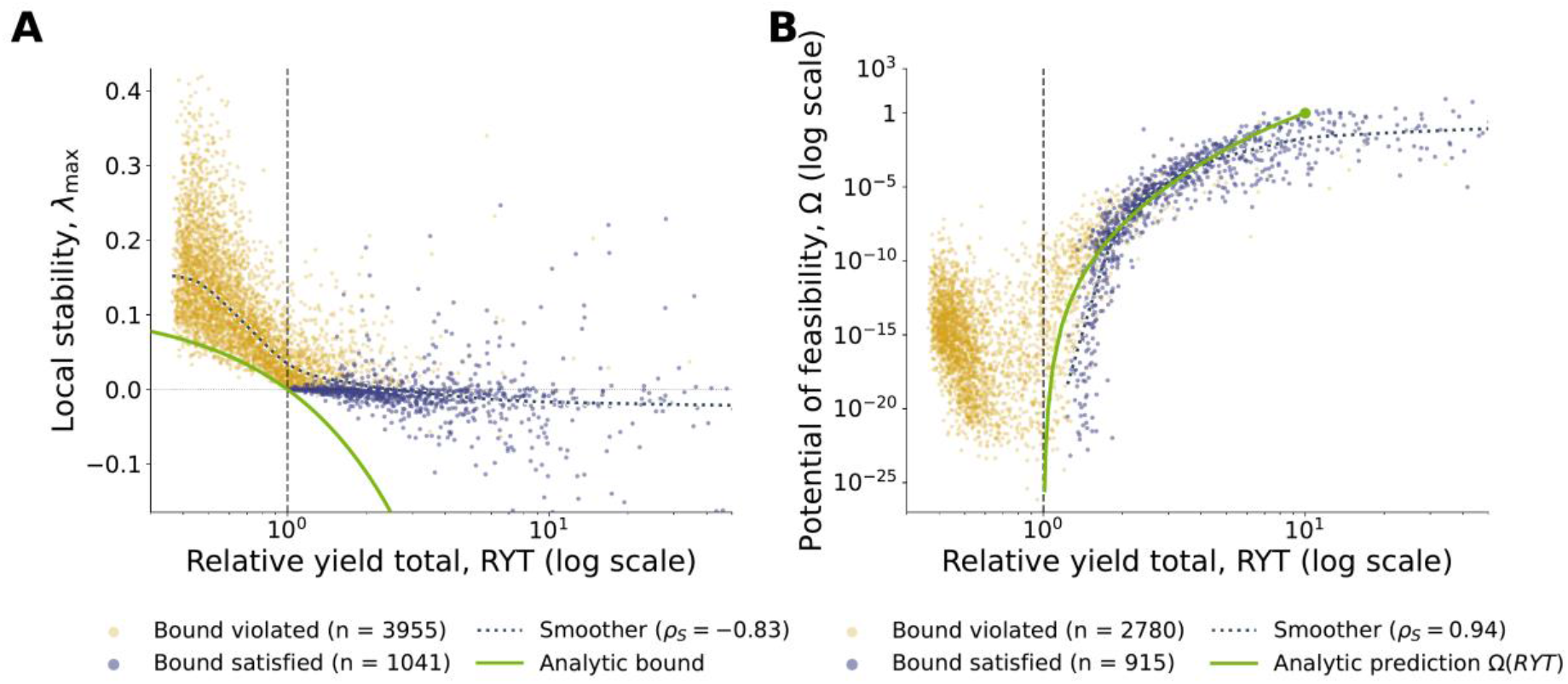
Numerical simulations confirm that the pairwise interaction bound jointly controls overyielding, local stability, and the feasibility domain. Joint distributions of RYT (log scale) with the feasibility-domain size Ω and the leading Jacobian eigenvalue *λ*_max_ across feasible communities at *S* = 10, partitioned by whether the pairwise interaction bound *a*_*ij*_ + *a*_*ji*_ > −2 is satisfied (purple) or violated (yellow). Robustness across *S* ∈ {3,5,10,15} and four interaction structures is documented in SI S4.1 (Figs. S4 and S5). **(A)** *λ*_max_ as a function of RYT. The leading Jacobian eigenvalue decreases continuously with RYT across the full cloud (Spearman *ρ*_*S*_ = −0.83, dotted smoother); communities satisfying the bound are concentrated in the locally stable region (*λ*_max_ < 0, large RYT), while violators dominate the unstable region (*λ*_max_ > 0). The green curve is the analytical bound of Eq. S54, derived at equal growth rates (SI S3.5.1). **(B)** Ω (log scale) as a function of RYT. The bound condition forbids the half-plane RYT < 1 entirely (consistent with Theorem 1); communities satisfying it concentrate at large Ω and RYT > 1, while violators occupy a separate region with RYT that may fall below 1 (Spearman *ρ*_*S*_ = 0.94). The green curve is the analytical prediction of Eqs. S57–S59 (SI S3.5.2).

This yields a general conclusion. Either every pair of species respects the competition bound, in which case Theorem 1 lets a community underyield only by losing feasibility—losing the equilibrium itself—or the bound is violated somewhere, in which case the feasible communities involved are overwhelmingly unstable (Fig. 2; SI S3.3.1). A positive complementarity effect is therefore necessary for coexistence throughout the broad competitive regime the bound defines, and—as the simulations show and SI S3.3.1 explains analytically—for the overwhelming majority of communities beyond it. Past the bound, a feasible community may remain stable while underyielding, but only by virtue of a particular, finely tuned pattern in how fast its species grow coupled with asymmetric interactions; this is impossible when all species grow at the same rate (SI S3.3.2).

As in two species, the more a community overyields, the wider the range of environments in which all its species still stably coexist (SI S3.5). Both the analytical relations displayed in Fig. 2 recover Eqs. 1 and 2 at *S* = 2, and simulations at *S* = 10 show that the two orderings survive heterogeneous growth rates and structured interactions.

A stable underyielding equilibrium is therefore highly unlikely (Fig. S3). Any observation of RYT < 1 in a real-world experiment is therefore an indication of a transient community. However, understanding whether a transient underyielding community will head toward extinction requires extending the framework to non-equilibrium dynamics.

### Away from equilibrium, underyielding forecasts species loss

A community’s dynamics can be partitioned in two aspects: the dynamics of its RYT and the dynamics of its composition ***p***, the vector of relative shares *p*_*i*_ = *RY*_*i*_/RYT. We observe a time-scale separation between the two: RYT quickly reaches the value its current species composition ***p*** can support, following a logistic curve, while the composition ***p*** itself shifts far more slowly (SI S2.4 and S3.6).

This is readily seen in the two-species case (Fig. 3A, B): trajectories first move along near-rays through the origin—both abundances increasing in near-constant proportion, the composition ***p*** barely changing—until they approach the teal curve, where RYT temporarily settles at RYT^*^(***p***); only then do they slide along that curve, as the slower compositional dynamics take over. Numerically, this separation persists across a broad range of growth-rate heterogeneity, with heterogeneous interactions and in multispecies communities (Fig. S7). An RYT measured from a transient community, therefore, reports RYT^*^(***p***)—the settling value set by the species currently present, their relative proportions, and their interactions. This transient RYT value thus carries information about species interactions: RYT^*^(***p***) < 1 tells us that the current species composition is dominated by species that compete strongly, making the path to stable coexistence very unlikely (Fig. 3D and SI S3.6), whereas a community on its way to stable coexistence typically shows RYT^*^(***p***) > 1 throughout (Fig. 3A).

**Fig 3.**
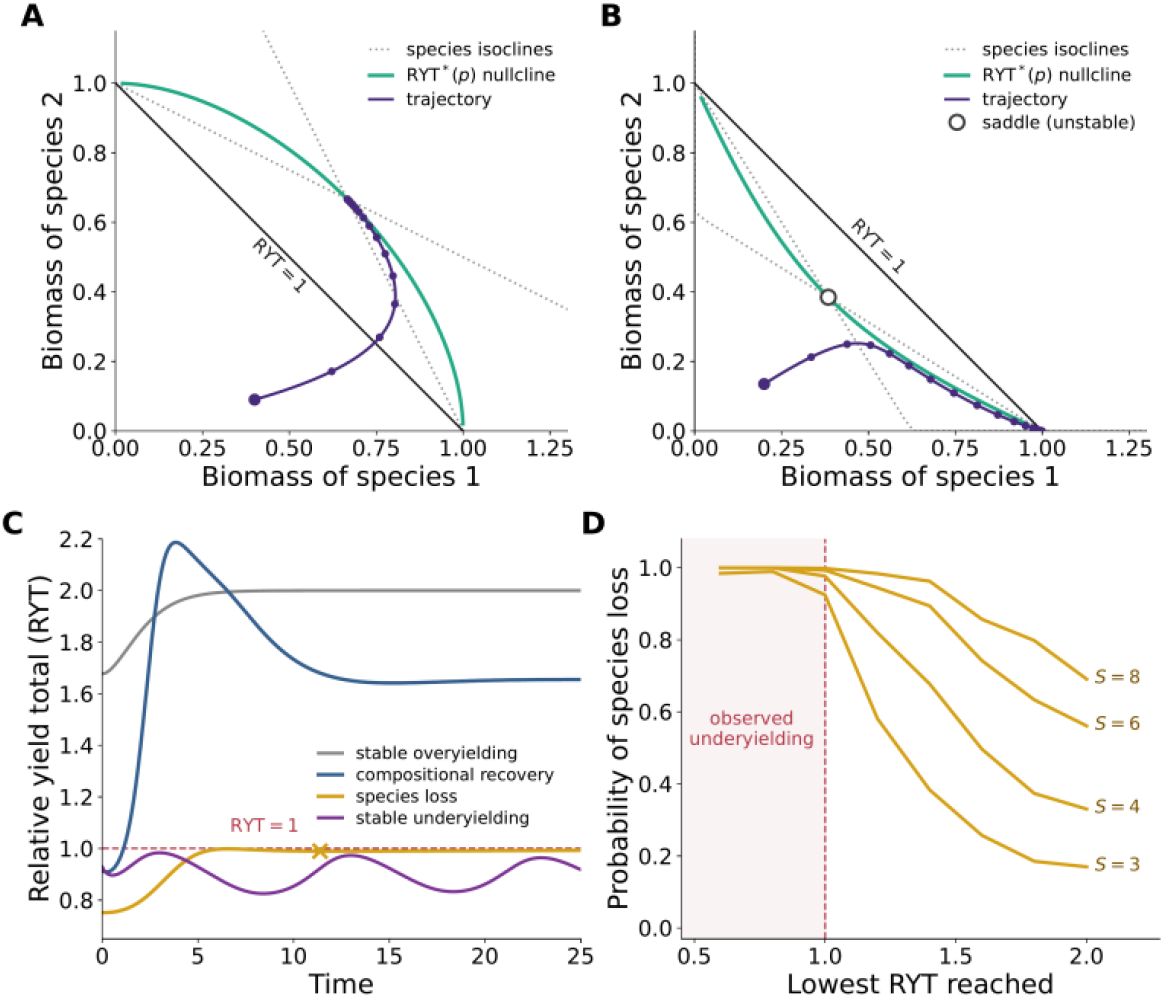
Read along a community’s trajectory, the relative yield total forecasts species loss. **(A, B)** Two-species phase planes with equal growth rates. Gray dotted lines, isoclines; black diagonal, RYT = 1; teal curve, RYT^*^(***p***), the total the community quickly settles to for a given mixture ***p***. Trajectories move fast onto this curve, then drift slowly along it (dots mark equal time intervals). **(A)** A pair within the bound of Theorem 1: the teal curve lies wholly outside the RYT = 1 line, and both species coexist and overyield. **(B)** A pair beyond the bound: the teal curve lies wholly inside the line, and the community reaches an edge of the plane, where one species is lost; open circle, unstable balance point. **(C)** Relative yield total through time for four communities (dashed line, RYT = 1): stable overyielding, all species kept (gray); underyielding, then loss of a species (amber, ×); compositional recovery, dipping below 1 with every species kept (blue); stable underyielding (purple). **(D)** Probability of losing at least one species against the lowest RYT reached, for *S* = 3, 4, 6 and 8 (sampling in SI S4.2). Once a community underyields (shaded), loss is all but certain at every richness; above 1, the chance of loss falls as RYT rises, most steeply when species are few. At *S* = 4, 4,410 communities (23%) underyielded; 4,398 of them (99.7%) lost a species, and only seven (0.16%) recovered with every species kept. Of the 2,601 communities whose total fell below 0.9, not one recovered intact.

An observed underyielding thus forecasts species loss. Only two configurations escape this fate. In the first, which we call *compositional recovery* (blue curve, Fig. 3C), underyielding is transient yet does not lead to extinction. Pairs competing beyond the interaction bound dominate the community early and make RYT^*^(***p***) < 1, but the community returns to overyielding as weakly competing species regain relative abundance. In the second, a community underyields persistently yet remains stable (purple curve, Fig. 3C). This is the case of *stable underyielding* described in the previous section. The mechanism is the following: two species that compete too strongly to coexist on their own can persist thanks to a third, faster-growing species. Because the fast species settles almost at once, it forms a steady background through which the slower pair interacts indirectly. Their effective competition is thus softened into something both survive (SI S3.3.2), leading to a persistently underyielding state approached through slow, weakly damped oscillations (Fig. 3C). Simulations show that compositional recovery and stable underyielding are highly unlikely: a transient RYT < 1 is mostly followed by an extinction (Fig. 3D and Fig. S3). Therefore, an experimentally measured underyielding value, though transient, is informative enough to predict an impending extinction, a prediction we test in the following section.

### Underyielding precedes species loss in long-term experiments

The theorem, its non-equilibrium extension and the quantitative relationships between complementarity and stable coexistence make a prediction we can test in the field: a community that underyields—so that its complementarity effect is negative—should be on its way to losing species. We tested this in four long-term biodiversity experiments that together span three orders of magnitude in generation time: protist microcosms^26^, two grassland experiments^27–29^, and a multi-site agricultural experiment^30,31^. At each observation of each experimental unit—a field plot, or a microcosm in the protist study—we extracted: i) whether the community underyielded, and ii) whether it then lost a species (Methods). We asked whether loss follows within two years or sampling steps, estimating each experiment’s odds ratio by penalized logistic regression and combining the four experiments into a single pooled estimate (Methods); cluster-robust estimates, which allow repeated observations of the same unit to be correlated, lead to the same pooled conclusions (SI S7.5).

Underyielding raised the risk of subsequent species loss. Two observations ahead, the effect is clear: across the four experiments combined, underyielding multiplied the odds of losing a species by about 2.6-fold (pooled odds ratio 2.61, 95% confidence interval [1.40,4.87]; Fig. 4A). The odds ratio here is the factor by which underyielding multiplies the odds of a subsequent loss, a value of one meaning no association. One and three observations ahead, the association points the same way (pooled odds ratios 2.29 and 2.31; SI Tables S4 and S2). The effect is largest in the most controlled system, the protist microcosms, where the odds of loss were multiplied close to six-fold; it is also clear in the multi-site agricultural experiment, and the two grassland experiments reach the same conclusion.

**Fig 4.**
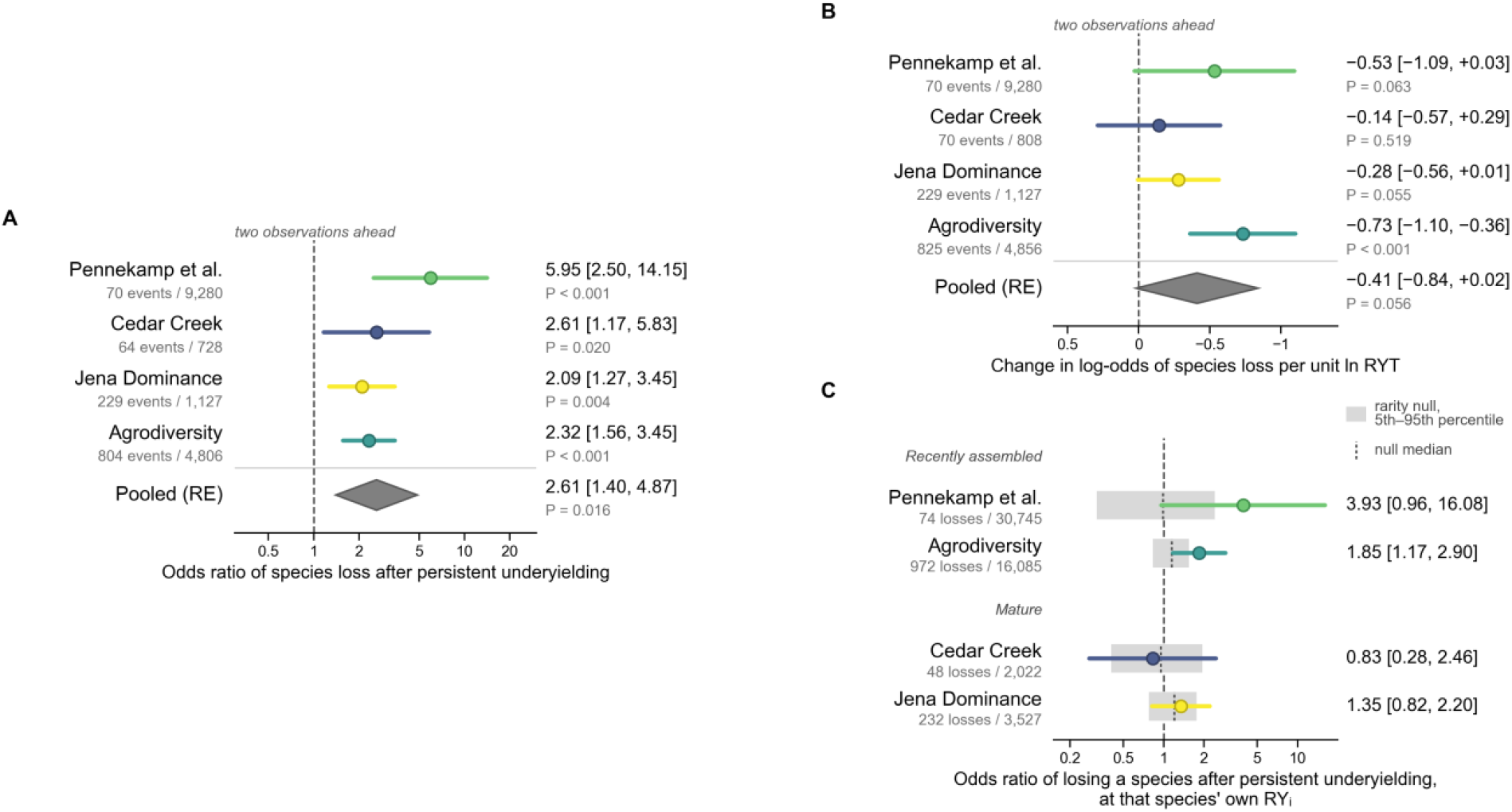
A negative complementarity effect forecasts species loss across four long-term biodiversity experiments. All panels refer to the next two observations of each plot or microcosm. **(A, B)** Community-level tests: the event is the persistent loss of at least one reference-community species; labels give the number of baselines followed by such a loss and the total number of baselines (*N* events / *n*). B can include more baselines, as the continuous test does not need the previous observation. Gray diamonds, pooled random-effects estimates (SI Table S4). **(A)** Threshold test: odds ratio of species loss after persistent underyielding (RYT < 1 at two consecutive observations), relative to its absence. **(B)** Continuous test: change in the log-odds of species loss per unit of lnRYT, on a reversed axis so that the predicted direction points right, as in A. **(C)** Test against rarity: odds ratio of losing a given species after persistent underyielding, at that species’ own relative yield RY_*i*_. Here the unit is a reference-community species present at a baseline: labels give the number of species-baselines followed by the loss of that species and the total number of species-baselines (*N* losses / *n*), which therefore differ from those of A and B. Gray bands span the 5th to 95th percentiles of the same estimate under a null model in which losses depend only on the species’ rarity; dotted lines, null median. Estimates are covariate-adjusted (Methods); dashed lines, no association; bars, 95% confidence intervals; *P* values are two-sided.

The same prediction reads as a gradient, not only a threshold: does the risk of loss fall as relative yield rises? Fig. 4B shows that it does, dataset by dataset—the slope 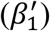 is negative, meaning a higher relative yield total lowers the odds of loss, in all four experiments, steeply in the agricultural experiment (*β* = −0.73; *P* < 0.001) and the protist microcosms, weaker in the filtered grasslands. Pooled, the gradient points the predicted way, but the four systems differ too much in steepness for a single slope to reach significance (*β* = −0.41 [−0.84, +0.02]).

A species that is already rare because of an ongoing extinction automatically lowers the relative yield total. We therefore asked whether underyielding forecasts the loss of a given species beyond what that species’ own relative yield predicts and compared the answer with a null model in which losses depend on rarity alone (Methods). In the two recently assembled systems, the protist microcosms and the agricultural experiment, it does: the estimate lies beyond the 95th percentile of the rarity-only null. In the two mature grasslands, it does not (Fig. 4C).

### Community assembly, a filter that retains only beneficial diversity

A community whose complementarity is negative is not on the path to a feasible and stable coexistence: either no feasible equilibrium exists, or—when competition between some species grows too strong—the equilibrium is almost always unstable (SI S3). Either way, the community is on a trajectory toward losing species. Community assembly therefore acts as a filter: configurations that underyield are removed by species extinctions. The configurations that remain are those that overyield, offering an explanation for the observed tendency for complementarity to increase over time in grassland experiments^1,4,32,33^.

This filter makes a prediction our datasets bear out. Where communities have not yet been sorted by long coexistence—assembled experimentally or freshly combined—underyielding should be informative; where assembly has already removed the incompatible combinations^34^, it should add little to what the rarity of the species lost already predicts. Our results show the strongest signal in the protist microcosms, where feasibility and stability can be broken by experimental design^26^, and a weaker one in the long-running grasslands (Fig. 4C), where mature communities have largely been filtered toward coexistence. The weaker signal there is not a failure of the prediction; it is what the prediction implies once the filter has done its work. Across the largest compilation of measured plant–plant interactions^35^, the vast majority of species pairs satisfy the competition bound the theorem requires—88% of all pairs, and 80% of the strictly competitive ones (SI S5)— so real communities seem to sit overwhelmingly in the regime where a negative complementarity marks a community that does not converge to stable coexistence.

### Mechanisms that can decouple productivity and stability

Our results link two properties that need not, in general, coincide: producing more biomass in a mixture than in a monoculture (a positive complementarity effect) and returning to equilibrium after a disturbance (local stability). Across the broad competitive regime the bound on pairwise interaction defines (Theorem 1), the link is tight in one direction: an underyielding community is almost always unstable (Fig. 2). However, they are not the same property, and mapping where they part brings particular stabilizing and destabilizing factors to light.

Two particular cases were flagged above and deserve to be investigated further. The first case is stable underyielding (purple curve, Fig. 3C), where two species compete too strongly to coexist on their own but are rescued through an indirect interaction pathway involving a third fast-growing species. Such a community is therefore stabilized by the ordering of characteristic time scales itself—a dimension that the classical stabilizing ingredients of coexistence theory, niche and fitness differences^13^, do not capture: neither carrying capacities nor interaction strengths need change for the coexistence to appear or vanish, only the rates at which species approach their equilibria (SI S3.3.2). The second case is compositional recovery (blue curve, Fig. 3C; its rarity, Fig. 3D), where RYT dips below one only in passing—while pairs violating the interaction bound of the theorem are transiently dominant—before the community settles at an overyielding state with every species kept. Both cases are forms of emergent coexistence^36^: outcomes that cannot be deduced from the pairwise interactions alone and that arise instead through indirect pathways within the community—time-scale ordering being, to our knowledge, an unrecognized route to it. Both are routes by which our empirical test can return a false positive—an underyielding observation that no species loss follows. Our results suggest, however, that both are vanishingly rare and require finely tuned conditions that we characterize analytically in full (SI S3.3.2 and S3.6).

A third boundary case runs the other way, and it is neither rare nor easily foreseen. Theorem 1 runs in one direction only: feasibility, under the pairwise interaction bound, forces overyielding—it does not promise that an overyielding community is stable. For instance, a few species whose net effect on each other is positive can form a coalition that, flourishing together, drives the rest away from equilibrium (SI S3.3.1), even though every pairwise interaction respects the bound. Such instabilities cannot be read from the pairwise interactions alone, and they are not marginal (Fig. 2, SI S3.3.1). This asymmetry is precisely why our empirical test rests on the contrapositive of Theorem 1: RYT < 1 is an early-warning signal, since feasibility or stability must already be failing, whereas RYT > 1 marks a strong potential for stable coexistence, not a guarantee of it.

### Assumptions and limits of the framework

The framework inherits the assumptions of the classical generalized Lotka–Volterra model, such as linearity of per capita interactions. This is a genuine restriction: the two-species analysis already showed that non-linear functional responses can sever the link between overyielding and stable coexistence^6^. In practice, though, the linear model predicts mixture yields well across plant communities^37^, and it is this tractable form that carries the two-species link between complementarity and coexistence^6^ up to entire communities.

Complementarity and relative yields were built for a single trophic level—plant communities above all—and no equivalent measure yet exists for ecosystems spanning several trophic levels^38^; extending one would carry the diagnosis from plant communities to whole food webs. Moreover, because the complementarity effect, and thus RYT, is defined only relative to monoculture yields, it cannot be computed for the many communities observed without them. Partial-sampling estimators^39^ and the mechanistic dissection of complementarity^12^ are imperfect but workable routes around this requirement and would let our complementarity diagnosis be read from observational data alone.

### Biodiversity that endures cannot but benefit ecosystem functioning through complementarity

Two questions opened this paper: what diversity does for an ecosystem, and whether that diversity can persist. Complementarity, we have shown, binds the two together. Many studies have established that biodiversity is, on the whole, beneficial for ecosystem functioning^40–43^. Here we find something stronger: biodiversity that endures cannot but be beneficial through complementarity. Underyielding is, for all practical purposes, not a resting state, and a community caught in it is one that is losing species. The positive and increasing relationship between biodiversity and ecosystem functioning that decades of experiments have measured is, in this light, the steady state of a self-regulating filter rather than a fortunate tendency.

## Methods

### Theoretical framework

We consider communities of *S* species governed by the generalized Lotka–Volterra system. To straightforwardly link population dynamics to the complementarity effect^5^, we adopted the parametrization making explicit species monoculture carrying capacities *K*_*i*_ > 0,

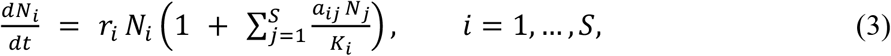

with intrinsic growth rates *r*_*i*_ > 0, and normalized interaction coefficients *a*_*ij*_ satisfying *a*_*ii*_ = −1. Note that the normalized interaction coefficients *a*_*ij*_ are, by definition, the *per capita* interspecific interactions relative to intraspecific competition^25^. That is, this *K*-parameterization makes the *a*_*ij*_ truly dimensionless and habitat-invariant^25^, and requires *r*_*i*_ > 0 for the carrying capacities themselves to be well-defined^44^—a condition satisfied by construction in BEF assemblages, where we assume every species grows in monoculture (SI S1). It covers any mixture of competition and facilitation among species that each persist in isolation (*K*_*i*_ > 0); obligate predator–prey interactions, which involve a consumer unable to survive alone, fall outside its scope. The equilibrium solves ***AN***^*^ = −***K***, with ***A*** = (*a*_*ij*_) and ***K*** = (*K*_*i*_). The equilibrium is *feasible* when 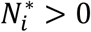 for all species *i*.

### Complementarity and relative yield total

The relative yield of species *i* is 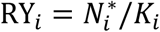 and the relative yield total is RYT = ∑_*i*_ RY_*i*_ (11, 23). The relative yield total is related to the complementarity effect 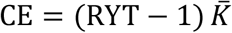 under the standard assumption of uniform expected proportions (SI S1.3). Because each relative yield is referenced to its species’ own contemporaneous monoculture, a decline driven by the shared environment lowers mixture and monoculture biomass together and leaves RYT largely unchanged; a fall in RYT therefore reflects a change in how species interact, not a deterioration of conditions common to mixture and monoculture. We carry the complementarity effect through the analysis in this relative-yield form because RYT is dimensionless, comparable across systems of different productivity, and—being strictly positive—admits the logarithm used in the empirical analyses, whereas the additive complementarity effect is scaled by mean monoculture yield and turns negative under underyielding, so it is neither dimensionless nor log-transformable. The theorem, simulations, and statistics are therefore stated in RYT, while the complementarity effect supplies their logical reading.

### Potential of stable coexistence

Stable coexistence requires the existence of a feasible and stable equilibrium. The potential of feasibility is quantified by feasibility-domain size Ω, which is the normalized solid-angle volume of the cone of carrying-capacity vectors compatible with ***N***^*^ > 0 (9, 10) (SI S1.4). That is, a larger Ω implies a larger potential of feasibility, as it has been stated that communities with larger Ω can undergo larger environmental variations without species extinction^45^. If, in addition, the feasible equilibrium ***N***^*^ is *locally stable*, i.e., all eigenvalues of the Jacobian ***J***^*^ = diag(*r*_*i*_ RY_*i*_) ***A*** at ***N***^*^ have negative real parts, the community achieves *stable coexistence*. The more negative the leading eigenvalue *λ*_max_, the further ***N***^*^ is from being unstable. Thus, the potential of stable coexistence is quantitatively linked to Ω and *λ*_max_.

### Non-equilibrium analysis

The non-equilibrium counterpart of our analysis—how a relative yield total measured away from equilibrium still reports the interactions, and why a transient underyielding community is mainly bound to lose a species—is developed analytically in SI S3.6, while the simulations behind Fig. 3, including the sampling behind panel D and the robustness of our results against heterogeneity of growth rates, are detailed in SI S4.2 and SI S3.6.4.

### Datasets

We analyzed four long-term biodiversity experiments providing species-resolved biomass and contemporaneous monoculture references: the Pennekamp et al. protist microcosms^26^ (580 mixture microcosms, six bacterivorous species, six experimental temperatures from 15 to 25 °C, 19 observations over 40 days); Cedar Creek E120 (27, 28) (56 mixture plots, 16-species perennial-prairie pool, annual species-sorted sampling 2001–2021, except 2009); the Dominance Experiment of the Jena Experiment^29,46^ (188 plots); and the Agrodiversity Experiment^30,31^ (796 plots at 28 of the experiment’s 33 sites, 27 in Europe and one in Canada; grass–legume mixtures, multiple harvests per year). Dataset descriptors, sampling intervals, and harmonization details are given in SI S6 (SI Table S1). All datasets were harmonized into unit–time panels with consistent species resolution. Throughout, the species considered in a unit are those sown or introduced in it. For each unit, the reference community was defined as the set of species present at the first eligible observation *t*_0_; species-level biomass was never inferred from unresolved observations. Mixture-only filtering (sown richness *S*_init_ > 1) was applied throughout. Monoculture references were computed contemporaneously. Dataset-specific protocols (notably averaging across replicate monocultures at each species × temperature × day combination for the Pennekamp microcosms) are described in SI S6.

### Statistical analysis

For each unit *p* (a field plot, or a microcosm for the protist data) and observation time *t*, the binary exposure was underyielding over two strictly adjacent observations,

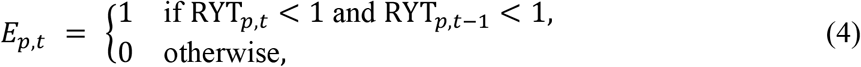

where *t* and *t* − 1 are consecutive observations (difference of exactly one in the temporal index). A persistent-extinction event was recorded when a species present in the previous observation was absent at the focal time and also at the immediately following observation; this requires three consecutive observations, so events at the first (*t*_0_) and last observation of each unit were excluded from event detection. The outcome *Y*_*p,t,h*_ was set to one if at least one species of the reference community of unit *p* underwent a persistent-extinction event within the prospective window (*t, t* + *h*], and zero otherwise. The primary horizon was *h* = 2, which leaves a declining species time to be lost; *h* = 1 and *h* = 3 are reported as sensitivity analyses (SI S7.4). Horizons count observations: years in the two grassland experiments, harvests in the agricultural experiment and sampling dates in the microcosms.

For each dataset, we fitted two binomial regressions to assess the effect of underyielding *E*_*p,t*_ and the total RYT on the further extinction probabilities *Y*_*p,t,h*_. The two statistical models read as

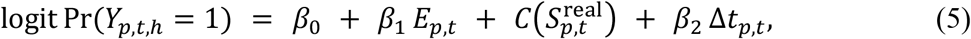

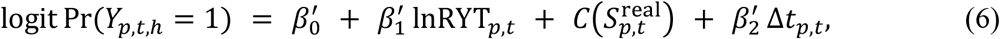

where exp(*β*_1_) is the odds ratio of further extinction when underyielding is observed, and *β*^′^ is the change in the log-odds of further extinction per unit increase in lnRYT_*p,t*_ (a decrease when negative) . The other terms are 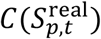, a categorical adjustment for realized species richness, the number of species present at *t* (SI S7.2), and Δ*t*_*p,t*_, the time since the reference observation *t*_0_ (in days for the microcosms, in years otherwise). For the Pennekamp microcosms the linear predictor additionally included a six-level fixed effect *C*(temperature) for the experimental temperature gradient (15–25 °C), the principal design factor of that study.

The binary community odds ratios (Fig. 4A) are estimated by Firth penalized logistic regression^47^; the continuous coefficient 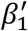 is estimated by cluster-robust generalized estimating equations (GEE)^48^. The working-correlation fallback, the persistence-filter and horizon robustness checks, the Firth–GEE cross-walk, and the full estimator specification are given in SI S7.

To summarize each effect across the four experiments we combined the dataset-specific estimates by random-effects meta-analysis. Between-experiment heterogeneity *τ*^2^ was estimated by the DerSimonian–Laird method^49^; confidence intervals for the pooled effect use the modified Hartung–Knapp–Sidik–Jonkman small-sample adjustment^50,51^ (SI S7.5), referring the pooled estimate to a *t* distribution on *k* − 1 = 3 degrees of freedom. The pooled estimates are the gray *Pooled (RE)* diamonds in Fig. 4A, B; the heterogeneity *I*^2^ of the pooled odds ratios is given in SI Table S4, and full meta-analytic detail in SI S7.5.

To test whether underyielding carries information beyond the rarity of the species that is lost, we fitted a species-level model at *h* = 2. The unit is a species *i* of the reference community present at a baseline *t*, with RY_*i,p,t*_ > 0, and the outcome *Y*_*i,p,t*_ is its own persistent extinction within (*t, t* + 2], under the eligibility and window rules of the community test:

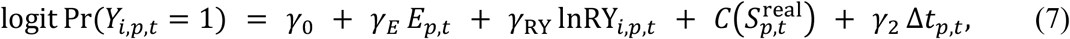

with *C*(temperature) for the microcosms. It is estimated by GEE with an independence working correlation and sandwich standard errors clustered by unit. exp(*γ*_*E*_) is the odds ratio of losing a species after underyielding, at that species’ own relative yield; *n* is the number of species-baselines (Fig. 4C). We calibrated it against a rarity-only null model. A Firth logistic regression of species loss on deciles of lnRY_*i,p,t*_, *C*(*S*^real^) and Δ*t*_*p,t*_ (and temperature), with no community term, gave each species’ probability of loss. Losses were then drawn as independent Bernoulli variables, keeping all relative yields, exposures and the plot structure as observed, and exp(*γ*_*E*_) was re-estimated on each of 10,000 simulated datasets. Datasets in which the model could not be fitted (separation or non-convergence: 6.8% for the microcosms, at most 0.02% elsewhere) were discarded. Underyielding was taken to carry information beyond rarity when the observed exp(*γ*_*E*_) exceeded the 95th percentile of its null distribution.

All tests are two-sided, and all intervals are 95% confidence intervals. Dataset-specific *P* values are Wald tests, based on penalized-likelihood standard errors for the Firth odds ratios and on robust (sandwich) standard errors for the GEE coefficients; pooled *P* values and intervals refer the pooled estimate to a *t* distribution with three degrees of freedom. No correction for multiple comparisons was applied. The sampling unit is the unit–time baseline: *n* is the number of baselines in the estimation sample of a dataset and horizon, and the number of events is the number of baselines followed by at least one persistent extinction within the window. Both are given in Fig. 4A, B (*h* = 2) and in SI Table S2 (*h* = 3); *n* is also given in SI Table S1 (*h* = 1 and 2); the baselines come from 580 microcosms (Pennekamp et al.), 56 plots (Cedar Creek), 188 plots (Jena Dominance) and 796 plots (Agrodiversity). Because the windows of successive baselines of the same unit overlap, the Firth odds ratios were checked against cluster-robust GEE estimates of the same quantity (SI Table S4) and, for Cedar Creek at two years, against a cluster bootstrap that resamples units with replacement (1,500 resamples, percentile interval). Spearman correlations in Fig. 2 are descriptive and were not tested. Analyses were run in Python 3.13 (numpy 2.4.4, pandas 3.0.2, scipy 1.18.1, statsmodels 0.14.6); the Firth regression and the random-effects meta-analysis were implemented by the authors and are included in the archived code. The simulations of Fig. 2 were run in R with the packages deSolve, mvtnorm and CVXR.

### Use of large language models

A large language model was used to assist with writing and checking analysis code and scripts, with translation, and with improving the language of the manuscript. The authors designed the study, derived the theoretical results, reviewed all code and outputs, and take full responsibility for the content of this article.

### Reporting summary

Further information on research design is available in the Nature Portfolio Reporting Summary linked to this article.

## Data availability

All data used in this study are publicly available. Cedar Creek E120 data are archived at the Environmental Data Initiative^52^. Agrodiversity Experiment data are archived in Ecological Archives^31^. Jena Experiment biomass data are archived at PANGAEA^46^; the assignment of species to the small monoculture plots is available at https://store.pangaea.de/Publications/Jena_Experiment/PlotInformationSmallMonos.txt. Pennekamp et al. protist data are archived at Zenodo under a CC0 license^53^. The compilation of pairwise competition coefficients of Adler et al. used in SI S5 is archived at Dryad under a CC0 license^54^.

## Code availability

All code used for the analyses, simulations and figures is archived at Zenodo^55^ and will be made public upon publication; editors and referees can access it through https://zenodo.org/records/23322417?token=eyJhbGciOiJIUzUxMiJ9.eyJpZCI6IjM4NDdkZDM5LWJiMDEtNDZlYS1iMTU0LTM2MjQwMmJmOTA2MiIsImRhdGEiOnt9LCJyYW5kb20iOiI5NDgyYWY4MzFmOThhNmY0OTY4MjRkZWQ1NzdkNWY2NiJ9.MkVsIA3sC5_Fjj5mjVe6OsoBRNFEmP1jyyu7GswnCKGzb7eBoROHw9kMjc-vSKCa3ZljW9QOfKd3y-b6CIbjVQ. The archive also contains the outputs of the Fig. 2 simulations and the community ensembles of Figs. S4–S5, which were not run with a fixed random seed and are therefore provided as data. The code is released under the MIT license and the simulation outputs under a CC BY 4.0 license.

## Author contributions

Conceptualization: M.D., R.P.R., M.L., J.-F.A.; Methodology: M.D., J.-F.A., R.P.R.; Investigation: M.D.; Software: M.D., R.P.R.; Formal analysis: M.D., J.-F.A., R.P.R.; Validation: M.D., R.P.R., M.L.; Data curation: M.D.; Visualization: M.D.; Writing – original draft: M.D.; Writing – review & editing: M.D., M.L., J.-F.A., R.P.R.; Funding acquisition: R.P.R.; Supervision: R.P.R.

## Additional information

Supplementary Information is available for this paper.

