## Supplementary Information for "Mixtures of stably coexisting species overyield"

Supplementary Text ........ S4
S1 Generalized Lotka–Volterra framework ........ S4
S1.1 Generalized Lotka–Volterra equations in the $K$-parameterization ........ S4
S1.2 Biodiversity–ecosystem-functioning quantities ........ S5
S1.3 Equivalence between RYT and the complementarity effect ........ S5
S1.4 Stability and potential of feasibility measures ........ S7
S2 Two-species case ........ S9
S2.1 Derivation of $\lambda_{\max} = -r\,(\mathrm{RYT} - 1)$, when growth rates are equal to $r$ ........ S9
S2.2 Reading the potential of feasibility Ω in the phase space ........ S10
S2.3 The Ω–RYT relation in two species ........ S10
S2.4 Away from equilibrium: underyielding in two species ........ S12
S3 Multispecies case ........ S16
S3.1 Equal-growth-rate case: stability requires overyielding ........ S16
S3.2 Theorem on feasibility ........ S17
S3.2.1 The pairwise interaction bound ........ S17
S3.2.2 Theorem 1: feasibility implies overyielding under bounded interactions ........ S17
S3.3 Stability implications of the pairwise interaction bound ........ S19
S3.3.1 The symmetric part governs stability ........ S19
S3.3.2 Stable underyielding case ........ S20
S3.3.3 Why strong facilitative regimes are filtered ........ S22
S3.4 Solvable mean-field case for S species ........ S22
S3.5 Quantitative relations between RYT and stable coexistence ........ S23
S3.5.1 An analytical $\lambda_{\max}$–RYT bound at homogeneous growth ........ S23
S3.5.2 An analytical Ω–RYT relation in the mean-field case ........ S29
S3.6 Non-equilibrium analysis ........ S31
S3.6.1 The model in relative yields ........ S31
S3.6.2 What a relative yield total below one says about the interactions ........ S31
S3.6.3 The total settles before the composition ........ S32
S3.6.4 Unequal growth rates ........ S33
S3.6.5 The loss of a species ........ S34
S4 Numerical simulations ........ S35
S4.1 Equilibrium ensembles ........ S35
S4.2 Dynamical ensembles ........ S36
S5 Empirical assessment of the interaction bound ........ S37

S6 Empirical datasets ........ S39
S6.1 The four datasets ........ S39
S6.2 Panel construction ........ S40
S7 Statistical analysis ........ S42
S7.1 The regression model ........ S42
S7.2 Adjustment variables and binning ........ S42
S7.3 Statistical caveats ........ S43
S7.4 Robustness and power ........ S43
S7.5 Firth penalized estimation and the GEE cross-walk ........ S44

### Supplementary Text

#### S1 Generalized Lotka–Volterra framework

This section restates, in compact form, the modeling conventions and coexistence metrics used throughout the main text and the supplementary material. Its purpose is to remove any residual ambiguity in sign or scaling.

**Table of contents of S1**

S1.1 Generalized Lotka–Volterra equations in the $K$-parameterization ................................ S4

S1.2 Biodiversity–ecosystem-functioning quantities ............................................................ S5

S1.3 Equivalence between RYT and the complementarity effect ........................................ S5

S1.4 Stability and potential of feasibility measures ............................................................ S7

##### S1.1 Generalized Lotka–Volterra equations in the $\boldsymbol{K}$-parameterization

We consider communities of $S \geq 2$ species governed by the generalized Lotka–Volterra (gLV) system. The biomass dynamics of $N_i$ are given by

$$\frac{dN_i}{dt} = r_i N_i \left(1 + \sum_{j=1}^{S} \frac{a_{ij} N_j}{K_i}\right), \qquad i = 1, \ldots, S, \tag{S1}$$

with intrinsic growth rates $r_i > 0$, monoculture carrying capacities $K_i > 0$, and normalized interaction coefficients $a_{ij}$ collected in the matrix $\boldsymbol{A} = (a_{ij})$. Here, we adopted the $K$ notation for the gLV model[25]. Note that in this notation, the interactions are normalized by the intraspecific competition. That is, $a_{ii} = -1$ and $a_{ij}$ is the per capita effect of species $j$ on species $i$ normalized by species $i$ intraspecific competition[25].

**Why this $K$-parameterization.** We use the carrying-capacity form (Eq. S1), rather than the equivalent $dN_i/dt = N_i\ (r_i - \sum_j \alpha_{ij} N_j)$ form with implicit carrying capacities, for two reasons[25]. First, $a_{ij}$ is truly dimensionless and invariant to the size of the habitat in which the community is observed, which is a prerequisite for any statement that aggregates interaction data across systems (as we do in Sec. S5). Second, strictly positive carrying capacities $K_i > 0$ are required for the relative yields $\mathrm{RY}_i = N_i^*/K_i$ and the relative yield total to be defined; the $K$-parameterization makes this requirement explicit. The $K_i$ are themselves well-defined only when $r_i > 0$ (44), which is the regime of the present manuscript, since every species in a biodiversity–ecosystem functioning (BEF) assemblage has, by definition, positive intrinsic growth in monoculture. Note that a negative interaction value $a_{ij} < 0$ represents a negative effect of species $j$ on species $i$, and vice versa for a positive interaction.

**Equilibrium, feasibility, and stability.** The interior equilibrium solves

$$\boldsymbol{A}\boldsymbol{N}^* = -\boldsymbol{K}, \qquad \boldsymbol{K} = (K_1, \ldots, K_S)^\top. \tag{S2}$$

The community is *feasible* when $N_i^* > 0$ for every $i$, and *locally asymptotically stable* when all eigenvalues of the Jacobian at $\boldsymbol{N}^*$,

$$\boldsymbol{J}^* = \mathrm{diag}(r_i\,\mathrm{RY}_i)\,\boldsymbol{A}, \qquad \mathrm{RY}_i = N_i^*/K_i, \tag{S3}$$

have strictly negative real parts. A community is said to *stably coexist* when both conditions hold. The leading Jacobian eigenvalue is denoted $\lambda_{\max} = \max_i \mathrm{Re}\,\lambda_i(\boldsymbol{J}^*)$.

##### S1.2 Biodiversity–ecosystem-functioning quantities

We assume the community to hold a feasible and stable equilibrium $N_i^* > 0$. The *relative yield* of species $i$ in mixture is defined, following Vandermeer[23] and Loreau[11], as the ratio of its equilibrium biomass to its monoculture carrying capacity:

$$\mathrm{RY}_i = \frac{N_i^*}{K_i}. \tag{S4}$$

The *relative yield total* is the sum across species,

$$\mathrm{RYT} = \sum_{i=1}^{S} \mathrm{RY}_i = \sum_{i=1}^{S} \frac{N_i^*}{K_i}. \tag{S5}$$

A community is said to *overyield* when $\mathrm{RYT} > 1$ and to *underyield* when $\mathrm{RYT} < 1$.

Intuitively, each $\mathrm{RY}_i$ measures the fractional occupation of species $i$'s monoculture niche when it is grown in the mixture: if species $i$ alone fills its niche entirely, $\mathrm{RY}_i = 1$; if mixture conditions limit it to half of that, $\mathrm{RY}_i = 0.5$. Summing across species, RYT is therefore the *total niche occupation* of the mixture, expressed in monoculture-equivalents. Importantly, RYT is not a difference of biomasses (which is the construction of the net biodiversity effect of the Loreau–Hector partition; see Sec. S1.3 for the exact algebraic link), but a sum of species-level ratios with no reference to mixture biomass on a common scale. A value $\mathrm{RYT} > 1$ then means that the mixture collectively occupies more than one monoculture's worth of niche space[11,12], the operational signature of niche differentiation—including reduced shared natural enemies—or facilitation; a value $\mathrm{RYT} < 1$ means that the sum of niche occupations falls short of a single monoculture, which is the empirical signal whose dynamical consequences this manuscript sets out to characterize.

In experimental settings, $\mathrm{RY}_i$ is computed as the ratio of the observed mixture biomass of species $i$ to its observed monoculture biomass under matched conditions; the algebraic identity with $N_i^*/K_i$ holds at equilibrium and is the operational identification used throughout the empirical analyses of the main text.

Finally, note that the relative yields and their total can also be computed out of equilibrium by using the biomass $N_i(t)$ at time $t$, instead of the equilibrium value. That is $\mathrm{RY}_i(t) = N_i(t)/K_i$ and $\mathrm{RYT}(t) = \sum_i \mathrm{RY}_i\,(t)$.

##### S1.3 Equivalence between RYT and the complementarity effect

The relative yield total is not an *ad hoc* construction: it coincides exactly with the complementarity component of the biodiversity-effect partition of Loreau and Hector, under the standard assumption of equal expected relative yields $\mathrm{RY}_i^E = 1/S$. We make the two equivalences—one in additive (difference) form, one in ratio form—explicit, as the discussion of the main text relies on this anchor to interpret $\mathrm{RYT} > 1$ as the observable signature of the mechanisms (niche differentiation, including reduced shared natural enemies, and facilitation) that the complementarity literature has documented for two decades[12,56].

**Additive partition[5].** Let $\mathrm{RY}_i = N_i^*/K_i$ be the realized relative yield of species $i$ in mixture, and let $\mathrm{RY}_i^E$ be its expected relative yield in the absence of biodiversity effects—the fixed fraction of its monoculture carrying capacity that each species would realize under the null. With $\bar{K} = 1/S \sum_i K_i$ the arithmetic mean of the monoculture carrying capacities, the Loreau–Hector additive partition[5] writes the net biodiversity effect as

$$\mathrm{NBE} \;=\; B - B^E \;=\; \underbrace{\bar{K} \cdot \textstyle\sum_i (\mathrm{RY}_i - \mathrm{RY}_i^E)}_{\text{complementarity term CE}} \;+\; \underbrace{S \cdot \mathrm{cov}(K_i,\, \mathrm{RY}_i - \mathrm{RY}_i^E)}_{\text{selection term SE}}, \qquad \text{(S6)}$$

where $B = \sum_i N_i^* = \sum_i K_i \ \mathrm{RY}_i$ is the mixture biomass, $B^E = \sum_i K_i \ \mathrm{RY}_i^E$ is its expected counterpart, and the covariance is taken across species (the selection effect SE in the historical Loreau–Hector form).

Under the standard assumption $\mathrm{RY}_i^E = 1/S$ for every species (so $\sum_i \mathrm{RY}_i^E = 1$), and using $\sum_i \mathrm{RY}_i = \mathrm{RYT}$, the additive complementarity effect reduces to

$$\mathrm{CE} \;=\; \bar{K}\,(\mathrm{RYT} - 1), \qquad \text{(S7)}$$

which gives the exact algebraic equivalence

$$\mathrm{CE} \;>\; 0 \quad \Leftrightarrow \quad \mathrm{RYT} \;>\; 1, \qquad \text{(S8)}$$

together with the proportional relation $|\mathrm{CE}| = \bar{K}\,|\mathrm{RYT} - 1|$ in magnitude. The sign of the complementarity effect is the sign of $\mathrm{RYT} - 1$, and its magnitude is the deviation from unity rescaled by the mean monoculture carrying capacity.

**Ratio partition.** The same conclusion is reached, with a stronger algebraic content, when biodiversity effects are partitioned in ratio form rather than in difference form[24,57]. The ratio-based complementarity component of $B/B^E$ is defined as

$$\widetilde{\mathrm{CE}} \;=\; \frac{\sum_i \mathrm{RY}_i}{\sum_i \mathrm{RY}_i^E}. \qquad \text{(S9)}$$

Under the hypothesis $\sum_i \mathrm{RY}_i^E = 1$, we obtain

$$\widetilde{\mathrm{CE}} \;=\; \mathrm{RYT} \qquad \text{(S10)}$$

exactly—not only in sign but in magnitude. The ratio complementarity effect and the relative yield total are the same quantity once expected relative yields are uniform.

**Synthesis.** Equations S7 and S10 establish that RYT directly relates to the complementarity effect of the Loreau–Hector partition—proportional to it in additive form, identical to it in ratio form.

Throughout the manuscript we therefore treat $\mathrm{RYT} > 1$ and $\mathrm{CE} > 0$ as equivalent statements about the sign of complementarity, and use RYT as the canonical observable: it is dimensionless, and aggregates the contributions of all species into a single ratio with a direct interpretation as total monoculture-equivalent niche occupation (Sec. S1.2).

#### S1.4 Stability and potential of feasibility measures

Stable coexistence requires the existence of a feasible and stable equilibrium, which relates to two complementary notions that can be computed. The leading Jacobian eigenvalue is a measure of stability, while the potential of feasibility is measured by the size of the so-called feasibility domain.

**Leading Jacobian eigenvalue $\lambda_{\max}$.** As defined in Eq. S3, $\lambda_{\max}$ is the largest real part of the eigenvalues of $\boldsymbol{J}^*$. The equilibrium is locally asymptotically stable if and only if $\lambda_{\max} < 0$, and the magnitude $|\lambda_{\max}|$ measures a *distance to the stability boundary*: how far the community sits from the switch between stability and instability. This quantity is also the asymptotic decay rate of the slowest mode. That long-term rate, however, need not equal the response seen at short times[8].

**Feasibility-domain size $\Omega$.** For an interaction matrix $\boldsymbol{A}$, the domain of feasibility is defined, in the $K$-parameterization of the generalized Lotka–Volterra model, as the set of carrying-capacity vectors $\boldsymbol{K}$ leading to feasibility, i.e., such that $\boldsymbol{N}^* = -\boldsymbol{A}^{-1}\boldsymbol{K} > 0$. Mathematically, the feasibility domain is thus defined by

$$\mathcal{D}(\boldsymbol{A}) = \{\boldsymbol{K} = x_1\boldsymbol{v_1} + x_2\boldsymbol{v_2} + \cdots + x_S\boldsymbol{v_S} \quad | \quad x_1, x_2, \ldots, x_S > 0\}, \tag{S11}$$

where the vectors $\boldsymbol{v_1}, \ldots, \boldsymbol{v_S}$ are given by minus the columns of the interaction matrix $\boldsymbol{A}$,

$$\boldsymbol{A} = \begin{bmatrix} \vdots & \vdots & & \vdots \\ -\boldsymbol{v_1} & -\boldsymbol{v_2} & \cdots & -\boldsymbol{v_S} \\ \vdots & \vdots & & \vdots \end{bmatrix} \tag{S12}$$

Geometrically, the feasibility domain is a convex cone, given by all strictly positive linear combinations of the vectors $\boldsymbol{v_1}, \ldots, \boldsymbol{v_S}$. Figure S1 illustrates this domain in dimension $S = 2$ and $S = 3$, and Fig. 1B of the main text in dimension $S = 2$. Finally, the opening of the cone quantifies the potential of feasibility, i.e., how likely it is to find a combination of carrying capacities leading to feasibility. The opening of the cone is simply quantified by its normalized solid angle $\Omega$. We normalize the solid angle such that, in the case where species do not interact, i.e., the interaction matrix is $-1$ on the diagonal and 0 outside the diagonal ($\boldsymbol{A} = -\boldsymbol{Id}$), the normalized solid angle $\Omega = 1$. In this case, the domain of feasibility is simply all possible combinations of strictly positive carrying capacities ($K_i > 0$), i.e., $\mathcal{D}(\boldsymbol{A} = -\boldsymbol{Id}) = \mathbb{R}^S_{>0}$.

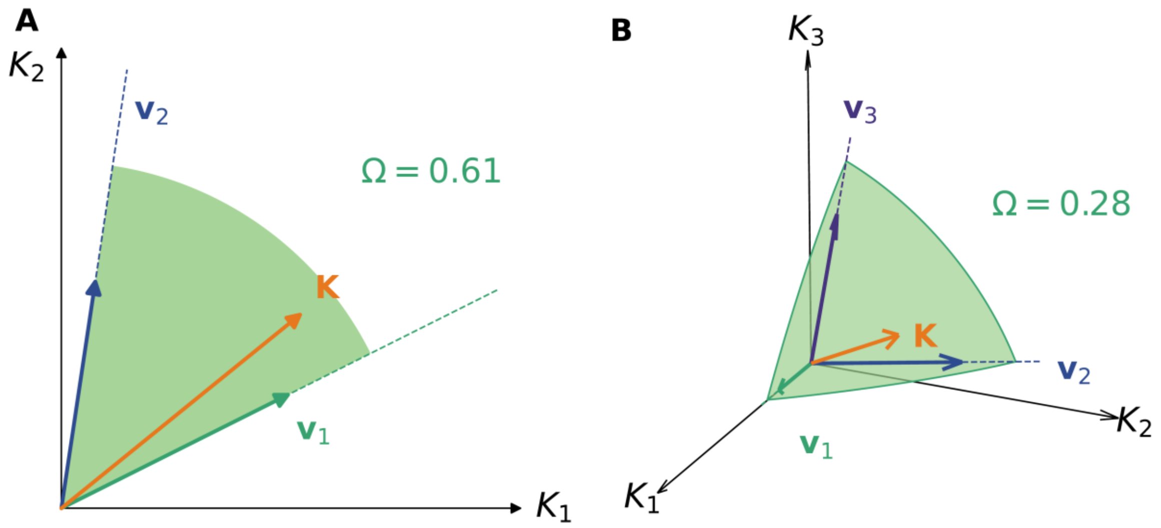


**Fig. S1. Geometric representation of the potential of feasibility for two and three species.** Both panels illustrate the feasibility domain. The feasibility domain is the convex cone generated by the vectors $\boldsymbol{v}_i$ ($\boldsymbol{v}_1$ in green, $\boldsymbol{v}_2$ in blue, $\boldsymbol{v}_3$ in purple), determining the interaction matrix $\boldsymbol{A}$, within the space of carrying capacities ($K_i$) (black axes). If the vector of carrying capacities $\boldsymbol{K}$ (in orange) lies within the cone, the system admits a feasible equilibrium point. The opening of the cone, measured by the normalized solid angle Ω (see text), is a measure of the potential of coexistence, i.e., how large is the set of $K_i$ allowing for feasibility.

In dimension $S = 2$, Ω is directly derived from the dot product between the vectors $\boldsymbol{v_1}$ and $\boldsymbol{v_2}$:

$$\Omega(\boldsymbol{A}) = \frac{2}{\pi} \cos^{-1}\left(\frac{\boldsymbol{v_1}\cdot\boldsymbol{v_2}}{||\boldsymbol{v_1}||\,||\boldsymbol{v_2}||}\right). \tag{S13}$$

In general, in dimension $S$, Ω is computed as the following multivariate integral:

$$\Omega(\boldsymbol{A}) = \frac{2^S\,|\det\boldsymbol{A}|}{\pi^{S/2}} \int \cdots \int_{\mathbb{R}^S_{>0}} e^{-\boldsymbol{x}^T\boldsymbol{A}^T\boldsymbol{A}\boldsymbol{x}}\, d\boldsymbol{x} \tag{S14}$$

This integral cannot be computed explicitly, but efficient numerical methods, based on the quasi-Monte-Carlo method, do exist[10,58,59].

Intuitively, Ω quantifies the breadth of environments—as carrying-capacity configurations—that the interaction matrix admits as feasible: a community with large Ω tolerates wide environmental variation before some species drops out, while $\Omega \to 0$ marks the collapse of the feasibility cone to a singular ray, i.e., the boundary at which the community cannot persist except for one fine-tuned environment. For instance, if the interactions are dominated by competition, then the cone reduces in size and Ω is very likely to be smaller than 1, while positive interactions can open the cone, and Ω may reach a value larger than 1. The potential of feasibility Ω is intrinsic to $\boldsymbol{A}$ in the sense that it is independent of the carrying-capacity vector $\boldsymbol{K}$ once the matrix is specified.

### S2 Two-species case

In two species, RYT relates analytically to the two measures of coexistence potential introduced in Sec. S1.4: the leading Jacobian eigenvalue $\lambda_{\max}$ and the feasibility-domain size Ω. This section derives these relations and supplies the analytical envelope behind Fig. 1C and Fig. 1D, and Fig. 3A, B of the main text.

**Table of contents of S2**


S2.1 Derivation of $\lambda_{\max} = -r\,(\mathrm{RYT}-1)$, when growth rates are equal to $r$ ...................... S9

S2.2 Reading the potential of feasibility Ω in the phase space ........................................... S10

S2.3 The Ω–RYT relation in two species .......................................................................... S10

S2.4 Away from equilibrium: underyielding in two species ............................................. S12


#### S2.1 Derivation of $\boldsymbol{\lambda_{\max} = -r\,(\mathrm{RYT}-1)}$, when growth rates are equal to $\boldsymbol{r}$

Consider two interacting species with arbitrary interaction coefficients $a_{12}, a_{21}$ satisfying $a_{12} + a_{21} > -2$. Let the equilibrium $\boldsymbol{N}^*$ be feasible. The Jacobian at $\boldsymbol{N}^*$ from Eq. S3 reads as

$$\boldsymbol{J}^* \;=\; \mathrm{diag}(r_1\mathrm{RY}_1, r_2\mathrm{RY}_2)\begin{pmatrix}-1 & a_{12}\\ a_{21} & -1\end{pmatrix} \;=\; \begin{pmatrix}-r_1\mathrm{RY}_1 & r_1\mathrm{RY}_1 a_{12}\\ r_2\mathrm{RY}_2 a_{21} & -r_2\mathrm{RY}_2\end{pmatrix}. \qquad \text{(S15)}$$

The two eigenvalues of $\boldsymbol{J}^*$ are

$$\lambda_\pm \;=\; -\,\frac{r_1\mathrm{RY}_1 + r_2\mathrm{RY}_2}{2} \;\pm\; \sqrt{\left(\frac{r_1\mathrm{RY}_1 - r_2\mathrm{RY}_2}{2}\right)^2 + r_1 r_2 \mathrm{RY}_1\mathrm{RY}_2\, a_{12}a_{21}}. \qquad \text{(S16)}$$

When growth rates are equal, $r_1\mathrm{RY}_1 + r_2\mathrm{RY}_2 = r(\mathrm{RY}_1 + \mathrm{RY}_2) = r\,\mathrm{RYT}$. The leading eigenvalue (largest real part) is

$$\lambda_{\max} \;=\; -\,\frac{r}{2}\,\mathrm{RYT} + \sqrt{\left(\frac{r(\mathrm{RY}_1 - \mathrm{RY}_2)}{2}\right)^2 + r^2\mathrm{RY}_1\mathrm{RY}_2\, a_{12}a_{21}}. \qquad \text{(S17)}$$

The equilibrium relation $\boldsymbol{A}\boldsymbol{N}^* = -\boldsymbol{K}$ reads $K_i = N_i^* - a_{ij}N_j^*$ in two species, so

$$1 - \mathrm{RY}_i \;=\; \frac{K_i - N_i^*}{K_i} \;=\; -\,a_{ij}\,\frac{N_j^*}{K_i} \qquad (i \neq j), \qquad \text{(S18)}$$

and multiplying the two gives $(1-\mathrm{RY}_1)(1-\mathrm{RY}_2) = a_{12}a_{21}\,\mathrm{RY}_1\mathrm{RY}_2$. The term under the square root in Eq. S17 thus collapses to a perfect square,

$$(\mathrm{RY}_1 - \mathrm{RY}_2)^2 + 4\,\mathrm{RY}_1\mathrm{RY}_2\, a_{12}a_{21} \;=\; (\mathrm{RY}_1 - \mathrm{RY}_2)^2 + 4(1-\mathrm{RY}_1)(1-\mathrm{RY}_2) \;=\; (\mathrm{RYT} - 2)^2, \qquad \text{(S19)}$$

so the two Jacobian eigenvalues are *exactly*

$$\lambda \;\in\; \{-r\,(\mathrm{RYT}-1),\; -r\} \qquad (r_1 = r_2 = r): \qquad \text{(S20)}$$

Near equilibrium, any small displacement of the community can be partitioned into a few elementary displacements, each of which fades away at its own rate (the eigenvalues); we call these displacements modes. The equation above shows the system admits two modes: the *composition* mode $-r(\mathrm{RYT}-1)$ and the *size* mode $-r$. Under competition each $\mathrm{RY}_i < 1$ (Eq. S18 with $a_{ij} < 0$), so RYT $< 2$ and the composition mode is the leading one:

$$\lambda_{\max} \;=\; -r\,(\mathrm{RYT}-1), \qquad r_1 = r_2 = r,\;\; \mathrm{RYT} \le 2. \tag{S21}$$

This is the identity displayed in Eq. 1 of the main text, exact when growth rates are equal. Transgressive overyielding (RYT $> 2$, reachable under facilitation with $\boldsymbol{K} > 0$) makes the size mode $-r$ leading instead, so that at equal rates the two regimes combine into

$$\lambda_{\max} \;=\; -r\,\min(\mathrm{RYT}-1,\ 1), \qquad r_1 = r_2 = r. \tag{S22}$$

For unequal growth rates, the relation is no longer exact; its scatter grows with growth-rate heterogeneity, and the line marks the average trend across the simulation cloud. Numerical exploration confirms that, for RYT $\le 2$, $\lambda_{\max}$ follows $-(\mathrm{RYT}-1)$ closely throughout the range $r_i \in [1/3,3]$ used in Fig. 1C of the main text; the relation is therefore approximate ($\approx$) rather than exact for heterogeneous $r$.

##### S2.2 Reading the potential of feasibility Ω in the phase space

Here, we demonstrate that the potential of feasibility can also directly be read on the phase space of the Lotka–Volterra model. More precisely, we will demonstrate that the angle between the two zero-growth isoclines (Fig. 1A of the main text) is the same as the angle of the feasibility cone (Fig. 1B of the main text). In panel A, the directions of the two isoclines are given by

$$\boldsymbol{u}_1 = \begin{bmatrix} -K_1 \\ -K_1/a_{12} \end{bmatrix}, \qquad \boldsymbol{u}_2 = \begin{bmatrix} K_2/a_{21} \\ K_2 \end{bmatrix}. \tag{S23}$$

Rescaled, $\tilde{\boldsymbol{u}}_1 = (a_{12}/K_1)\,\boldsymbol{u}_1 = (-a_{12}, -1)^\top$ and $\tilde{\boldsymbol{u}}_2 = (a_{21}/K_2)\,\boldsymbol{u}_2 = (1, a_{21})^\top$ span the same lines. The cosine of the angle between them is then given by

$$\frac{\boldsymbol{u}_1 \cdot \boldsymbol{u}_2}{|\boldsymbol{u}_1|\,|\boldsymbol{u}_2|} = \frac{\tilde{\boldsymbol{u}}_1 \cdot \tilde{\boldsymbol{u}}_2}{|\tilde{\boldsymbol{u}}_1|\,|\tilde{\boldsymbol{u}}_2|} = \frac{-a_{12}-a_{21}}{\sqrt{a_{12}^2+1}\,\sqrt{1+a_{21}^2}}. \tag{S24}$$

In turn, the cosine between the two vectors $\boldsymbol{v}_1$ and $\boldsymbol{v}_2$ determining the cone of feasibility is given by

$$\frac{\boldsymbol{v}_1 \cdot \boldsymbol{v}_2}{|\boldsymbol{v}_1|\,|\boldsymbol{v}_2|} = \frac{-a_{12}-a_{21}}{\sqrt{1+a_{21}^2}\,\sqrt{a_{12}^2+1}}. \tag{S25}$$

As both expressions are equal, the angle of the cone of feasibility is the same as the angle between the two isoclines.

##### S2.3 The Ω–RYT relation in two species

In two species, the feasibility cone is the angular wedge in the $(K_1, K_2)$ plane bounded by the two lines $K_2 = -a_{21}K_1$ and $K_1 = -a_{12}K_2$ (Fig. 1B of the main text). Figure S2 makes this correspondence visually explicit: as the mutual competition strength increases (from $a_{12} = a_{21} = -0.10$ to $-0.90$), the feasibility cone shrinks monotonically (right column), and RYT at the symmetric carrying-capacity point $\boldsymbol{K} = (1,1)$ approaches unity from above (left column). The same geometric mechanism that drives Ω toward zero also drives RYT toward 1.

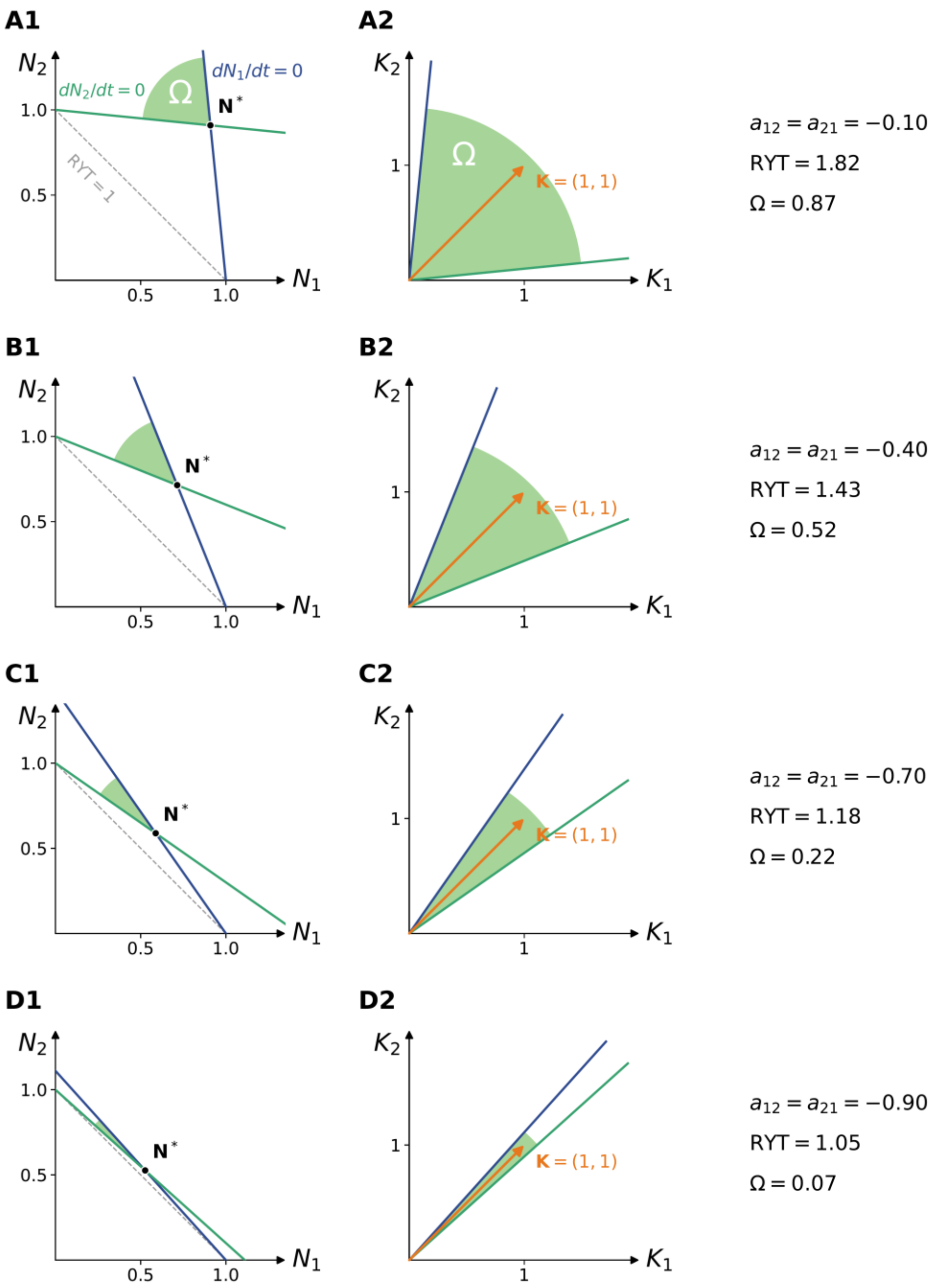


**Fig. S2. Joint shrinking of the feasibility cone and of** RYT **as mutual competition strengthens, in two species.** Four rows of two-species systems, each with diagonal entries $a_{ii} = -1$ and symmetric off-diagonals $a_{12} = a_{21}$ becoming more negative from top to bottom (competition strengthening): **(A)** $a_{12} = a_{21} = -0.10$, yielding RYT = 1.82 and Ω = 0.873; **(B)** $a_{12} = a_{21} = -0.40$, RYT = 1.43, Ω = 0.516; **(C)** $a_{12} = a_{21} = -0.70$, RYT = 1.18, Ω = 0.222; **(D)** $a_{12} = a_{21} = -0.90$, RYT = 1.05, Ω = 0.067. *Left column (A1–D1):* zero-growth isoclines $\mathrm{d}N_1/\mathrm{d}t = 0$ (dark blue) and $\mathrm{d}N_2/\mathrm{d}t = 0$ (dark green) in the $(N_1, N_2)$ plane, with the interior equilibrium $\boldsymbol{N}^*$ (black dot) evaluated at $\boldsymbol{K} = (1,1)$. The shaded light-green wedge at $\boldsymbol{N}^*$ is the angular sector Ω between the isoclines; the dashed gray line marks RYT = 1. Color coding matches Fig. 1A of the main text. *Right column (A2–D2):* the same feasibility cone seen in the $(K_1, K_2)$ plane, with the species-1 and species-2 boundaries of the cone in matching colors and the reference carrying-capacity vector $\boldsymbol{K} = (1,1)$ shown for comparison. Top to bottom, the cone shrinks continuously and monotonically as RYT approaches 1 at the reference point—the geometric content of Eqs. S26–S30.

Beyond the geometric visualization, this can also be seen analytically. Assuming feasibility and stability, the equilibrium relative yields are given by $\mathrm{RY}_i = (1 + a_{ij}K_j/K_i)/(1 - a_{12}a_{21})$ and summing gives,

$$\mathrm{RYT} = \frac{2+(a_{12}K_2/K_1+a_{21}K_1/K_2)}{1-a_{12}a_{21}}. \tag{S26}$$

Within the feasibility cone, RYT is a smooth function of $(K_1, K_2)$ that attains the boundary value $\mathrm{RYT} = 1$ exactly on the edges of the cone, and increases continuously and monotonically with the angular distance from these edges toward the interior of the cone.

Specializing to symmetric interactions $a_{12} = a_{21} = \rho$, the equilibrium relative yield and potential of feasibility Ω are given by

$$\mathrm{RY}_i = \frac{1+\rho\, K_j/K_i}{1-\rho^2} \quad \text{and} \quad \Omega = \frac{2}{\pi}\ \cos^{-1}\left(\frac{-2\rho}{1+\rho^2}\right). \tag{S27}$$

The relative yield total is then given by

$$\mathrm{RYT} = \frac{2+\rho\,(K_1/K_2+K_2/K_1)}{1-\rho^2}. \tag{S28}$$

From here to the end of this subsection we restrict to competition, $-1 < \rho < 0$. Under facilitation ($\rho > 0$), every $\boldsymbol{K} > 0$ remains feasible and RYT → ∞ as $K_1/K_2 \to \infty$—so no maximum exists, and $K_1 = K_2$ gives the smallest attainable total instead. The quantity $\mathrm{RYT_{max}}$ below, and with it Eq. S30 and its $S$-species counterpart (Sec. S3.5.2), are therefore defined only in the competitive regime; the curve drawn for $\mathrm{RYT} > S$ in Fig. 1D is the same parametric relation continued to $\rho > 0$, where it traces the total realized at equal carrying capacities rather than a maximum.

Within that regime, the maximum relative yield total is

$$\mathrm{RYT_{max}} = \frac{2+2\rho}{1-\rho^2} = \frac{2}{1-\rho} \tag{S29}$$

This expression is equivalent to $\rho = 1 - 2/\mathrm{RYT_{max}}$, which we replace in the equation for the feasibility potential and obtain

$$\Omega = \frac{2}{\pi}\ \cos^{-1}\left(\frac{\mathrm{RYT_{max}}\cdot(2-\mathrm{RYT_{max}})}{(\mathrm{RYT_{max}}-1)^2+1}\right). \tag{S30}$$

The same single matrix-level scalar Ω, therefore, indexes both the breadth of carrying-capacity vectors compatible with feasibility and the maximum value of RYT attainable within the cone. The geometric coincidence visualized in Fig. 1D of the main text—the binary feasibility partition at $\mathrm{RYT} = 1$ together with the continuous monotonic Ω–RYT relation within the feasible-stable region—is the analytical content of Eqs. S26–S30.

##### S2.4 Away from equilibrium: underyielding in two species

Here, we first present the reasoning and derivations for the simple 2-species case, illustrated in Fig. 3A, B; while the general case is discussed in Sec. S3.6. We show here two properties: that a RYT below one, even transient, still reports on the interaction between the two species and that once reached, it cannot be undone without the loss of one of the two species.

For two species with equal growth rates $r_1 = r_2 = r$, Eq. S1 reads $dN_i/dt = r\,N_i\,(1 - N_i/K_i + a_{ij}\,N_j/K_i)$. The relative yield of species $i$, $\mathrm{RY}_i = N_i/K_i$, can be read at every moment rather than at equilibrium (Sec. S1.2); being the abundance divided by a constant, it obeys the same equation divided by $K_i$, with the carrying capacities absorbed into the interaction coefficients:

$$\frac{d\mathrm{RY}_1}{dt} = r\,\mathrm{RY}_1\,(1 - \mathrm{RY}_1 + \tilde{a}_{12}\,\mathrm{RY}_2), \qquad \frac{d\mathrm{RY}_2}{dt} = r\,\mathrm{RY}_2\,(1 - \mathrm{RY}_2 + \tilde{a}_{21}\,\mathrm{RY}_1).$$

Each species now has a carrying capacity of one, and $\tilde{a}_{12} = a_{12}K_2/K_1$ measures the effect of species 2 on species 1 in units of species 1's own self-limitation (at equal carrying capacities, $\tilde{a}_{ij} = a_{ij}$); the $S$-species form is Eq. S60.

Rather than following $\mathrm{RY}_1$ and $\mathrm{RY}_2$ separately, we describe the pair by two other numbers: how much relative yield it holds in total, and how that total is shared,

$$\mathrm{RYT} = \mathrm{RY}_1 + \mathrm{RY}_2, \qquad p_1 = \frac{\mathrm{RY}_1}{\mathrm{RYT}}, \qquad p_2 = 1 - p_1.$$

This is only a change of variables, since $\mathrm{RY}_i = \mathrm{RYT}\,p_i$, but the two new coordinates answer different questions: the total is what we measure, and the composition $\boldsymbol{p}$ is where a species is lost, when its share $p_i$ reaches zero.

**Relative yield dynamics.** For RYT, adding the two species differential equations gives its rate of change. The terms linear in the yields pool into RYT; the quadratic ones pool into the self-limitation of each species and the interaction of the pair, counted once from each side:

$$\frac{d\mathrm{RYT}}{dt} = r\,[\,(\mathrm{RY}_1 + \mathrm{RY}_2) - (\mathrm{RY}_1^2 + \mathrm{RY}_2^2) + (\tilde{a}_{12} + \tilde{a}_{21})\,\mathrm{RY}_1\mathrm{RY}_2\,].$$

Substituting $\mathrm{RY}_i = \mathrm{RYT}\,p_i$ then pulls the total out of every term—RYT from the linear ones, $\mathrm{RYT}^2$ from the quadratic ones, then

$$\frac{d\mathrm{RYT}}{dt} = r\,\mathrm{RYT} \;-\; r\,\mathrm{RYT}^2\,[\,p_1^2 + p_2^2 - (\tilde{a}_{12} + \tilde{a}_{21})\,p_1p_2\,],$$

and what is left inside the bracket depends on the composition only. Naming its inverse $\mathrm{RYT}^*(\boldsymbol{p})$ and factoring $r\,\mathrm{RYT}$, the total follows,

$$\frac{d\mathrm{RYT}}{dt} = r\,\mathrm{RYT}\left[1 - \frac{\mathrm{RYT}}{\mathrm{RYT}^*(\boldsymbol{p})}\right], \qquad \frac{1}{\mathrm{RYT}^*(\boldsymbol{p})} = p_1^2 + p_2^2 - (\tilde{a}_{12} + \tilde{a}_{21})\,p_1p_2. \tag{S31}$$

The structure is that of the logistic equation $dN/dt = rN\,(1 - N/K)$: the linear term is the growth of the pair at the common rate $r$ when both species are rare, and the quadratic term is all the density dependence the pair experiences, within and between species, pooled into one. At a fixed composition the pair therefore grows like a single logistic population whose carrying capacity, $\mathrm{RYT}^*(\boldsymbol{p})$, is the total the current mixture can support—the value a measured RYT relaxes to, at the rate $r$.

**Composition dynamics.** The composition, in turn, is obtained by differentiating $p_1 = \mathrm{RY}_1/\mathrm{RYT}$, which gives $\frac{dp_1}{dt} = p_1p_2\,(g_1 - g_2)$ with $g_i = \frac{dRY_i}{dt}/\mathrm{RY}_i$ the per capita growth rates: the share of species 1 rises exactly when it grows faster than species 2. Reading the two per capita rates off the species equations, $g_1 - g_2 = r\,[(\mathrm{RY}_2 - \mathrm{RY}_1) + \tilde{a}_{12}\,\mathrm{RY}_2 - \tilde{a}_{21}\,\mathrm{RY}_1]$, and substituting $\mathrm{RY}_i = \mathrm{RYT}\,p_i$ once more, the composition follows a replicator equation[60],

$$\frac{dp_1}{dt} = r\,\mathrm{RYT}\,p_1 p_2\,[\,p_2(1+\tilde{a}_{12}) - p_1(1+\tilde{a}_{21})\,]. \quad \text{(S32)}$$

The bracket compares the net advantage of each species and vanishes at the interior composition $p_1^*/p_2^* = (1+\tilde{a}_{12})/(1+\tilde{a}_{21})$, the equilibrium mixture of the pair; the factor RYT in front says that the composition moves faster when the pair is more abundant. Whether $\mathrm{RYT}^*(\boldsymbol{p})$ lies above or below one is decided by the quadratic form in Eq. S31. With a single interacting pair, subtracting $1 = (p_1+p_2)^2 = p_1^2 + p_2^2 + 2p_1p_2$ from it leaves a single margin,

$$\frac{1}{\mathrm{RYT}^*(\boldsymbol{p})} - 1 \;=\; -p_1p_2\,m_{12}, \qquad m_{12} = (\tilde{a}_{12}+\tilde{a}_{21}) + 2 \quad \text{(S33)}$$

The margin $m_{12}$ measures how far the pair sits from the pairwise interaction bound $\tilde{a}_{12}+\tilde{a}_{21} = -2$, at which the two species limit each other, on average, as strongly as each limits itself: it is positive when the pair competes within the bound and negative when it competes beyond it. Because $p_1p_2 > 0$ as long as both species are present, the sign of $\mathrm{RYT}^*(\boldsymbol{p}) - 1$ is the sign of $m_{12}$, whatever the composition. Therefore, $\mathrm{RYT}^*(\boldsymbol{p})$ gives us information on whether the bound on interspecific interaction of Theorem 1 is respected—or not—within the community at the transient composition $\boldsymbol{p}$.

**How a measured RYT relates to** $\mathrm{RYT}^*(\boldsymbol{p})$**.** A RYT measured along a trajectory can be read as $\mathrm{RYT}^*(\boldsymbol{p})$ only if the total reaches its settling value before the composition has had time to move. In two species this separation of time scales is exact, and it can be read off the two equations by asking how fast each coordinate returns to its resting value after a small displacement.

For RYT, differentiating Eq. S31 with respect to RYT gives $\partial_{\mathrm{RYT}}\frac{d\mathrm{RYT}}{dt} = r\,(1 - 2\,\mathrm{RYT}/\mathrm{RYT}^*)$, which at $\mathrm{RYT} = \mathrm{RYT}^*$ equals $-r$: the total returns to its ceiling at the intrinsic growth rate, whatever the interactions. This is the size mode.

For the composition, displace $p_1$ slightly from the interior point $p_1^*/p_2^* = (1+\tilde{a}_{12})/(1+\tilde{a}_{21})$ and differentiate Eq. S32 with respect to $p_1$. Because the bracket vanishes at $p_1^*$, every term in which it appears undifferentiated drops out, and only its own derivative, $-m_{12}$, survives; the prefactor $r\,\mathrm{RYT}\,p_1p_2$ is simply evaluated at the point. The rate is thus

$$-\,r\,\mathrm{RYT}^*\,p_1^*p_2^*\,m_{12} \;=\; -\,r\,(\mathrm{RYT}^* - 1),$$

the equality being Eq. S33 multiplied through by $\mathrm{RYT}^*$. This is the composition mode: the composition returns to its balanced mixture at a rate set by how far the settling total is from one. Note that nothing in this computation assumed the total to be faster than the composition; the separation is a result, not a premise.

The two rates just found are exactly the two Jacobian eigenvalues of Sec. S2.1 (Eq. S20); what the present derivation adds is which motion each one governs. Their ratio,

$$\frac{\text{composition rate}}{\text{size rate}} \;=\; |\mathrm{RYT}^* - 1|,$$

is at most one whenever $\mathrm{RYT}^* \leq 2$, as under competition: the total then leads, and leads by more the closer the settling total is to one. A measured RYT is therefore a reading of $\mathrm{RYT}^*(\boldsymbol{p})$ at the current composition.

**Species extinction.** Now consider a pair that competes beyond the bound, $m_{12} < 0$. Because $m_{12}$ does not depend on $\boldsymbol{p}$, Eq. S33 gives $\mathrm{RYT}^*(\boldsymbol{p}) < 1$ at *every* composition in which both species are present (Fig. 3B; conversely, within the bound, $\mathrm{RYT}^*(\boldsymbol{p}) > 1$ everywhere, Fig. 3A). For the RYT, Eq. S31 is a logistic law: RYT rises only while it lies below its current ceiling $\mathrm{RYT}^*(\boldsymbol{p})$, and falls otherwise. A total that starts above one therefore falls below it; and a total below one cannot climb back, since it can only ever rise toward a ceiling that itself never reaches one. Underyielding, once reached, is permanent: no shift in composition can undo it, because every composition has a ceiling below one.

For the composition, the same margin decides the fate of the balanced mixture. Its rate, the composition mode $-r\,(\mathrm{RYT}^* - 1)$, is positive when $\mathrm{RYT}^* < 1$: the interior point $\boldsymbol{p}^*$ is unstable, and the composition runs away from it toward one of the two edges, $p_1 \to 0$ or $p_1 \to 1$. (When only one of the two coefficients exceeds self-limitation in magnitude there is no interior point at all: the bracket of Eq. S32 keeps a fixed sign and the composition runs to the same edge from any start.) Either way, one species is lost.

What RYT does meanwhile closes the argument. As the composition approaches an edge, the product $p_1 p_2$ that weights the margin in Eq. S33 goes to zero, so $\mathrm{RYT}^* = 1/(1 - p_1 p_2\, m_{12})$ returns to one from below; and the effective number of species, $1/\sum_i p_i^2 = 1/(1 - 2p_1 p_2)$, returns to one as well, governed by the same product. The two events are one: the total recovers to one exactly as the pair becomes a single species at its own carrying capacity, where $\mathrm{RY} = 1$ by definition. In two species, then, the only way out of underyielding is the loss of a species—the recovery *is* the loss. We extend this result to $S$-species communities and unequal growth rates in Sec. S3.6.

### S3 Multispecies case

Here we first develop the case of equal growth rates, where any stable feasible equilibrium must overyield (Sec. S3.1). Theorem 1 then removes the assumption of equal growth rates at the price of a bound on pairwise competition, so that feasibility alone forces overyielding (Sec. S3.2). We then explain why and how this same bound on interactions turns out to govern stability (Sec. S3.3). We detail the various special cases and implications of this theorem: Why the interaction bound does not necessarily guarantee stability (Sec. S3.3.1), why certain underyielding cases are still stable (Sec. S3.3.2) and how the known result that strong facilitation destabilizes communities does not contradict our theory (Sec. S3.3.3). Once this general theory is described, we present a mean-field analytically solvable case that does not depend on the theorem but points in the same direction, demonstrating the link between overyielding and stability (Sec. S3.4). As for the two-species case, the link between complementarity, feasibility, and stability isn't only qualitative; we therefore detail the derivation of their quantitative links (Sec. S3.5). Finally, we describe the extension of these results to the non-equilibrium case, but this time in multi-species communities (Sec. S3.6).

**Table of contents of S3**


S3.1 Equal-growth-rate case: stability requires overyielding ........ S16
S3.2 Theorem on feasibility ........ S17
S3.2.1 The pairwise interaction bound ........ S17
S3.2.2 Theorem 1: feasibility implies overyielding under bounded interactions ........ S17
S3.3 Stability implications of the pairwise interaction bound ........ S19
S3.3.1 The symmetric part governs stability ........ S19
S3.3.2 Stable underyielding case ........ S20
S3.3.3 Why strong facilitative regimes are filtered ........ S22
S3.4 Solvable mean-field case for S species ........ S22
S3.5 Quantitative relations between RYT and stable coexistence ........ S23
S3.5.1 An analytical $\lambda_{\max}$–RYT bound at homogeneous growth ........ S23
S3.5.2 An analytical Ω–RYT relation in the mean-field case ........ S29
S3.6 Non-equilibrium analysis ........ S31
S3.6.1 The model in relative yields ........ S31
S3.6.2 What a relative yield total below one says about the interactions ........ S31
S3.6.3 The total settles before the composition ........ S32
S3.6.4 Unequal growth rates ........ S33
S3.6.5 The loss of a species ........ S34


##### S3.1 Equal-growth-rate case: stability requires overyielding

Let $r_i \equiv r$ and let $\boldsymbol{N}^* > 0$ be a feasible equilibrium, $\boldsymbol{A}\boldsymbol{N}^* = -\boldsymbol{K}$. The Jacobian of Eq. S3 reads $\boldsymbol{J}^* = r\,\mathrm{diag}(\mathrm{RY}_i)\,\boldsymbol{A}$. Since $\mathrm{RY}_i K_i = N_i^*$,

$$\boldsymbol{J}^*\boldsymbol{N}^* \;=\; r\,\mathrm{diag}(\mathrm{RY}_i)\,\boldsymbol{A}\boldsymbol{N}^* \;=\; -r\,\mathrm{diag}(\mathrm{RY}_i)\,\boldsymbol{K} \;=\; -r\,\boldsymbol{N}^*. \qquad \text{(S34)}$$

Moreover, one can see that the trace is set by the relative yield total. With $a_{ii} = -1$, the diagonal entries of $\boldsymbol{J}^*$ are $-r\ \mathrm{RY}_i$, so

$$\mathrm{tr}\,\boldsymbol{J}^* \;=\; -r\textstyle\sum_i \mathrm{RY}_i \;=\; -r\,\mathrm{RYT}. \qquad \text{(S35)}$$

The trace is the sum of the $S$ eigenvalues. Removing the size mode $-r$ leaves the $S-1$ remaining eigenvalues $\lambda_1, \dots, \lambda_{S-1}$—the composition modes—whose sum is real and equals

$$\textstyle\sum_{k=1}^{S-1} \lambda_k \;=\; -r\,(\mathrm{RYT} - 1). \qquad \text{(S36)}$$

Local asymptotic stability requires every eigenvalue of $\boldsymbol{J}^*$ to have a negative real part. Their sum is then negative, and by Eq. S36, RYT $> 1$.

Consequently, a feasible equilibrium with RYT $\leq 1$ carries at least one composition mode with a non-negative real part, strictly positive when RYT $< 1$: an underyielding feasible equilibrium is unstable. With the $r_i$ free, $\boldsymbol{J}^* = \mathrm{diag}(r_i \mathrm{RY}_i)\,\boldsymbol{A}$. The abundance vector is no longer an eigenvector and the trace becomes $-\sum_i r_i\,\mathrm{RY}_i$, which no longer isolates RYT. The argument does not extend to heterogeneous growth rates; yet in this case, feasibility alone can still force overyielding under a bound on pairwise competition we discuss hereafter (Theorem 1, Sec. S3.2).

#### S3.2 Theorem on feasibility

##### S3.2.1 The pairwise interaction bound

The central analytical condition of the paper concerns the symmetrized pairwise interaction sum. We say that the matrix $\boldsymbol{A}$ with $a_{ii} = -1 \forall i$ satisfies the *pairwise interaction bound* when

$$a_{ij} + a_{ji} \;>\; a_{ii} + a_{jj} = -2 \qquad \text{for every } i \neq j. \qquad \text{(S37)}$$

Condition S37 states that the reciprocal interspecific effect on the pair $(i,j)$ is weaker than the combined self-regulation of the two species—a pairwise statement of niche differentiation[13].

Because facilitative coefficients are non-negative in our sign convention, any *symmetrically* facilitative pair automatically satisfies the bound: $a_{ij} + a_{ji} \geq 0 > -2$. Where the bound can fail is in two regimes. First, symmetric mutual competition that is strong enough on both sides: $a_{ij}, a_{ji} < 0$ with $a_{ij} + a_{ji} \leq -2$. Second, strongly asymmetric mixed-sign pairs in which a directional antagonism in one direction is not compensated by a facilitative effect of comparable magnitude in the other: such pairs satisfy $a_{ij} + a_{ji} \leq -2$ even though one of the coefficients is positive. The bound therefore restricts *both* the strength of mutual competition and the magnitude of strong directional antagonisms, while allowing arbitrary asymmetry $a_{ij} \neq a_{ji}$ and arbitrary mixtures of facilitative and competitive entries within these limits.

##### S3.2.2 Theorem 1: feasibility implies overyielding under bounded interactions

This section states and proves the central analytical result of the manuscript: under the pairwise interaction bound of Eq. S37, any feasible equilibrium of the gLV system with positive monoculture carrying capacities satisfies RYT $> 1$. The result holds for arbitrary species number, arbitrary growth-rate heterogeneity, and arbitrary asymmetry of the interaction matrix within the bound.

**Theorem 1 (Feasibility implies overyielding).** *Let $\boldsymbol{A} = (a_{ij})$ be an $S \times S$ matrix with $a_{ii} = -1\,\forall i$ and let $\boldsymbol{K} = (K_1 > 0, \dots, K_S > 0)^\top$. Let $\boldsymbol{N}^*$ be the equilibrium of the gLV system Eq. S1 solving $\boldsymbol{AN}^* = -\boldsymbol{K}$. If $\boldsymbol{N}^*$ is feasible, then the interaction bound (Eq. S37) implies overyielding:*

$$\begin{cases} a_{ij} + a_{ji} > -2 \quad \text{for all } i \neq j, \\ N_i^* > 0 \quad \text{for all } i \end{cases} \Rightarrow \text{ RYT } > 1. \tag{S38}$$

This result holds for any number of species $S$ and any heterogeneity in growth rates. The proof requires neither symmetry of $\boldsymbol{A}$, nor any specific sign structure beyond the pairwise bound. Growth rates $r_i$ play no role: they enter the temporal dynamics (Eq. S1) but not the equilibrium relation $\boldsymbol{AN}^* = -\boldsymbol{K}$ on which the theorem depends. As a consequence, Theorem 1 applies to arbitrarily heterogeneous communities of competitors, facilitators, and mixed-sign interaction types, provided that every pair $(i, j)$ satisfies $a_{ij} + a_{ji} > -2$. As a corollary, if RYT $\leq 1$ at a feasible equilibrium of the gLV system with $\boldsymbol{K} > 0$, then there exists at least one pair $i \neq j$ such that $a_{ij} + a_{ji} \leq -2$.

*Proof.* Let $\boldsymbol{N}^* > 0$ and define the normalized abundance vector $\boldsymbol{n} = \boldsymbol{N}^* / \sum_j N_j^*$, so $n_i > 0$ for every $i$ and $\sum_i n_i = 1$. The relative yield total is homogeneous of degree zero in $\boldsymbol{N}^*$: from $\boldsymbol{AN}^* = -\boldsymbol{K}$, we have for each species $K_i = -(\boldsymbol{AN}^*)_i$, hence

$$\text{RY}_i = \frac{N_i^*}{K_i} = \frac{N_i^*}{-(\boldsymbol{AN}^*)_i} = \frac{n_i}{-(\boldsymbol{An})_i}.$$

Summing across species yields the expression

$$\text{RYT} = \sum_{i=1}^{S} \frac{n_i}{-(\boldsymbol{An})_i}. \tag{S39}$$

*Positivity of $-(\boldsymbol{An})_i$.* Before applying the Engel–Titu inequality, we verify that $-(\boldsymbol{An})_i > 0$ for every $i$. Since $\boldsymbol{AN}^* = -\boldsymbol{K}$ and $\boldsymbol{K} > 0$, we have $(\boldsymbol{AN}^*)_i = -K_i < 0$, and dividing by $\sum_j N_j^* > 0$ gives $-(\boldsymbol{An})_i > 0$.

*Step 1: lower bound by Engel–Titu.* We rewrite each term of RYT as a ratio of squares,

$$\text{RYT} = \sum_{i=1}^{S} \frac{n_i^2}{-n_i\,(\boldsymbol{An})_i}.$$

The Engel–Titu lemma (equivalent to the Cauchy–Schwarz inequality applied to fractions[61]) states that, for any real numbers $x_1, \dots, x_S$ and any strictly positive $y_1, \dots, y_S$,

$$\sum_{i=1}^{S} \frac{x_i^2}{y_i} \geq \frac{(\sum_{i=1}^{S} x_i)^2}{\sum_{i=1}^{S} y_i}. \tag{S40}$$

Applying the lemma with $x_i = n_i$ and $y_i = -n_i(\boldsymbol{An})_i > 0$ gives

$$\text{RYT} \geq \frac{(\sum_i n_i)^2}{\sum_i (-n_i(\boldsymbol{An})_i)} = \frac{1}{-\boldsymbol{n}^\top \boldsymbol{A}\,\boldsymbol{n}}, \tag{S41}$$

since $\sum_i n_i = 1$ and $\sum_i (-n_i(\boldsymbol{An})_i) = -\boldsymbol{n}^\top \boldsymbol{An}$.

*Step 2: bounding the denominator*. We expand $-\boldsymbol{n}^\top \boldsymbol{An}$ using $a_{ii} = -1$ and $\sum_i n_i = 1$:

$$-\boldsymbol{n}^\top \boldsymbol{An} \;=\; \sum_{i=1}^{S} n_i^2 \;-\; \sum_{i<j} (\,a_{ij} + a_{ji})\, n_i n_j.$$

By the pairwise interaction bound $a_{ij} + a_{ji} > -2$ and the strict positivity of $n_i n_j$ for every $i \neq j$, we have

$$-\sum_{i<j} (\,a_{ij} + a_{ji})\, n_i n_j \;<\; 2\sum_{i<j} n_i\, n_j,$$

and therefore

$$-\boldsymbol{n}^\top \boldsymbol{An} \;<\; \sum_i n_i^2 + 2\sum_{i<j} n_i\, n_j \;=\; (\sum_i n_i)^2 \;=\; 1.$$

Combining with Eq. S41 gives RYT $\geq 1/(-\boldsymbol{n}^\top \boldsymbol{An}) > 1$. □

##### S3.3 Stability implications of the pairwise interaction bound

Theorem 1 connects overyielding and feasibility, using a bound on pairwise competition. Here we show that the same pairwise bound also governs local stability.

Finally, we showcase unlikely counter-examples of feasible, *stable underyielding* states—which are impossible if species growth rates are equal and require at least three species.

###### S3.3.1 The symmetric part governs stability

Local stability is determined by both the feasible equilibrium and the interaction values, which are combined in the Jacobian matrix of the community[62,63]. So far we have focused on the interaction matrix only, with the aim of determining sufficient conditions granting local stability of any associated Jacobian matrix evaluated at a feasible equilibrium. Among these conditions, $\boldsymbol{A}$ may be negative definite, Volterra-dissipative, or D-stable. Negative definite is the strongest condition, which implies Volterra-dissipative and, in turn, D-stable, but it is also the easiest to mathematically verify. Negative-definite is defined as all eigenvalues of the symmetric part $\boldsymbol{A}_s$ are negative; with the symmetric part computed as $\boldsymbol{A}_s = 1/2\,(\boldsymbol{A} + \boldsymbol{A}^\top)$. A negative definite matrix has all its principal submatrices negative definite, in particular every $2 \times 2$ submatrix of $\boldsymbol{A}_s$. That is, the eigenvalues of

$$\begin{bmatrix} -1 & \frac{1}{2}(a_{ij} + a_{ji}) \\ \frac{1}{2}(a_{ij} + a_{ji}) & -1 \end{bmatrix} \tag{S42}$$

must be negative. This condition holds if and only if $-1 < \frac{1}{2}(a_{ij} + a_{ji}) < 1$. The pairwise interaction bound is thus the two-species shadow of the true condition $\boldsymbol{A}_s \prec 0$. Its lower half, $a_{ij} + a_{ji} > -2$, is our bound and caps mutual *competition*; its upper half, $a_{ij} + a_{ji} < 2$, caps mutual *facilitation* and is irrelevant here, because the requirement $K_i > 0$ already filters out too strongly facilitative pairs (Sec. S3.3.3). The interaction bound of Theorem 1 is therefore the competitive half of negative definiteness.

The bound is necessary for $\boldsymbol{A}_s \prec 0$, not sufficient. At $S = 2$ the two coincide, so the bound alone guarantees stability. For $S \geq 3$ it does not: full negative definiteness constrains whole rows of $\boldsymbol{A}_s$, not isolated pairs, and the gap between the pairwise condition and the full one is where the exceptions live.

**Bounded yet unstable.** The symmetric matrix

$$\boldsymbol{A} = \begin{pmatrix} -1 & -0.9 & -0.9 \\ -0.9 & -1 & 0.9 \\ -0.9 & 0.9 & -1 \end{pmatrix}, \qquad \boldsymbol{N}^* = (1,\, 0.1,\, 0.1)^\top > 0, \tag{S43}$$

has every pairwise sum inside the bound ($-1.8$ or $+1.8$) and a feasible, overyielding equilibrium (RYT $\approx 1.07$, consistent with Theorem 1), yet one positive Jacobian eigenvalue: species 2 and 3 facilitate each other and rise together while their shared competitor, species 1, falls, closing a positive feedback loop—a facilitative coalition collectively excluding a competitor, which no single coefficient can diagnose. The same escape occurs in a purely competitive community. The symmetric matrix

$$\boldsymbol{A} = \begin{pmatrix} -1 & 0 & -0.9 & -0.9 \\ 0 & -1 & -0.9 & -0.9 \\ -0.9 & -0.9 & -1 & 0 \\ -0.9 & -0.9 & 0 & -1 \end{pmatrix}, \qquad \boldsymbol{K} = \boldsymbol{1}, \tag{S44}$$

has every pairwise sum ($-1.8$ or $0$) inside the bound, yet a feasible, overyielding equilibrium (RYT $\approx 1.43$) carrying a positive Jacobian eigenvalue ($\lambda_{\max}(\boldsymbol{J}^*) = +0.286$); being symmetric, it cannot be stabilized by any growth rates (Sylvester's law of inertia[64]).

**Violating the bound destabilizes, and overwhelmingly.** Crossing the interaction bound on pairwise interaction is the opposite case: strong mutual competition ($a_{ij} + a_{ji} < -2$) makes the $2 \times 2$ block of $\boldsymbol{A}_s$ indefinite directly, a *localized* destabilizing direction. In the simulated ensemble ($S = 10$, feasible; Sec. S4, Fig. 2) stability is predominant among bound-respecting communities and almost absent among violators: about $76\%$ of feasible bound-respecting communities are stable, against less than 1% of bound-violating ones; and roughly $94\%$ of unstable feasible communities violate the bound. Most violators also lose overyielding (RYT $< 1$), but a residual fraction remain overyielding yet unstable.

##### S3.3.2 Stable underyielding case

Stable underyielding cases must have uneven growth rates (Sec. S3.1) and violate the bound on interaction (Sec. S3.2). But not every heterogeneity in growth rates can stabilize communities; it requires a fine-tuned hierarchy of species time scales (see Fig. S3).

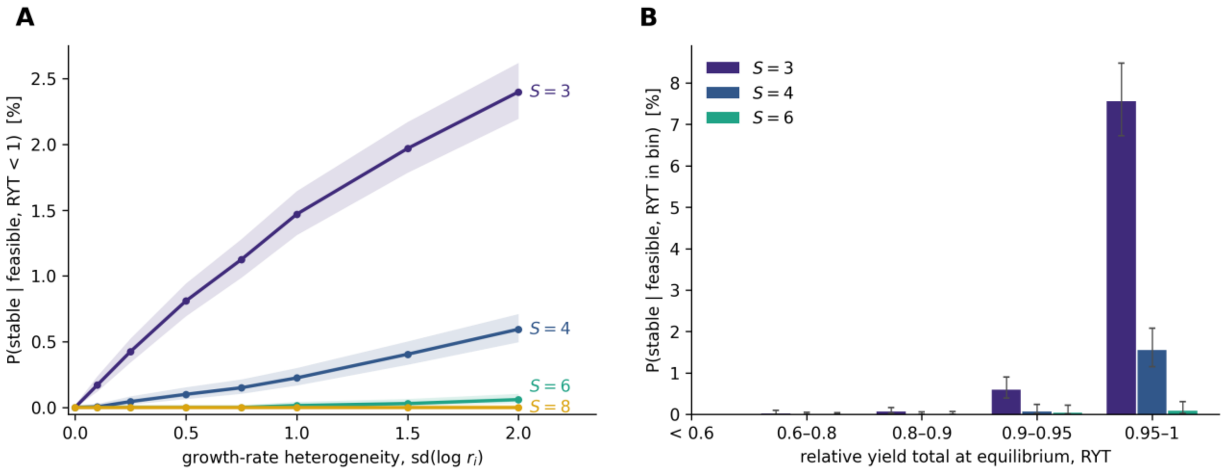


**Fig. S3. Stable underyielding requires widely spread growth rates and marginal underyielding, and remains rare under both.** Feasible underyielding equilibria ($\boldsymbol{N}^* > 0$, RYT $< 1$; ensemble in Sec. S4.1). **(A)** Probability of local stability against the growth-rate heterogeneity $\mathrm{sd}(\log r_i)$, for different numbers of species (bands: 95% Wilson intervals). **(B)** Same probability at $\mathrm{sd}(\log r_i) = 1$ by bins of RYT (bars: 95% Wilson intervals; bins with fewer than 30 communities omitted): rescue is confined almost entirely to equilibria close to RYT $= 1$, and vanishes as richness grows.

The mechanism is easy to see in the 3 species case: a pair of species that compete too strongly to coexist on their own, and a third species that grows much faster than both. Take species 1 and 2 with strong interspecific interaction $a_{12}a_{21} > 1$. On their own, the pair has a feasible equilibrium (both $1 + a_{12}$ and $1 + a_{21}$ are negative), but it is a saddle—the two-species Jacobian has determinant $r_1 r_2 \mathrm{RY}_1 \mathrm{RY}_2 (1 - a_{12}a_{21}) < 0$—so whichever species establishes first excludes the other. Now add species 3, linked to each of the two by weaker interactions ($a_{13}a_{31} < 1$ and $a_{23}a_{32} < 1$), and let $r_3 \gg r_1, r_2$. Because species 3 is fast, it reaches its own zero-growth condition long before the pair has moved: at every moment, $N_3 \approx K_3 + a_{31}N_1 + a_{32}N_2$. Substituting this into the equations of species 1 and 2 (Eq. S1) removes species 3 from the dynamics and leaves a two-species system in which every parameter of the pair has been altered by its passage through species 3:

$$\frac{dN_1}{dt} = r_1 N_1 \left[\left(1 + a_{13}\frac{K_3}{K_1}\right) - (1 - a_{13}a_{31})\frac{N_1}{K_1} + (a_{12} + a_{13}a_{32})\frac{N_2}{K_1}\right], \tag{S45}$$

and symmetrically for species 2. The new effective interaction coefficients are now

$$a_{12}^{\mathrm{eff}} = \frac{a_{12} + a_{13}a_{32}}{1 - a_{13}a_{31}}, \qquad a_{21}^{\mathrm{eff}} = \frac{a_{21} + a_{23}a_{31}}{1 - a_{23}a_{32}}.$$

The pair coexists stably when $a_{12}^{\mathrm{eff}} a_{21}^{\mathrm{eff}} < 1$: the indirect paths through species 3 must soften the pair's mutual competition enough for the effective pair to fall back within the coexistence condition that the direct pair violated. Written out, this condition is $\det \boldsymbol{A} < 0$; it involves the interactions alone and is in any case necessary for stability whatever the growth rates. Growth rates therefore do not change *which* interaction matrices can be stabilized; what they change is whether species 3 is fast enough for the reduction to Eq. S45 to hold. At equal growth rates the condition cannot be met for an underyielding community, as shown above.

Considering the relative yields, they are set by $\boldsymbol{A}\boldsymbol{N}^* = -\boldsymbol{K}$ and do not depend on growth rates at all, so the community underyields exactly as it would at equal rates; only its stability has changed, and it has changed through the ordering of time scales.

##### S3.3.3 Why strong facilitative regimes are filtered

Random-matrix and mutualism-dynamics studies establish that strong positive interactions destabilize large communities[65–67]. The common message is that strong facilitation, on its own, can destabilize large communities.

Our framework does not contradict this. The normalization $a_{ii} = -1$ with $K_i > 0$ enforces strict self-regulation at every node and the boundary $a_{ij} + a_{ji} = -2$ is exactly where that self-regulation is not enough to stabilize the pair of species; the threshold beyond which their destabilizing competitive regime begins. Strong facilitation is ruled out by a separate route: requiring $\boldsymbol{K} = -\boldsymbol{A}\boldsymbol{N}^* > 0$ turns some $K_i$ non-positive once mutual facilitation is too strong. As we also require $K_i > 0$, it removes these overly mutualistic configurations before they can destabilize.

#### S3.4 Solvable mean-field case for S species

Let every pair interact with the same coefficient, $a_{ij} = \rho$ for all $i \neq j$, with $a_{ii} = -1$ as throughout: $\rho < 0$ is mean-field competition and $\rho > 0$ mean-field facilitation. The interaction matrix is then $\boldsymbol{A} = -(1+\rho)\,\boldsymbol{I} + \rho\,\boldsymbol{J}$, with $\boldsymbol{J}$ the all-ones matrix, and its spectrum is explicit: the eigenvalue $-[1-(S-1)\rho]$ on the eigenvector $(1,\dots,1)^\top$ (the size mode) and the eigenvalue $-(1+\rho)$ with multiplicity $S-1$ on the subspace orthogonal to it (the composition modes). Both are negative if and only if

$$-1 \;<\; \rho \;<\; \frac{1}{S-1}, \tag{S46}$$

and in that range $\boldsymbol{A} = \boldsymbol{A}_s \prec 0$: every feasible equilibrium is locally asymptotically stable, whatever the growth rates and carrying capacities (Sec. S3.3.1). Which mode is the slower one depends on the sign of $\rho$: the composition eigenvalue $-(1+\rho)$ leads under competition ($\rho < 0$) and reaches 0 exactly as $\rho \to -1$, i.e., as $a_{ij} + a_{ji} \to -2$, the pairwise interaction bound; the size eigenvalue $-[1-(S-1)\rho]$ leads under facilitation ($\rho > 0$) and reaches 0 as $\rho \to 1/(S-1)$.

The equilibrium relation $\boldsymbol{A}\boldsymbol{N}^* = -\boldsymbol{K}$, that is $[(1+\rho)\,\boldsymbol{I} - \rho\,\boldsymbol{J}]\boldsymbol{N}^* = \boldsymbol{K}$, is solved explicitly. With $K_{\text{tot}} = \sum_i K_i$,

$$N_i^* \;=\; \frac{K_i[1-(S-1)\rho] + \rho\, K_{\text{tot}}}{(1+\rho)[1-(S-1)\rho]}. \tag{S47}$$

Writing $\pi_i = K_i / K_{\text{tot}}$ for the share of each carrying capacity, the community is feasible when every share exceeds a common threshold,

$$N_i^* > 0 \;\text{ for all } i \quad \Leftrightarrow \quad \pi_i \;>\; \frac{-\rho}{1-(S-1)\rho} \;\text{ for all } i, \tag{S48}$$

and the relative yield total reads

$$\text{RYT} \;=\; \frac{S}{1+\rho} \;+\; \frac{\rho}{(1+\rho)[1-(S-1)\rho]} \sum_{i=1}^{S} \frac{1}{\pi_i}. \tag{S49}$$

At equal carrying capacities ($\pi_i = 1/S$, so $\sum_i 1/\pi_i = S^2$) this collapses to

$$\text{RYT} = \frac{S}{1-(S-1)\rho}, \tag{S50}$$

which exceeds one exactly when $\rho > -1$. Everything else in Eq. S49 turns on the sign of $\rho$, which is the sign of the coefficient of $\sum_i 1/\pi_i$.

When $\rho < 0$ (competition), the feasibility threshold in Eq. S48 is positive and restricts the carrying capacities to a polytope; the coefficient of $\sum_i 1/\pi_i$ is negative, so RYT is largest at equal capacities (Eq. S50, by Cauchy–Schwarz, $\sum_i 1/\pi_i \geq S^2$) and smallest where $\sum_i 1/\pi_i$ is largest. Being strictly convex, $\sum_i 1/\pi_i$ attains its maximum at a vertex of the polytope, where $S-1$ species sit exactly at the threshold: there their equilibrium abundances vanish, the last species stands alone at $N_S^* = K_S$, and $\text{RYT} = 1$ by inspection. Hence, under mean-field competition,

$$1 < \text{RYT} \leq \frac{S}{1-(S-1)\rho} \qquad (-1 < \rho < 0), \tag{S51}$$

the lower bound being approached only as the community collapses to a monoculture—in agreement with Theorem 1. Beyond the bound, $\rho < -1$, feasible equilibria still exist but $\text{RYT} < 1$ at equal capacities and the composition eigenvalue is positive: the community underyields and is unstable, the regime read along trajectories in Sec. S3.6.3.

When $\rho > 0$ (facilitation), the threshold in Eq. S48 is negative, so feasibility is automatic for any $K_i > 0$; and the coefficient of $\sum_i 1/\pi_i$ is now positive, so the same Cauchy–Schwarz inequality runs the other way and equal capacities give the *minimum*:

$$\text{RYT} \geq \frac{S}{1-(S-1)\rho} > S > 1 \qquad (0 < \rho < 1/(S-1)). \tag{S52}$$

Facilitation thus produces an overyielding that grows at least linearly with richness, a much stronger guarantee than the competitive case, whose lower bound does not scale with $S$. Above $\rho = 1/(S-1)$ no feasible equilibrium exists for $\boldsymbol{K} > 0$: this is the feasibility filter of Sec. S3.3.3, which removes too strongly facilitative communities before the size mode can destabilize them. In the whole range of Eq. S46, then, feasibility, local stability, and overyielding hold jointly.

#### S3.5 Quantitative relations between RYT and stable coexistence

##### S3.5.1 An analytical $\lambda_{\max}$–RYT bound at homogeneous growth

The numerical $\lambda_{\max}$–RYT relation of Fig. 2A (see also Figs. S4–S5) admits an analytical envelope when growth rates are common, $r_i \equiv r$. As established in Sec. S3.1, the equilibrium abundance vector $\boldsymbol{N}^*$ is a Jacobian eigenvector, with eigenvalue $-r$ (the size mode), and the remaining $S-1$ eigenvalues (the composition modes) have real parts summing to

$$\sum_{k=1}^{S-1} \text{Re}\,\lambda_k = -r\,(\text{RYT}-1). \tag{S53}$$

Because the maximum of a set of reals is at least their mean, the leading composition eigenvalue—and hence $\lambda_{\max}$ whenever it exceeds the size mode $-r$—obeys

$$\lambda_{\max} \geq -\frac{r\,(\text{RYT}-1)}{S-1} \qquad (r_i \equiv r), \tag{S54}$$

with equality when the composition eigenvalues share a common real part. The bound is an *upper limit on stability*: a community cannot be more stable than $-r(\mathrm{RYT}-1)/(S-1)$, so damping improves only as RYT grows, and underyielding ($\mathrm{RYT} < 1$) forces the right-hand side positive—at least one composition mode is unstable. At $S = 2$ the single composition mode saturates the bound and Eq. S54 becomes the exact two-species identity $\lambda_{\max} = -r(\mathrm{RYT}-1)$ of Eq. S21. For $S > 2$ the $1/(S-1)$ factor is the analytical counterpart of the softening of the $\lambda_{\max}$–RYT correlation with richness visible in Fig. S5: the $S-1$ composition modes share the same budget $-r(\mathrm{RYT}-1)$, so the leading one is pulled toward zero as $S$ grows. Heterogeneous growth rates loosen the bound—by redistributing real parts across modes they can stabilize an underyielding community, the mechanism characterized in Sec. S3.3.2.

666

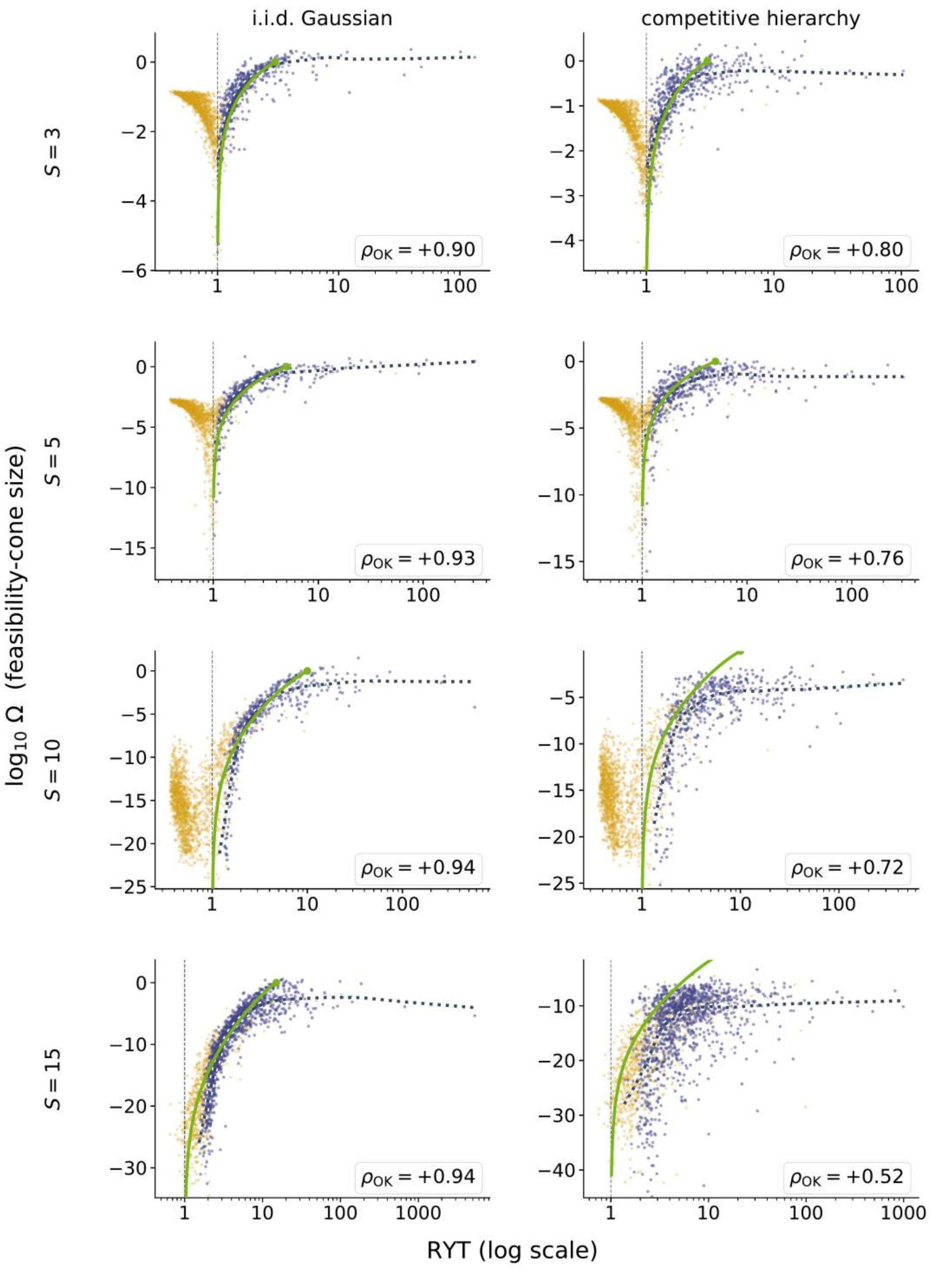

i.i.d. Gaussian
competitive hierarchy
S = 3
S = 5
S = 10
S = 15
log10 Ω (feasibility-cone size)
RYT (log scale)
ρOK = +0.90
ρOK = +0.80
ρOK = +0.93
ρOK = +0.76
ρOK = +0.94
ρOK = +0.72
ρOK = +0.94
ρOK = +0.52

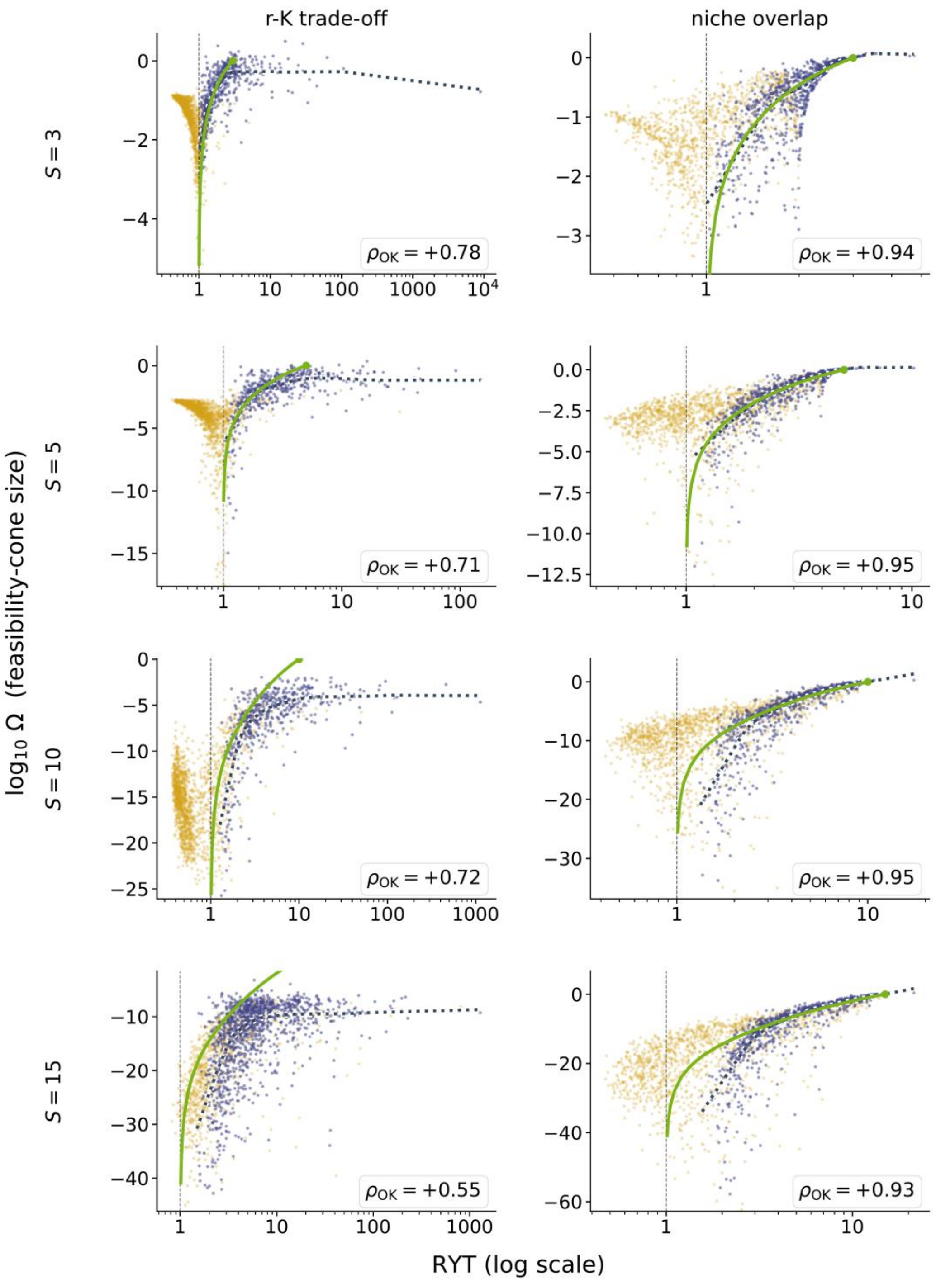


**Fig. S4. Robustness of the** Ω–RYT **relation across species number and interaction structure.** Each panel shows the feasibility-domain size $\log_{10}\Omega$ against RYT (log scale) for one combination of species number $S$ (rows: $S = 3{,}5{,}10{,}15$) and interaction structure (columns: i.i.d. Gaussian and competitive hierarchy on the first page, $r$–$K$ trade-off and niche overlap on the second). Purple points satisfy the pairwise bound $a_{ij} + a_{ji} > -2$; yellow points violate it. The LOWESS smoother (dark, dotted) and the Spearman correlation $\rho_{\mathrm{OK}}$ are computed on the bound-satisfied subset, matching the Ω–RYT panel of Fig. 2. The green line is the mean-field relation (Sec. S3.5.2). The positive Ω–RYT relation holds across all sixteen combinations.

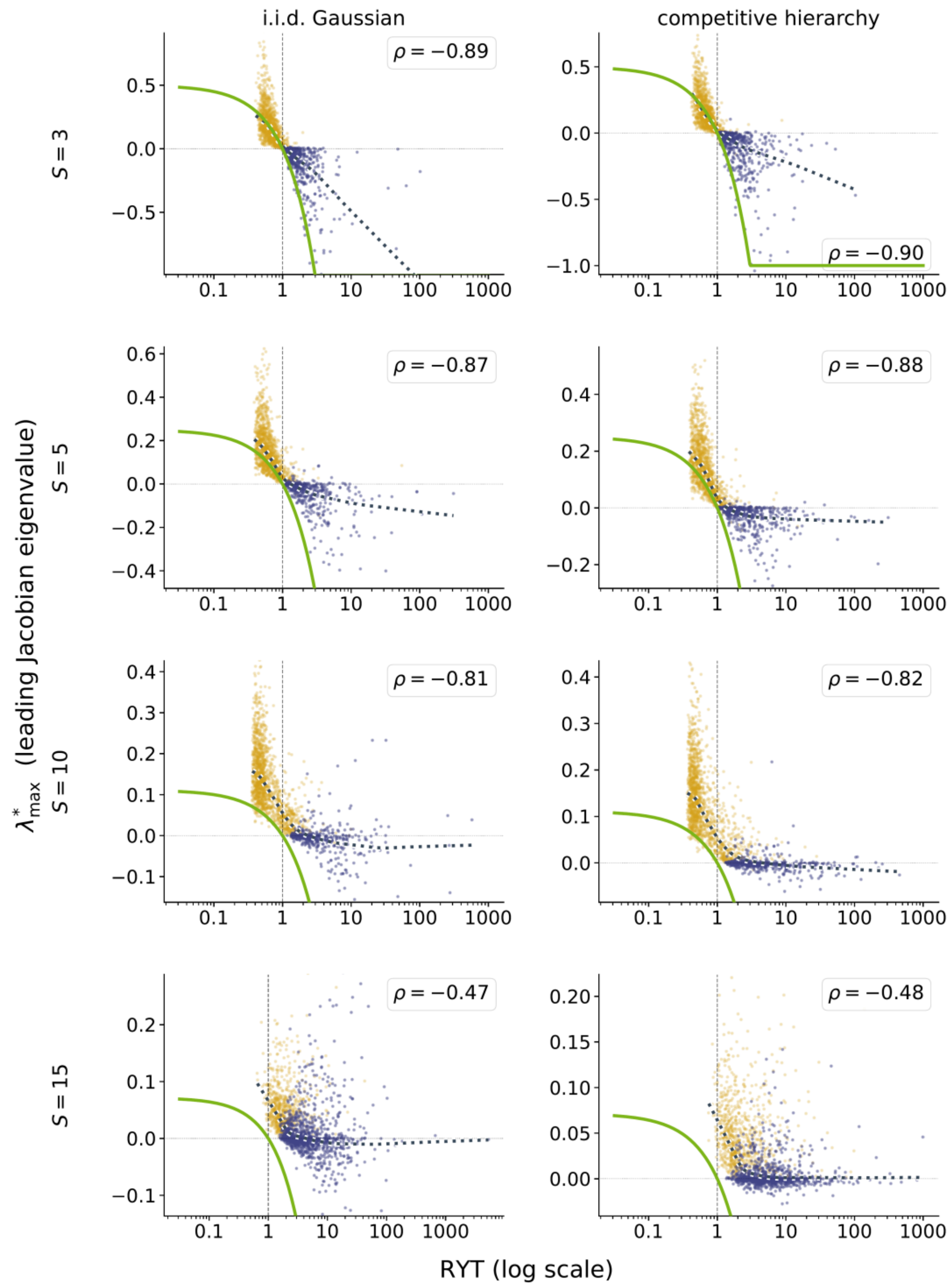
i.i.d. Gaussian
competitive hierarchy
S = 3
S = 5
S = 10
S = 15
ρ = −0.89
ρ = −0.90
ρ = −0.87
ρ = −0.88
ρ = −0.81
ρ = −0.82
ρ = −0.47
ρ = −0.48
λ*max (leading Jacobian eigenvalue)
RYT (log scale)

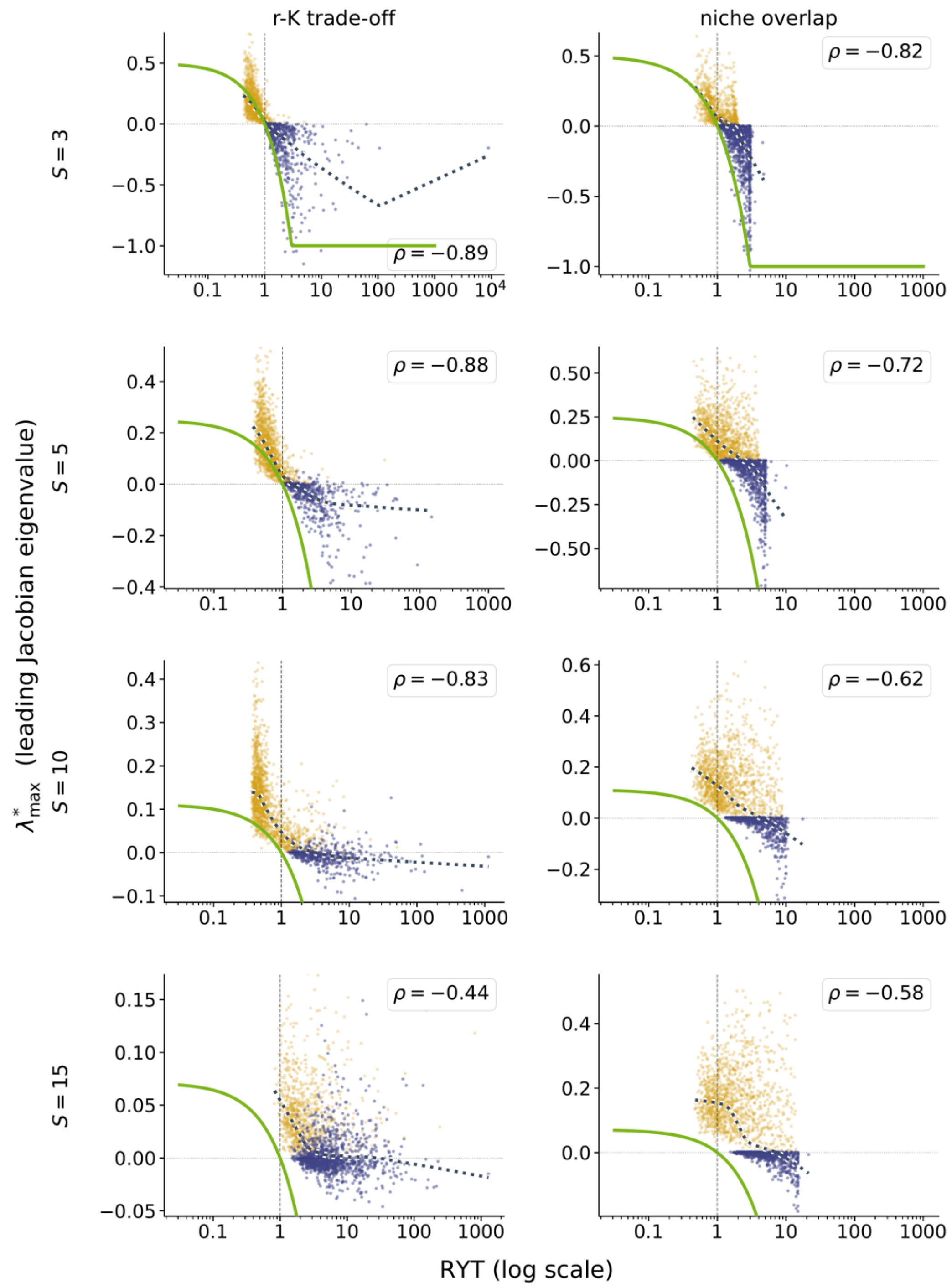


**Fig. S5. Robustness of the $\lambda_{\max}$–RYT relation across species number and interaction structure.** Each panel shows the leading real part of the Jacobian spectrum $\lambda_{\max}$ against RYT (log scale) for one combination of species number $S$ (rows) and interaction structure (columns, as in Fig. S4). Purple points satisfy the pairwise bound $a_{ij} + a_{ji} > -2$; yellow points violate it. The LOWESS smoother (dark, dotted) and the Spearman correlation $\rho$ are computed on the full cloud, matching the $\lambda_{\max}$–RYT panel of Fig. 2. The green line is the analytical bound of Eq. S54. The negative $\lambda_{\max}$–RYT relation holds across all sixteen combinations, and underyielding communities are almost always unstable ($\lambda_{\max} > 0$ for at least 99.5% of those with RYT $< 1$ in every panel).

##### S3.5.2 An analytical Ω–RYT relation in the mean-field case

The path to relate Ω to RYT is similar to the two-dimensional case (Sec. S2.3), but more complicated regarding Ω computation. We start by finding how $\rho$ relates to the maximum RYT. From Sec. S3.4, the maximum total relative yield, in the competition case ($-1 < \rho < 0$) is given at distributed carrying capacities $\pi = 1/S$. This results in

$$\mathrm{RYT}_{\mathrm{max}} = \frac{1}{1+\rho}\left[S + \frac{\rho}{1-(S-1)\rho}\sum_{i=1}^{S} S\right] = \frac{S}{1-(S-1)\rho} \rightarrow \rho = \frac{\mathrm{RYT}_{\mathrm{max}}-S}{(S-1)\cdot\mathrm{RYT}_{\mathrm{max}}} \quad \text{(S55)}$$

The computation of Ω needs several steps. First, we compute the pairwise angles between any two columns of $-\boldsymbol{A}$. As we are in the mean-field case, these angles are all equal, and their cosine is given by

$$\cos\theta = \frac{\boldsymbol{v}_1 \cdot \boldsymbol{v}_2}{||\boldsymbol{v}_1||||\boldsymbol{v}_2||} = \frac{-2\rho+(S-2)\rho^2}{1+(S-1)\rho^2} \quad \text{(S56)}$$

At this stage, it is already interesting to write $\theta$ as a function of $\mathrm{RYT}_{\mathrm{max}}$. This is given by

$$\theta = \cos^{-1}\left(\frac{(S-\mathrm{RYT}_{\mathrm{max}})\cdot(\mathrm{RYT}_{\mathrm{max}}+S-2)}{(S-1)[(\mathrm{RYT}_{\mathrm{max}}-1)^2+S-1]}\right) \quad \text{(S57)}$$

Fig. S6B shows that $\theta$ increases with $\mathrm{RYT}_{\mathrm{max}}$; since a larger $\theta$ yields a larger Ω, Ω also increases with $\mathrm{RYT}_{\mathrm{max}}$.

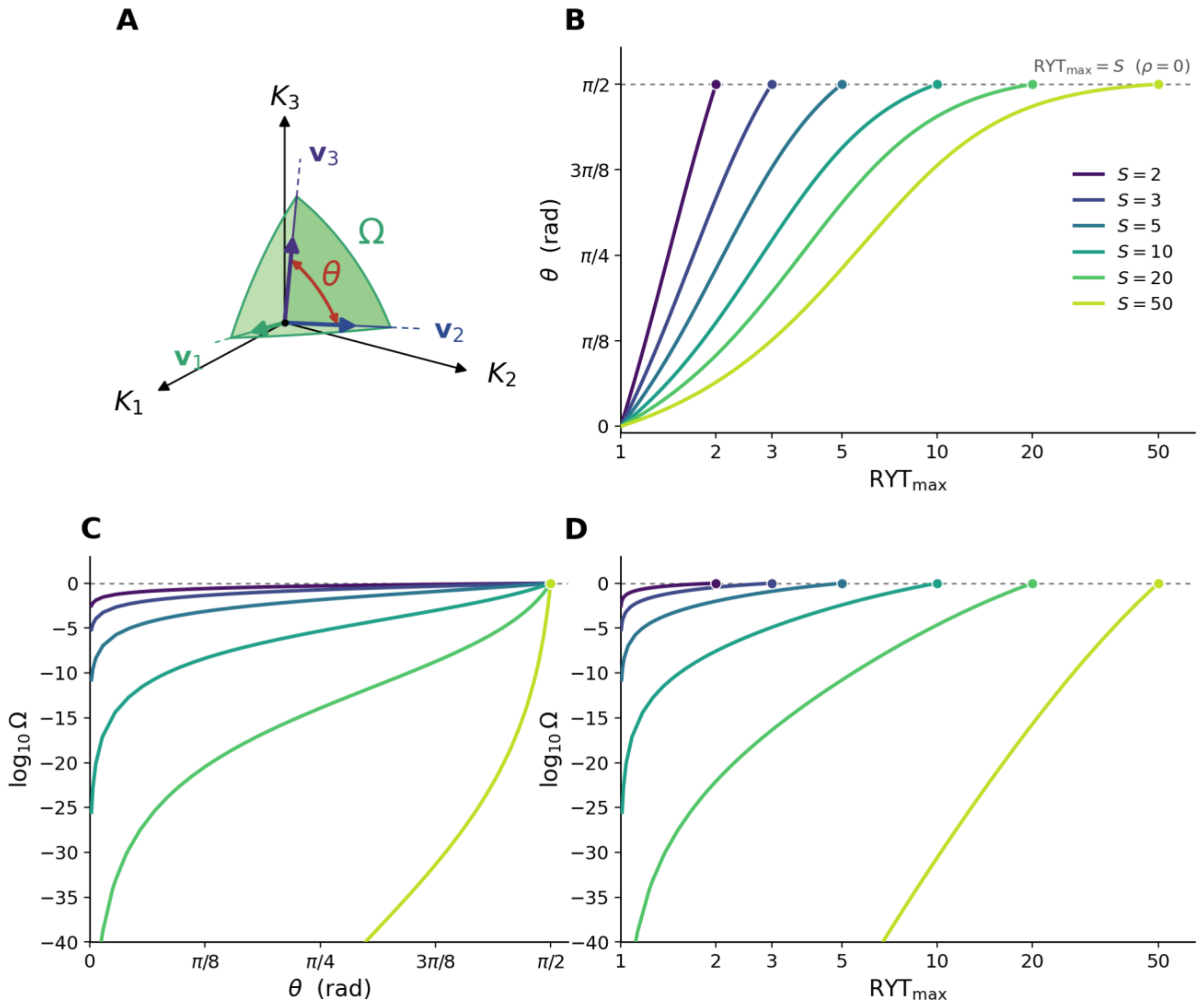


**Fig. S6. Edge angle $\theta$ of the feasibility cone and its relation to $\mathrm{RYT_{max}}$ in the mean-field case.** (**A**) Feasibility domain of a three-species community in the space of carrying capacities $(K_1, K_2, K_3)$: the cone generated by the vectors $\boldsymbol{v}_i$, whose opening is measured by the normalized solid angle $\Omega$. $\theta$ is the angle between two edges of the cone (here $\boldsymbol{v}_2$ and $\boldsymbol{v}_3$); in the mean-field case all pairwise angles are equal. (**B**) Analytical relation between $\theta$ and $\mathrm{RYT_{max}}$ (Eq. S57) for $S = 2$ to 50 ($\mathrm{RYT_{max}}$ on a log scale). $\theta$ increases monotonically with $\mathrm{RYT_{max}}$, from $\theta = 0$ at the pairwise competitive bound ($\rho = -1$, $\mathrm{RYT_{max}} = 1$) to $\theta = \pi/2$ in the absence of interaction ($\rho = 0$, $\mathrm{RYT_{max}} = S$, dashed line). (**C**) Analytical relation between $\theta$ and $\Omega$ (log10), for $S = 2$ to 50 (Eq. S58–S59). (**D**) Analytical relation between $\mathrm{RYT_{max}}$ and $\Omega$ (log10), for $S = 2$ to 50 (Eq. S57–S59).

In the mean-field case, we can go a step further and relate $\theta$ to $\Omega$ through an integral for any number of species $S > 1$:

$$\Omega = 2^S \int_{-\infty}^{+\infty} \Phi'(t)\Phi\left(t \cdot \sqrt{\frac{c}{1-c}}\right)^S dt \qquad \text{(S58)}$$

where $\Phi$ is the cumulative distribution function of the normal distribution of mean 0 and standard deviation 1 ($\Phi'$ its derivative, i.e., the density function), and $c$ given by

$$c = \frac{-\cos(\theta)}{1+(S-2)\cdot\cos(\theta)} \qquad \text{(S59)}$$

The proof of the formula is as follows. First, we note that, in Eq. S14 the term $\frac{|\det \boldsymbol{A}|}{\pi^{S/2}} \int \cdots \int_{\mathbb{R}^S_{>0}} e^{-\boldsymbol{x}^T\boldsymbol{A}^T\boldsymbol{A}\boldsymbol{x}}\, d\boldsymbol{x}$, that is $\Omega/2^S$, represents the probability, if the vector $\boldsymbol{x}$ is sampled uniformly on the sphere in dimension $S$, to have $-\boldsymbol{A}^{-1}\boldsymbol{x} > 0$. In ecological terms, it is the probability of sampling a vector of carrying capacities ($\boldsymbol{x} = \boldsymbol{K}$) such that the corresponding equilibrium $\boldsymbol{N}^* = -\boldsymbol{A}^{-1}\boldsymbol{K} > 0$ is feasible, i.e., $P(-\boldsymbol{A}^{-1}\boldsymbol{K} > 0)$ assuming $\boldsymbol{K}$ to be sampled uniformly on the sphere. This is equivalent to the probability of a feasible equilibrium $P(\boldsymbol{N}^* > 0)$, but $\boldsymbol{N}^*$ is sampled from a multivariate normal distribution of mean $\boldsymbol{0}$ and variance-covariance $(\boldsymbol{A}^T\boldsymbol{A})^{-1}$. Under the mean-field assumption, this is an equicorrelated distribution of correlation given by the term $c$. For such a distribution, the probability $P(\boldsymbol{N}^* > 0)$ is given by the integration term of Eq. S58 (68).

##### S3.6 Non-equilibrium analysis

This section shows that a relative yield total below one, read at any moment of a community's dynamics, still reports on the interactions among its species—and that, outside one exceptional case we identify, it cannot be undone without the loss of a species. Everything rests on rewriting the model in relative yields and reading a single quadratic form; the reasoning behind Fig. 3 follows from it.

###### S3.6.1 The model in relative yields

The relative yield of species $i$, $\mathrm{RY}_i = N_i/K_i$ (Eq. S4), was defined at equilibrium; we now read it at every moment. Dividing Eq. S1 by $K_i$,

$$\frac{d\mathrm{RY}_i}{dt} = r_i\, \mathrm{RY}_i\Big(1 + \textstyle\sum_{j=1}^{S} \tilde{a}_{ij}\, \mathrm{RY}_j\Big), \qquad \tilde{a}_{ij} = a_{ij}\frac{K_j}{K_i}, \qquad \tilde{a}_{ii} = -1. \tag{S60}$$

In relative yields the system is the same generalized Lotka–Volterra model with every carrying capacity set to one: the carrying capacities have been absorbed into the rescaled coefficients $\tilde{a}_{ij}$, which are the interaction strengths measured in the currency of each species' own self-limitation. This rescaling leaves the equilibrium and its stability untouched, and at equal carrying capacities the $\tilde{a}_{ij}$ are just the $a_{ij}$.

###### S3.6.2 What a relative yield total below one says about the interactions

Describe the community at any moment by two quantities: how much relative yield it holds in total, and how that total is divided among the species,

$$\mathrm{RYT} = \sum_i \mathrm{RY}_i\,, \qquad p_i = \frac{\mathrm{RY}_i}{\mathrm{RYT}}, \qquad \sum_i p_i = 1.$$

The vector $\boldsymbol{p}$ is the community's composition, written in relative yields. Substituting $\mathrm{RY}_i = \mathrm{RYT}\, p_i$ into Eq. S60, the total obeys a logistic law, where for the moment the growth rates are equal and $r_i \equiv r$,

$$\frac{1}{\mathrm{RYT}}\frac{d\,\mathrm{RYT}}{dt} = r\left(1 - \frac{\mathrm{RYT}}{\mathrm{RYT}^*(\boldsymbol{p})}\right), \qquad \frac{1}{\mathrm{RYT}^*(\boldsymbol{p})} = -\boldsymbol{p}^{\top}\tilde{\boldsymbol{A}}\,\boldsymbol{p}, \tag{S61}$$

At a fixed composition the whole community thus grows like a single logistic population, and the total it settles to is $\mathrm{RYT}^*(\boldsymbol{p})$—the curve drawn in Fig. 3A, B. Whether this settling point lies above or below one is decided by a single quadratic form. Expanding it with $\tilde{a}_{ii} = -1$ and subtracting $1 = (\sum_i p_i)^2$ gives

$$\frac{1}{\mathrm{RYT}^*(\boldsymbol{p})} - 1 = -\sum_{i<j} p_i \, p_j \, m_{ij}, \qquad m_{ij} = \left(\tilde{a}_{ij} + \tilde{a}_{ji}\right) + 2. \tag{S62}$$

The number $m_{ij}$ is positive when the pair competes within the pairwise interaction bound and negative when it competes beyond it.

Note that the matrix $\widetilde{\boldsymbol{A}} = \mathrm{diag}(\boldsymbol{K})^{-1}\boldsymbol{A}\,\mathrm{diag}(\boldsymbol{K})$ still carries $-1$ on its diagonal, and the relative yields solve $\widetilde{\boldsymbol{A}}\,\mathrm{RY} = -\mathbf{1}$: the rescaled system is again a gLV community, now with unit carrying capacities, and it has the same RYT, the same feasibility, and—the two Jacobians being similar—the same spectrum (Sec. S3.6.1). Theorem 1 therefore applies to $\widetilde{\boldsymbol{A}}$ verbatim: if $\tilde{a}_{ij} + \tilde{a}_{ji} > -2$ for every pair, that is if every margin $m_{ij}$ of Eq. S62 is positive, then RYT $> 1$.

Because the weights $p_i p_j$ are never negative, if every pair competes within the bound, $\mathrm{RYT}^*(\boldsymbol{p}) > 1$ at every composition: the community cannot underyield at any moment. If every pair competes beyond it, $\mathrm{RYT}^*(\boldsymbol{p}) \leq 1$ everywhere, and the total can never climb back through one: by Eq. S61 and its logistic form, it rises only while below $\mathrm{RYT}^*(\boldsymbol{p})$, so it cannot cross a ceiling that never exceeds one. Underyielding, once reached, is then permanent, and the community is on its way to losing a species.

Only when the margins are of mixed sign—some pairs within the pairwise interaction bound, some beyond it—can the total cross one as the composition shifts: this is the single exception, the compositional recovery of the blue curve in Fig. 3C, in which the species of the too competitive pairs are abundant at first and the community recovers as the milder competitors gain abundances. It is rare because it needs both kinds of pair present and a particular community composition dynamics.

###### S3.6.3 The total settles before the composition

For a measured RYT to report $\mathrm{RYT}^*(\boldsymbol{p})$, the total must reach its settling point faster than the composition drifts away. Under equal growth rates $r_i \equiv r$it does automatically. The RYT relaxes at that common rate $r$ whatever the interactions; the composition moves only as fast as the species differ in the competition they feel. The two rates can be read off exactly in the mean-field case: with $a_{ij} = -\rho$ and $\mathrm{RYT} = S/[1 + (S-1)\rho]$ (Eq. S50 with $\rho$ replaced by $-\rho$), the total relaxes with the common rate

$$\lambda_{\mathrm{total}} = -r, \qquad \lambda_{\mathrm{comp}} = -r\,\frac{\mathrm{RYT}}{S}\,(1-\rho), \qquad \frac{\lambda_{\mathrm{total}}}{\lambda_{\mathrm{comp}}} = \frac{1+(S-1)\rho}{1-\rho}, \tag{S63}$$

and the ratio, in absolute value, stays above $S - 1$ throughout the underyielding regime $\rho > 1$, decreasing as the competition grows. The separation is thus not an assumption but a property of the regime we care about: wherever a community underyields, its total is already at $\mathrm{RYT}^*(\boldsymbol{p})$, and a measured RYT $< 1$ is a reading of the interactions among the species currently present.

##### S3.6.4 Unequal growth rates

At equal growth rates the separation of time scales is automatic. We now let the $r_i$ differ and repeat the decomposition of Sec. S3.6.2. Write $\bar{r} = \sum_i r_i\, p_i$ for the growth rate averaged over the composition, and

$$q_i \;=\; \frac{r_i\, p_i}{\bar{r}}, \qquad \textstyle\sum_i q_i \,=\, 1, \tag{S64}$$

for the composition reweighted by growth rates: $q_i$ is the share of the community's turnover carried by species $i$. Substituting $\mathrm{RY}_i = \mathrm{RYT}\, p_i$ into Eq. S60 and summing, the total still obeys a logistic law, now with rate $\bar{r}$ and ceiling

$$\frac{1}{\mathrm{RYT}}\frac{d\,\mathrm{RYT}}{dt} \;=\; \bar{r}\left(1 - \frac{\mathrm{RYT}}{\mathrm{RYT}^*(\boldsymbol{p})}\right), \qquad \frac{1}{\mathrm{RYT}^*(\boldsymbol{p})} \;=\; -\boldsymbol{q}^\top \widetilde{\boldsymbol{A}}\, \boldsymbol{p}. \tag{S65}$$

The ceiling is the quadratic form of Eq. S61 with one of its two composition vectors replaced by $\boldsymbol{q}$: species that grow faster weigh more in setting the RYT the community settles to. At equal rates $\boldsymbol{q} = \boldsymbol{p}$ and Eq. S61 is recovered. The composition, in turn, obeys

$$\frac{1}{p_i}\frac{dp_i}{dt} \;=\; (r_i - \bar{r}) \;+\; \bar{r}\,\mathrm{RYT}\left(\frac{1}{\mathrm{RYT}^*(\boldsymbol{p})} - \frac{r_i}{\bar{r}}\,\Phi_i\right), \qquad \Phi_i \;=\; -\textstyle\sum_j \tilde{a}_{ij}\; p_j, \tag{S66}$$

where $\Phi_i$ is the competition species $i$ feels per unit of total, so that $\sum_i q_i\, \Phi_i = 1/\mathrm{RYT}^*(\boldsymbol{p})$. Two terms have appeared relative to the equal-rate case: a direct one, $r_i - \bar{r}$, by which a faster species gains share whatever the interactions, and the factor $r_i/\bar{r}$ by which a faster species responds more strongly to the competition it feels.

Whether the settling point lies above or below one is again decided by a sum over pairs. Expanding $-\boldsymbol{q}^\top\widetilde{\boldsymbol{A}}\boldsymbol{p}$ with $\tilde{a}_{ii} = -1$ and subtracting $1 = (\sum_i q_i)\big(\sum_j p_j\big)$ gives

$$\frac{1}{\mathrm{RYT}^*(\boldsymbol{p})} \;-\; 1 \;=\; -\,\frac{1}{\bar{r}}\textstyle\sum_{i<j} p_i\; p_j\; m_{ij}^{(r)}, \qquad m_{ij}^{(r)} \;=\; r_i\,\big(1 + \tilde{a}_{ij}\big) + r_j\,\big(1 + \tilde{a}_{ji}\big). \tag{S67}$$

This is Eq. S62 with each side of an interaction weighted by the growth rate of the species that feels it; at equal rates $m_{ij}^{(r)} = r\, m_{ij}$. The conclusions of Sec. S3.6.2 therefore carry over unchanged, with the rate-weighted margin in place of $m_{ij}$: if every $m_{ij}^{(r)}$ is positive the community cannot underyield at any composition, if every one is negative underyielding is permanent, and only mixed signs allow the total to cross one. The rate-weighted margin is also the mechanism behind the stable underyielding of Sec. S3.3.2: a pair may respect the pairwise interaction bound, $m_{ij} > 0$, and yet have $m_{ij}^{(r)} < 0$ when the species that suffers the stronger competition is also the faster one, or the reverse.

Reading a measured RYT as $\mathrm{RYT}^*(\boldsymbol{p})$ still requires the total to settle before the composition drifts. Setting $\mathrm{RYT} = \mathrm{RYT}^*(\boldsymbol{p})$ in Eq. S66, the composition drifts at the rate $r_i(1 - \mathrm{RYT}^*\Phi_i)$: it is slow when the species feel nearly the same competition, and it is the faster species that drift fastest, which is why the separation narrows, without breaking, as growth rates spread. Across simulations it holds even under strong heterogeneity in growth rates (Fig. S7).

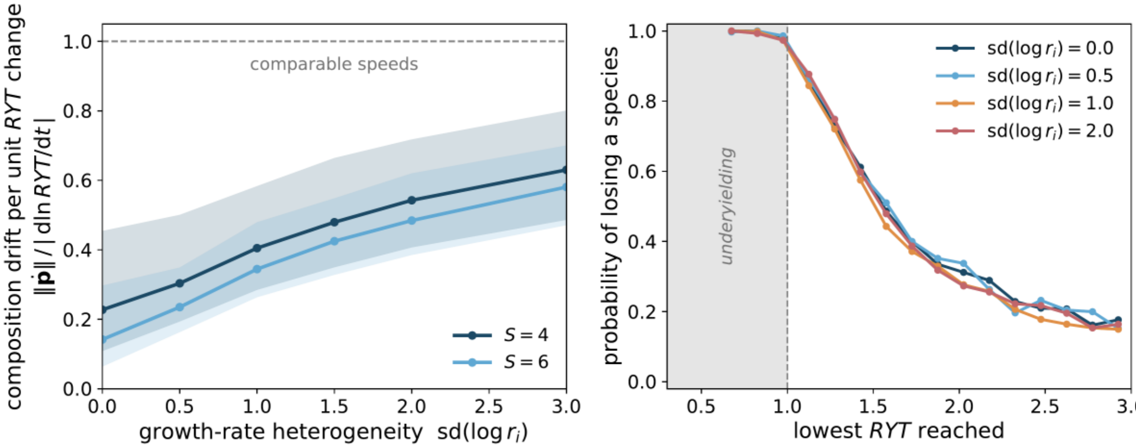


**Fig. S7.** The two-time-scale reading and the loss forecast are robust to heterogeneity in intrinsic growth rates. Communities are drawn independently at each level of growth-rate heterogeneity (sampling in Sec. S4.2); $sd(\log r_i) = 0$ is the equal-rate case. **(a)** Separation of time scales, measured as the composition's drift per unit change in the relative yield total, $\|d\boldsymbol{p}/dt\|/|d\ln \mathrm{RYT}/dt|$ averaged over the RYT's relaxation, against growth-rate heterogeneity (median with interquartile band, richness $S = 4$ and 6). The ratio rises with heterogeneity but stays below one (dashed): the RYT dynamic keeps its lead, with a narrowing margin. **(b)** Probability of losing at least one species against the lowest RYT a community reaches, one curve per level of heterogeneity. The curves nearly coincide; once a community underyields (shaded, RYT $< 1$), loss is all but certain—between 98.9% and 99.8% across all levels—regardless of the spread in growth rates.

##### S3.6.5 The loss of a species

That a community which underyields is losing a species can be seen directly. The reciprocal of Simpson's index $\sum_i p_i^2$ counts the effective number of species, and differentiating it along the composition dynamics gives, for any interactions,

$$\frac{d}{dt}\sum_i p_i^2 \;=\; -2\,r\,\mathrm{RYT}\;\mathrm{Cov}(p_i,\,\Phi_i), \tag{S68}$$

the covariance taken across species with each weighted by its share. Effective diversity falls whenever the abundant species are the ones under less than the average competition—which is what happens under the strong, mean-field competition that produces underyielding: there $\Phi_i = \rho + (1-\rho)p_i$ decreases with a species' share, so the abundant species feel the least competition and grow at the others' expense. Their ranking never reverses and every gap widens, so all the relative yield collects in the species that was most abundant to begin with, and the rest are lost. As they go, $\sum_i p_i^2 \to 1$ and $\mathrm{RYT}^* \to 1$ from below: the total climbs back toward one not because the community has recovered but because the species that caused the deficit are gone—the amber trajectory of Fig. 3C. We also explored this result numerically in Fig. S7 where the probability of losing a species stays close to 1 outside the mean-field case, whatever the heterogeneity in species growth rates.

#### S4 Numerical simulations

This section describes how the simulation figures (of both the manuscript and the SI) are generated. Throughout, interaction coefficients follow the convention of Eq. S1: $a_{ii} = -1$, with $a_{ij} < 0$ a competitive and $a_{ij} > 0$ a facilitative effect, so the pairwise bound reads $a_{ij} + a_{ji} > -2$. Figures S1, S2 and S6 are analytical or illustrative and involve no random sampling; the mean-field curves of Figs. 2B, S4 and S6 are evaluated numerically from Eq. S58.

##### S4.1 Equilibrium ensembles

For the gLV system of Eq. S1 we draw an interaction matrix $\boldsymbol{A}$ together with intrinsic growth rates and retain a community when it is feasible ($\boldsymbol{N}^* > 0$ with $\boldsymbol{K} > 0$). For each retained community we record the relative yield total $\mathrm{RYT} = \sum_i N_i^* / K_i$, the feasibility-domain size $\Omega$ (Eq. S14), the leading Jacobian eigenvalue $\lambda_{\max}$ (Eq. S3), and whether every pair satisfies the pairwise bound $a_{ij} + a_{ji} > -2$; the last splits each cloud into bound-satisfied and bound-violated subsets.

The mean off-diagonal coefficient $\mu$ is swept from facilitation to strong competition, so that each cloud spans both subsets, and $\sigma$ sets the spread around $\mu$. Four interaction structures are used:

- *i.i.d. Gaussian:* $a_{ij} \sim \mathcal{N}(\mu, \sigma^2)$, drawn independently; the baseline structureless prior.
- *Competitive hierarchy:* $a_{ij} = \mu + (h_i - h_j) + \xi_{ij}$, with $h_i$ a uniform ranking score and $\xi_{ij} \sim \mathcal{N}(0, \sigma_\xi^2)$.
- *r–K trade-off:* $a_{ij} = \mu + (c_i - c_j) + \zeta_{ij}$, with $c_i$ tied to the growth rate $r_i$ (fast-growing species compete less).
- *Niche overlap:* $a_{ij} = \mu \exp[-(x_i - x_j)^2 / 2\ell^2]$, with $x_i$ a one-dimensional trait and $\ell$ a niche width, so close pairs compete most.

In the ensembles of Figs. 2, S4 and S5, $\mu$ and $\sigma$ are drawn anew for each community, $\mu \sim \mathcal{U}(-3, 0.1/(S-1))$ and $\sigma \sim \mathcal{U}(0, 0.5)$; $\sigma$ also serves as the heterogeneity parameter of the structured ensembles ($\sigma_\xi = \sigma_\zeta = \ell = \sigma$), with $h_i, c_i \sim \mathcal{U}(-1/2, 1/2)$ and $x_i \sim \mathcal{U}(0,1)$. The equilibrium is drawn directly, $\log N_i^* \sim \mathcal{N}(0,1)$, and redrawn up to 1,000 times until $\boldsymbol{K} = -\boldsymbol{A}\boldsymbol{N}^* > 0$, the community being discarded otherwise; growth rates are half-normal, $r_i = |\xi_i|$ with $\xi_i \sim \mathcal{N}(0,1)$. $\Omega$ is estimated by quasi-Monte-Carlo integration (Sec. S1.4).

**Figure 1C, D.** Two-species systems with $K_1 = K_2 = 1$ and growth rates log-uniform on $[1/3, 3]$. The two interaction coefficients are drawn independently and uniformly, 3,500 pairs in each of three sets: competitive pairs both weaker than self-regulation, $a_{ij} \in (-0.985, -0.02)$ (bound satisfied, stable, RYT $> 1$), competitive pairs both stronger, $a_{ij} \in (-3, -1.015)$ (bound violated, unstable, RYT $< 1$), and pairs with at least one facilitative entry, both in $(-0.985, 0.5)$. The band $0.985 < |a_{ij}| < 1.015$ is deliberately left out: the equilibrium becomes singular there, and for a symmetric competitive pair $|a_{ij}| = 1$ is the bound $a_{ij} + a_{ji} = -2$ of Theorem 1. The first two sets appear as the stable and unstable points of panel D; facilitative entries ($a_{ij} > 0$) populate the range RYT $> S$, which pure competition cannot reach, and are the only source of the $\Omega > 1$ portion of the cloud. Panel C plots $\lambda_{\max}$ against RYT; panel D plots the two-species feasibility measure $\Omega$ against RYT. Panels A, B are fixed illustrative geometry, not sampled.

**Figure 2.** The i.i.d. Gaussian ensemble at $S = 10$. One panel plots $\lambda_{\max}$ against RYT over the full cloud, with the equal-rate analytical bound of Eq. S54 ($r = 1$, $S = 10$) overlaid; the other plots $\log_{10}\Omega$ against RYT, its trend taken over the bound-satisfied subset. 5,000 communities are drawn, of which 4,996 are feasible; panel B shows the 3,695 with a finite $\Omega$.

**Figures S4 and S5.** The same two relations across all sixteen combinations of species number $S \in \{3,5,10,15\}$ and the four structures; the first 2,000 feasible communities whose feasibility-domain size $\Omega$ is numerically positive are retained per combination. At $S = 15$ this filter discards most strongly competitive, underyielding communities, whose $\Omega$ falls below the precision of the numerical integration.

**Figure S3.** Feasible underyielding equilibria with unequal growth rates. The interactions are those of the dynamical ensemble of Fig. 3D (Sec. S4.2), $a_{ij} \sim \mathcal{N}(\mu, \sigma^2)$ with $\sigma = 0.48/\sqrt{S-1}$ and candidates spread equally over its sixteen values of $\mu$, so that this equilibrium analysis and the dynamical analyses of Figs. 3D and S7 rest on the same law of communities. The equilibrium is drawn directly, as in the ensembles of Figs. 2, S4 and S5, $\log N_i^* \sim \mathcal{N}(0,1)$, with $\boldsymbol{K} = -\boldsymbol{A}\boldsymbol{N}^*$, and a community is retained when $\boldsymbol{K} > 0$ and RYT $< 1$, which gives $\mathrm{sd}(\log K_i)$ from 0.45 ($S = 8$) to 0.66 ($S = 3$); 20,000 communities are retained per richness $S \in \{3,4,6,8\}$. Growth rates are $\log r_i \sim \mathcal{N}(0, s^2)$, one standard normal vector per community being scaled by $s$ so that all levels share their communities, and stability is read from the leading eigenvalue of $\boldsymbol{J}^*$ (Eq. S3).

##### S4.2 Dynamical ensembles

The same gLV system (Eq. S1 with $K_i = 1$, so each monoculture settles at $N_i = 1$ and $\mathrm{RYT}(t) = \sum_i N_i(t)$) is integrated forward from log-normal initial abundances, $\log N_i(0) \sim \mathcal{N}(0, 0.8^2)$. For each community we record the lowest RYT reached along the trajectory and whether at least one species is lost by the end of the integration, $N_i(T) \leq 10^{-3}$.

**Figure 3.** Panels A–C use fixed illustrative matrices (equal growth rates in A, B), each integrated from a chosen start; the four cases of panel C are described in the caption. Panel D is an ensemble at equal rates $r = 1$ with i.i.d. interactions $a_{ij} \sim \mathcal{N}(\mu, \sigma^2)$, $\sigma = 0.48/\sqrt{S-1}$ (so communities stay comparably far from the stability boundary as richness grows), and $\mu$ taking sixteen competitive values, seven evenly spaced from $-1.6$ to $-0.55$ and nine from $-0.42$ to $-0.03$, with 1,200 communities each; $S \in \{3,4,6,8\}$ (19,200 communities per richness). The loss probability is computed in classes of lowest RYT of width 0.2 from 0.3, shown when they hold more than 45 communities.

**Figure S7.** The same dynamical ensemble with unequal growth rates, $\log r_i \sim \mathcal{N}(0, s^2)$ rescaled to unit mean within each community ($s = 0$ recovers equal rates). Panel (a) plots the separation of time scales $\|d\boldsymbol{p}/dt\|/|\mathrm{dlnRYT}/\mathrm{d}t|$ against the heterogeneity $s$ (up to $s = 3$) at $S = 4$ and 6; panel (b) plots the probability of species loss against the lowest RYT reached, one curve per level $s \in \{0, 0.5, 1, 2\}$, with 19,200 four-species communities per level.

#### S5 Empirical assessment of the interaction bound

Here we quantify the empirical distribution of $a_{ij} + a_{ji}$ in the largest public compilation of pairwise plant–plant interaction coefficients[35], reporting the fraction of pairs that satisfy the pairwise interaction bound of our Theorem 1, both with all pairs and on the competition-only subset.

We used the Dryad release[54] of the Adler et al.[35] compilation, a meta-analytic compilation of intra- and interspecific interaction coefficients from 41 studies of terrestrial plant communities (39 of which enter the authors' analyses), reproducing the authors' data-cleaning procedure, with signs standardized so that negative values denote competition. Because studies use heterogeneous conventions and units, the raw coefficients $\alpha_{ij}^{(k)}$ are comparable only within a study, so we renormalize within each stratum (study × treatment × response × life stage),

$$a_{ij}^{(k)} = \frac{\alpha_{ij}^{(k)}}{|\alpha_{ii}^{(k)}|}. \quad \text{(S69)}$$

Equation S69 returns the dimensionless manuscript coefficient ($a_{ii} = -1$, with $a_{ij} < 0$ for competition and $a_{ij} > 0$ for facilitation) and is well-defined wherever intraspecific regulation is competitive ($\alpha_{ii}^{(k)} < 0$), a precondition of Theorem 1 in the main text. Within each stratum we form the unordered pairs $\{i, j\}$ whose four coefficients are jointly present with both intraspecific terms competitive, and exclude the 10 pairs in which one interspecific coefficient is exactly zero, whose sign cannot be classified—249 pairs from 22 studies—classified by their two interspecific signs into 136 both competitive (CC, 55%), 72 mixed (CF, 29%), and 41 both-facilitative (FF, 16%; Fig. S8B).

Across the 249 pairs the median of $a_{ij} + a_{ji}$ is $-0.31$ (Q1 $-1.14$, Q3 0.41), and 88% satisfy the bound (Fig. S8A); on the CC subset, where the bound is binding, the fraction is 80% (median $-0.90$) (Fig. S8B).

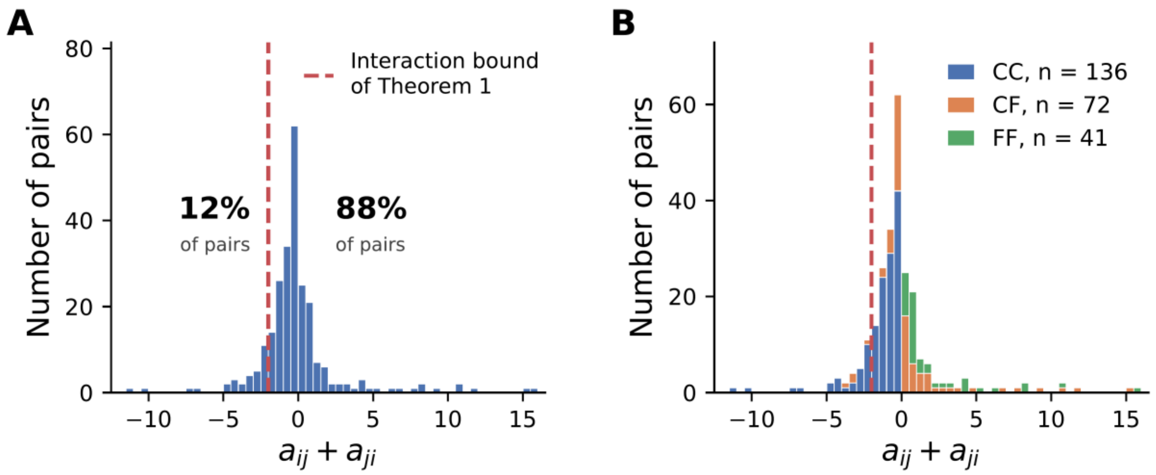


**Fig. S8. Empirical distribution of $a_{ij} + a_{ji}$ in the Adler et al.[35] compilation under the manuscript convention $a_{ii} = -1$. (A)** All 249 unordered pairs; the red dashed line is the interaction bound of Theorem 1, $a_{ij} + a_{ji} = -2$, satisfied by 88% of pairs. **(B)** Composition of the same distribution by pair type (CC: both directions competitive; CF: mixed; FF: both facilitative). FF pairs lie entirely on the positive side and satisfy the bound trivially.

### S6 Empirical datasets

This section describes the four datasets analyzed in the main text, their harmonization, and the construction of the unit–time panel on which the persistent-extinction outcome is defined; Table S1 summarizes the resulting panel.

#### S6.1 The four datasets

We retained four BEF panels—the Pennekamp et al. protist microcosms[26], Cedar Creek E120 (27, 28), the Dominance Experiment of the Jena Experiment[29,46], and the harvest-level Agrodiversity Experiment[30,31]—spanning three orders of magnitude in generation time and a range of controls from microcosms to multi-site agronomic systems. Two candidates were excluded: the full Jena Main Experiment, whose small-monoculture reference biomass falls threefold over the series (median 235 to 77 $\mathrm{g\,m^{-2}}$ between 2002 and 2009), and an annual aggregation of the Agrodiversity Experiment, which loses the within-year resolution at which RYT is naturally defined. Per-dataset composition is in Table S1; the harmonization specific to each follows.

***Pennekamp.*** Laboratory jars seeded with 1–6 ciliate species at six constant temperatures (15–25 °C, 115 microcosms each, 97 of them mixtures), sampled by automated video microscopy at 19 time points over 40 days[69]; two microcosms with no species detected at first sampling are excluded. Biomass is estimated from video and never inferred, and the monoculture reference for a species is its average biomass across replicate monocultures at matching temperature and day. Temperature enters the generalized estimating equation (GEE) as a fixed effect (Sec. S7).

***Cedar Creek E120.*** A perennial-prairie experiment in Minnesota, seeded in 1994 from 18 species. The two woody species[28] are excluded, and so are the plots sown with at least one of them (this removes 77 of the 172 plots). The 1994–1997 establishment phase is excluded, and so are 1998–2000, for which biomass was not sorted to species; the analysis window thus starts in 2001, with one late-summer clipping per plot and year. Monoculture references are computed per (species × year), as the mean over all contemporaneous monoculture plots of that species. Horizons are counted in calendar years; no plot was sorted to species in 2009, which is treated as unobserved (Eq. S70).

***Jena Dominance.*** The Dominance sub-experiment focuses on dominant species at moderate sown richness; its monoculture references come from the small-monoculture field (PANGAEA https://doi.org/10.1594/PANGAEA.866358; plot-to-species assignment from https://store.pangaea.de/Publications/Jena_Experiment/PlotInformationSmallMonos.txt), averaged per (species × year). Observations before 2004, the first two years after sowing in 2002, are excluded as an establishment phase.

***Agrodiversity.*** A multi-site agronomic experiment with four species (two grasses, two legumes), analyzed at 28 of its 33 sites (27 in Europe, one in Canada) and at the *harvest* resolution rather than annually, since one harvest is the finest unit at which RYT is naturally defined here; horizons are therefore counted in harvests. Monoculture references are computed per (site × year × harvest × species), and the reference community is the sown species present at $t_0$.

#### S6.2 Panel construction

For each unit, the reference community is the set of species present at the first *eligible observation* $t_0$—the first observation that is protocol-eligible (past the establishment-year exclusions above), has at least one species present, and has RYT computable (a monoculture reference for every species present). It fixes the extinction targets of the prospective test; species absent at $t_0$ are not tracked, deliberately excluding early-establishment failures, since the theorem concerns established communities.

A persistent-extinction event is recorded for a reference-community species when it is present at $t-1$, absent at $t$, and absent at $t+1$,

$$\text{event}(t) = \begin{cases} 1, & \text{if the species is present at } t-1 \text{ and absent at both } t \text{ and } t+1, \\ 0, & \text{otherwise.} \end{cases} \tag{S70}$$

under *strict adjacency*: $t-1$ and $t+1$ are the immediately preceding and following observations of the unit. The rule is undefined—and flagged NaN—at $t_0$ (no prior eligible observation, so establishment failures are not events), at each unit's last observation (persistence unverifiable), and at observations adjacent to a sampling gap (Cedar Creek 2009). Strict adjacency matches the exposure variable $E_{p,t}$ (Eq. 4), so both sides of the test use the same definition of persistence and a positive event count is not driven by year-spanning gaps.

The outcome $Y_{p,t_{\text{base}},h}$ is 1 if at least one reference-community species has an event (Eq. S70) within the prospective window $(t, t+h]$; events are read from biomass alone, so the window may extend beyond the RYT-computable period. A baseline is retained if at least half of its window steps are evaluable (as in the original Pennekamp analysis[26]), a step being evaluable when the event rule (Eq. S70) is defined there for at least one species. The resulting panel (mixtures only, $S_{\text{init}} > 1$) is summarized in Table S1; the binary and continuous GEE (Eqs. 5 and 6) use overlapping but distinct observations, since the binary exposure needs both the current and lag-1 RYT while the continuous predictor needs only the current, giving $n_{\text{cont}} \geq n_{\text{bin}}$.

**Table S1. Composition of the analysis panel for each dataset.** $n_{\text{plots}}$: number of mixture plots (microcosms for Pennekamp) entering the analysis after the mixtures-only filter $S_{\text{init}} > 1$. $n_{\text{bin}}^{(h)}$: number of (unit, baseline) observations entering the *binary* GEE at horizon $h$ (Eq. 5), which requires both the current and the lag-1 RYT to define the underyielding exposure $E_{p,t}$ (Eq. 4). $n_{\text{cont}}^{(h)}$: number of observations entering the *continuous* GEE at horizon $h$ (Eq. 6), which requires only the current RYT and is therefore at least as large as $n_{\text{bin}}^{(h)}$. Both counts may differ between horizons because the outcome window changes. Calendar range: first to last baseline. $S_{\text{init}}$ levels: sown-richness levels present in the mixture panel. RYT median: median of RYT across the panel. Frac. RYT $< 1$: empirical fraction of observations where the relative yield total is below 1.

| Dataset | Time unit | $n_{\text{plots}}$ | $n_{\text{bin}}^{(h=1)}$ | $n_{\text{bin}}^{(h=2)}$ | $n_{\text{cont}}^{(h=1)}$ | $n_{\text{cont}}^{(h=2)}$ | Calendar range | $S_{\text{init}}$ | RYT median | Frac. $< 1$ |
|---|---|---|---|---|---|---|---|---|---|---|
| Pennekamp | day | 580 | 9,280 | 9,280 | 9,280 | 9,280 | day 3–33 (40-d run) | {2,3,4,5,6} | 2.53 | 0.08 |
| Cedar Creek | year | 56 | 728 | 728 | 808 | 808 | 2002–2019 | {2,4,8,16} | 1.90 | 0.20 |
| Jena Dominance | year | 188 | 1,127 | 1,127 | 1,127 | 1,127 | 2005–2010 | {2,3,4,6,9} | 1.43 | 0.26 |
| Agrodiversity | harvest | 796 | 4,806 | 4,806 | 4,856 | 4,856 | 2003–2007 | {4} | 1.45 | 0.11 |

### S7 Statistical analysis

Here we present the regression analysis behind the odds-ratio estimates of the main text (Tables S2–S4): the generalized estimating equation (GEE) and its design choices, the caveats and the Firth and GEE estimators of the community odds ratios.

**Table of contents of S7**

S7.1 The regression model ........ S42
S7.2 Adjustment variables and binning ........ S42
S7.3 Statistical caveats ........ S43
S7.4 Robustness and power ........ S43
S7.5 Firth penalized estimation and the GEE cross-walk ........ S44

#### S7.1 The regression model

Inference uses a GEE with binomial family and logit link, clustered at the experimental unit (the microcosm for Pennekamp), with an exchangeable working correlation and cluster-robust Liang–Zeger sandwich standard errors[48]. The exchangeable structure assumes a common within-unit correlation and improves efficiency without affecting consistency of the point estimates; where it fails to converge or returns non-finite estimates—in eight of the twenty-four fits of this analysis —an independence working correlation is used as a fallback, under which the sandwich estimator remains the valid Liang–Zeger formula, so the fallback is a mild loss of efficiency, not a methodological discontinuity; every independence fit converges. The model is fit in two equally weighted forms: a binary exposure (underyielding over two consecutive observations) and a continuous predictor lnRYT, two tests of the same graded prediction. For the Pennekamp microcosms the linear predictor adds a six-level fixed effect for the experimental temperature gradient (15–25 °C), the study's principal designed driver of ciliate turnover[26], without which the RYT–extinction signal would be confounded with thermally driven turnover; this moderately attenuates the effect (the $h = 2$ GEE odds ratio moves from about 6.58 to 5.93) but the signal remains significant under either specification. The binary community odds ratios are estimated by Firth penalized logistic regression (Sec. S7.5) and the continuous coefficient $\beta_1'$ by the cluster-robust GEE; $P$-values are two-sided Wald, and the analysis code is in the archived repository.

#### S7.2 Adjustment variables and binning

The model adjusts for realized richness $S^{\mathrm{real}}$ at baseline (the number of species present at $t$) rather than sown richness $S_{\mathrm{init}}$: realized richness naturally controls for the effect of community size on a species' extinction risk, and reflects the community's actual state without absorbing the prospective exposure signal. Realized richness enters as a categorical effect $C(S^{\mathrm{real}}_{\mathrm{grouped}})$ rather than a linear term, because the richness–extinction relationship is not assumed linear and the sparse high-richness levels would induce separation. Levels are merged by the same rule in every dataset until every group holds at least two events and at least two non-events: scanning richness in increasing order, a deficient level is merged with its more populated neighbor, and the most populated group is the reference level. The merging is done separately per (dataset × horizon) cell and does not depend on the exposure, so it does not affect the test's nominal level. Requiring both events and non-events in every group guards against separation on the richness terms, while levels are merged only where that requirement is not met. The model also adjusts for $\Delta t_{p,t}$, the time since the reference observation $t_0$, as a linear term (Eqs. 5–6 of the main text).

##### S7.3 Statistical caveats

Three caveats attach to the community analysis. *Outcome compression:* the binary outcome collapses windows containing several extinctions into a single positive, discarding how many species are lost, most consequentially at the two datasets with the largest event counts (Jena Dominance and Agrodiversity). *Few clusters at Cedar Creek:* with 56 mixture plots, cluster-robust sandwich standard errors sit near the regime where they become anticonservative[70,71], so the Cedar Creek standard errors are read with caution. *Survivor bias:* species absent from the reference community at $t_0$ are not tracked—by design, since the theorem concerns established communities—and the early-establishment exclusions of Sec. S6 keep the analysis window well after the colonization phase.

##### S7.4 Robustness and power

**Three-observation horizon.** The primary horizon is $h = 2$, and $h = 1$ is reported in Tables S3 and S4; we refit the same specification at $h = 3$ for all four datasets (Table S2). The qualitative pattern is preserved: all four odds ratios stay in the predicted direction and the pooled estimates remain in the predicted direction and significant for the odds ratio (2.31, 95% CI [1.01,5.30], $P = 0.049$) and borderline for the continuous coefficient ($\beta_1' = -0.38$ $[-0.77, +0.01]$, $P = 0.052$). Under the Firth estimator, three of the four datasets are significant, the wide unpenalized intervals of the rarest cells collapsing once the separation bias is removed, and a mature post-transient grassland like Cedar Creek manifesting the signal once the window is long enough to capture the slow turnover of perennial communities. The lone non-significant cell is Jena Dominance, whose binary estimate softens while its continuous $\beta_1'$ stays in the protective direction ($P = 0.096$).

**Table S2. Three-observation horizon sensitivity.** Odds ratios estimated by Firth penalized logistic regression and the continuous coefficient $\beta_1'$ by cluster-robust GEE, as in the main analysis, at horizon $h = 3$. Horizons are counted in sampling dates for Pennekamp (every two days), in harvests for Agrodiversity and in calendar years for Cedar Creek and Jena Dominance. The *Pooled (RE)* row is a random-effects estimate (Sec. S7.5); its heterogeneity is $I^2 = 65\%$ for the odds ratio and 53% for $\beta_1'$. Bold indicates $P < 0.05$; significance codes $^{***}P < 0.001$, $^{**}P < 0.01$, $^{*}P < 0.05$, $\cdot$ $P < 0.10$.

| Dataset | $n_{\text{bin}}$ | Events | Exposed | OR (95% CI) | $P_{\text{OR}}$ | $\beta_1'$ ($P_{\beta_1'}$) |
|---|---|---|---|---|---|---|
| Pennekamp | 8,700 | 100 | 351 | $\mathbf{4.66}$ $[\mathbf{2.09}, \mathbf{10.37}]$ | $< \mathbf{0.001}^{***}$ | $\mathbf{-0.59}$ $(\mathbf{0.016}^{*})$ |
| Cedar Creek | 630 | 81 | 88 | $\mathbf{3.34}$ $[\mathbf{1.64},\ \mathbf{6.8}$ | $< \mathbf{0.001}^{***}$ | $-0.19$ $(0.342)$ |
| Jena Dominance | 1,127 | 356 | 140 | $1.42$ $[0.91,\ 2.21]$ | $0.125$ | $-0.20$ $(0.096\cdot)$ |
| Agrodiversity | 4,010 | 962 | 122 | $\mathbf{1.88}$ $[\mathbf{1.26},\ \mathbf{2.8}$ | $\mathbf{0.002}^{**}$ | $\mathbf{-0.60}$ $(< \mathbf{0.001}^{***})$ |
| *Pooled (RE)* | — | — | — | $\mathbf{2.31}$ $[\mathbf{1.01},\ \mathbf{5.3}$ | $\mathbf{0.049}^{*}$ | $-0.38$ $(0.052\cdot)$ |

**Working correlation.** The fallback of Sec. S7.1 is not confined to the horizon extension: it applies to Cedar Creek and Agrodiversity at $h = 2$ as well as to Cedar Creek and Jena Dominance at $h = 3$. Since the independence sandwich remains the valid Liang–Zeger estimator, the consequence is a loss of efficiency in those cells, not a change of model.

**Persistence filter.** The canonical exposure requires underyielding over two consecutive observations ($E_2$); we compared it with the instantaneous ($E_1$) and three-observation ($E_3$) exposures (Table S3). The effect of the stricter filter depends on the horizon. At $h = 1$, a third underyielding observation raises the odds ratio at Pennekamp (4.93 → 10.01), Agrodiversity (2.33 → 3.61) and Jena Dominance (1.87 → 2.89) and leaves Cedar Creek flat; at $h = 2$ it barely moves except at Cedar Creek (2.61 → 3.29); at $h = 3$ it falls in three of the four datasets. Stricter persistence isolates genuinely underyielding units, as the theorem predicts, but $E_3$ typically halves the number of exposed baselines and so costs power. The choice of $E_2$ balances the two effects.

**Table S3. Sensitivity of the OR estimate to the persistence filter on the exposure variable.** $E_1$: instantaneous underyielding (RYT $< 1$ at $t$ only). $E_2$: Underyielding over two strictly adjacent observations (the canonical exposure). $E_3$: Underyielding over three strictly adjacent observations. All cells use Firth penalized logistic regression with the specification of Sec. S7 (as in the main analysis), varying only the exposure variable. Its $E_2$ cells at $h = 1$ and $h = 2$ coincide with the Firth odds ratios of Table S4. Bold: $P < 0.05$. The primary specification is $E_2$ at $h = 2$.

| Dataset | Horizon | $E_1$ OR ($P$) | $E_2$ OR ($P$) | $E_3$ OR ($P$) |
|---|---|---|---|---|
| Pennekamp | 1 obs. | 2.81 (0.057)$^{\cdot}$ | **4.93 (0.009)**$^{**}$ | **10.01 (< 0.001)**$^{***}$ |
| Pennekamp | 2 obs. | **3.36 (0.002)**$^{**}$ | **5.95 (< 0.001)**$^{***}$ | **5.88 (< 0.001)**$^{***}$ |
| Pennekamp | 3 obs. | **4.58 (< 0.001)**$^{***}$ | **4.66 (< 0.001)**$^{***}$ | **4.75 (0.001)**$^{**}$ |
| Cedar Creek | 1 yr | 1.35 (0.538) | 1.90 (0.218) | 1.91 (0.315) |
| Cedar Creek | 2 yr | **2.25 (0.021)**$^{*}$ | **2.61 (0.020)**$^{*}$ | **3.29 (0.019)**$^{*}$ |
| Cedar Creek | 3 yr | **2.64 (0.002)**$^{**}$ | **3.34 (< 0.001)**$^{***}$ | **3.01 (0.018)**$^{*}$ |
| Jena Dominance | 1 yr | 0.88 (0.674) | 1.87 (0.072)$^{\cdot}$ | **2.89 (0.014)**$^{*}$ |
| Jena Dominance | 2 yr | 1.21 (0.354) | **2.09 (0.004)**$^{**}$ | **2.11 (0.029)**$^{*}$ |
| Jena Dominance | 3 yr | 1.29 (0.151) | 1.42 (0.125) | 1.27 (0.463) |
| Agrodiversity | 1 harv. | **2.31 (< 0.001)**$^{***}$ | **2.33 (< 0.001)**$^{***}$ | **3.61 (< 0.001)**$^{***}$ |
| Agrodiversity | 2 harv. | **2.23 (< 0.001)**$^{***}$ | **2.32 (< 0.001)**$^{***}$ | **2.32 (0.011)**$^{*}$ |
| Agrodiversity | 3 harv. | **2.56 (< 0.001)**$^{***}$ | **1.88 (0.002)**$^{**}$ | 1.44 (0.319) |

##### S7.5 Firth penalized estimation and the GEE cross-walk

The binary community odds ratios (Fig. 4A; Table S4) are estimated by Firth penalized logistic regression. Ordinary logistic maximum likelihood is biased away from the null when events are rare or an exposure cell is nearly empty, and its Wald intervals can become uninformative—unbounded in the limit of quasi-complete separation. Firth's method[47] removes the leading-order bias by maximizing the penalized log-likelihood

$$\ell^*(\boldsymbol{\beta}) = \ell(\boldsymbol{\beta}) + 1/2 \log\det\mathcal{I}(\boldsymbol{\beta}), \quad \text{(S71)}$$

where $\mathcal{I}(\boldsymbol{\beta})$ is the Fisher information; the penalty is Jeffreys' invariant prior and is equivalent to adding about half an event and half a non-event to each cell, keeping both the point estimate and its interval finite and approximately unbiased. The penalty matters most where events are scarce: relative to the unpenalized cluster-robust GEE estimate, Firth shrinks the Jena Dominance $h = 2$ odds ratio from $2.47$ to $2.09$, leaves the Agrodiversity $h = 2$ odds ratio essentially unchanged ($2.31$ under GEE, $2.32$ under Firth), and tightens the near-degenerate Pennekamp $h = 2$ interval from [1.27,27.67] to [2.50,14.15].

Because the penalized-Wald interval is not cluster-robust, Table S4 reports for every cell the cluster-robust GEE estimate of the same binary odds ratio—the exponentiated $\hat{\beta}_1$ of Eq. 5 with Liang–Zeger intervals—beside the Firth estimate. The two estimators agree on the substantive conclusion: Firth shrinks some point estimates toward the null (most visibly Jena Dominance $h = 2$, $2.47 \to 2.09$) and stabilizes the Pennekamp intervals, but the direction and approximate magnitude are preserved in every cell, and the two columns classify the same cells as significant in seven of the eight. The exception, Cedar Creek $h = 2$, is the most fragile cell: its Firth interval excludes one, but neither its GEE interval [0.97,6.86] nor its cluster-bootstrap interval [0.69,7.03] does, so we do not treat its per-dataset significance as firm. The inferential target for the community claim is instead the pooled random-effects estimate, which is significant and robust to the choice of estimator two observations ahead (GEE $\mathrm{OR} = 2.52$, $P = 0.014$; Firth $\mathrm{OR} = 2.61$, $P = 0.016$) and one observation ahead (GEE $\mathrm{OR} = 2.23$, $P = 0.024$; Firth $\mathrm{OR} = 2.29$, $P = 0.019$).

**Table S4. Cross-walk between the cluster-robust GEE and Firth penalized estimators of the community binary odds ratio.** Each row gives the odds ratio for subsequent species loss following underyielding ($\mathrm{RYT} < 1$ over two consecutive observations), estimated by GEE (exponentiated $\hat{\beta}_1$ of Eq. 5, cluster-robust Liang–Zeger interval) and by Firth penalized logistic regression (penalized-Wald interval; Sec. S7.5). *Pooled (RE)* rows are random-effects estimates (Sec. S7.5); their heterogeneity is $I^2 = 0\%$ (GEE, $h = 1$), 0% (GEE, $h = 2$), 0% (Firth, $h = 1$), 33% (Firth, $h = 2$), and their two-sided $P$ values are $0.024$ and $0.014$ (GEE) and $0.019$ and $0.016$ (Firth). Bold indicates $P < 0.05$ for that estimator. †Cedar Creek $h = 2$: significant under Firth ($P = 0.020$) but not under GEE ($P = 0.058$), and its cluster-bootstrap interval [0.69,7.03] includes one, so its per-dataset significance is not treated as firm (Sec. S7.5).

| Dataset | Horizon | GEE OR (95% CI) | Firth OR (95% CI) |
|---|---|---|---|
| Pennekamp | 1 | **4.51 [1.01, 20.20]** | **4.93 [1.48, 16.36]** |
| Pennekamp | 2 | **5.93 [1.27, 27.67]** | **5.95 [2.50, 14.15]** |
| Cedar Creek | 1 | 1.63 [0.41, 6.47] | 1.90 [0.68, 5.28] |
| Cedar Creek† | 2 | 2.58 [0.97, 6.86] | **2.61 [1.17, 5.83]** |
| Jena Dominance | 1 | 1.87 [0.99, 3.53] | 1.87 [0.94, 3.72] |
| Jena Dominance | 2 | **2.47 [1.42, 4.31]** | **2.09 [1.27, 3.45]** |
| Agrodiversity | 1 | **2.40 [1.43, 4.02]** | **2.33 [1.45, 3.77]** |
| Agrodiversity | 2 | **2.31 [1.37, 3.90]** | **2.32 [1.56, 3.45]** |
| *Pooled (RE)* | 1 | **2.23 [1.22, 4.07]** | **2.29 [1.30, 4.05]** |
| *Pooled (RE)* | 2 | **2.52 [1.44, 4.43]** | **2.61 [1.40, 4.87]** |

**Random-effects pooling.** Each headline effect is summarized across the four experiments by random-effects meta-analysis rather than by pooling raw observations, since the experiments differ in generation time, design, and species pool. Writing $\hat{\theta}_d$ for the dataset-specific estimate with within-experiment variance $v_d$, the between-experiment heterogeneity $\tau^2$ is the DerSimonian–

Laird moment estimator[49] and the experiments are combined with weights $(v_d + \tau^2)^{-1}$; because only $k = 4$ experiments are pooled, the asymptotic interval is anticonservative, so we apply the Hartung–Knapp–Sidik–Jonkman adjustment[50,51] (referring the result to a $t$ distribution on $k - 1 = 3$ degrees of freedom and taking the more conservative of the model-based and adjusted standard errors). Heterogeneity is reported as $I^2$, the share of total variation attributable to between-experiment differences. The same procedure produces the *Pooled (RE)* estimates of the community odds ratio (Fig. 4A) and of the continuous coefficient (Fig. 4B).

### References

References 1–55 are listed in the main text; references 56–71 are listed at the end of this document.


56. Loreau, M., Sapijanskas, J., Isbell, F. & Hector, A. Niche and fitness differences relate the maintenance of diversity to ecosystem function: comment. *Ecology* **93**, 1482–1487 (2012), doi:10.1890/11-0792.1.
57. Desallais, M., Arnoldi, J.-F. & Loreau, M. Partitioning net biodiversity effects on ecosystem resistance and resilience. *Ecology Letters* **28**, e70249 (2025), doi:10.1111/ele.70249.
58. Genz, A. & Bretz, F. *Computation of Multivariate Normal and t Probabilities*, vol. 195 of *Lecture Notes in Statistics* (Springer-Verlag, Heidelberg, 2009), doi:10.1007/978-3-642-01689-9.
59. Genz, A. *et al. mvtnorm: Multivariate Normal and t Distributions* (2016), https://CRAN.R-project.org/package=mvtnorm, R package version 1.0-5.
60. Hofbauer, J. & Sigmund, K. *Evolutionary Games and Population Dynamics* (Cambridge University Press, 1998).
61. Engel, A. *Problem-Solving Strategies* (Springer, New York, 1998), doi:10.1007/b97682.
62. May, R. M. Will a large complex system be stable? *Nature* **238**, 413–414 (1972), doi:10.1038/238413a0.
63. Logofet, D. O. *Matrices and Graphs: Stability Problems in Mathematical Ecology* (CRC Press, Boca Raton, FL, 1993).
64. Horn, R. A. & Johnson, C. R. *Matrix Analysis* 2nd edn (Cambridge University Press, 2012).
65. Allesina, S. & Tang, S. Stability criteria for complex ecosystems. *Nature* **483**, 205–208 (2012), doi:10.1038/nature10832.
66. Coyte, K. Z., Schluter, J. & Foster, K. R. The ecology of the microbiome: networks, competition, and stability. *Science* **350**, 663–666 (2015), doi:10.1126/science.aad2602.
67. Mougi, A. & Kondoh, M. Diversity of interaction types and ecological community stability. *Science* **337**, 349–351 (2012), doi:10.1126/science.1220529.
68. Steck, G. P. Orthant probabilities for the equicorrelated multivariate normal distribution. *Biometrika* **49**, 433–445 (1962), doi:10.1093/biomet/49.3-4.433.
69. Altermatt, F. *et al.* Big answers from small worlds: a user's guide for protist microcosms as a model system in ecology and evolution. *Methods in Ecology and Evolution* **6**, 218–231 (2015), doi:10.1111/2041-210X.12312.

70. Mancl, L. A. & DeRouen, T. A. A covariance estimator for GEE with improved small-sample properties. *Biometrics* **57**, 126–134 (2001), doi:10.1111/j.0006-341X.2001.00126.x.

71. Hin, L.-Y. & Wang, Y.-G. Working-correlation-structure identification in generalized estimating equations. *Statistics in Medicine* **28**, 642–658 (2009), doi:10.1002/sim.3489.
